# Transcriptome-Wide Gene-Gene Interaction Association Study Elucidates Pathways and Functional Enrichment of Complex Traits

**DOI:** 10.1101/2022.08.16.504187

**Authors:** Luke M. Evans, Christopher H. Arehart, Andrew D. Grotzinger, Travis J. Mize, Maizy S. Brasher, Jerry A. Stitzel, Marissa A. Ehringer, Charles A. Hoeffer

## Abstract

It remains unknown to what extent gene-gene interactions contribute to complex traits. Here, we introduce a new approach using predicted gene expression to perform exhaustive transcriptome-wide interaction studies (TWISs) for multiple traits across all pairs of genes expressed in several tissue types. Using imputed transcriptomes, we simultaneously reduce the computational challenge and improve interpretability and statistical power. We discover and replicate several interaction associations, and find several hub genes with numerous interactions. We also demonstrate that TWIS can identify novel associated genes because genes with many or strong interactions have smaller single-locus model effect sizes. Finally, we develop a method to test gene set enrichment of TWIS associations (E-TWIS), finding numerous pathways and networks enriched in interaction associations. Epistasis is likely widespread, and our procedure represents a tractable framework for beginning to explore gene interactions and identify novel genomic targets.

---

Genome-wide association studies (GWASs) have identified numerous individual loci that affect complex traits^1,2^. Recent developments in transcriptome imputation and transcriptome-wide association studies (TWASs) have enhanced our understanding of complex traits by providing biologically plausible mechanisms of action for associated genes and improving power by aggregating small individual variant effects on gene expression to identify associations^3-5^. The overwhelming majority of these identified loci have been detected using an additive model of alleles at individual loci^1,2,6^.

While GWAS and TWAS have expanded our understanding of the genetic architecture underlying complex traits, a fundamental, unresolved question is to what extent non-additive effects contribute. Specifically, epistasis, defined as the statistical dependence of the allelic effects at one locus on the genotype at another locus^7^, may influence quantitative traits^7-10^. It is increasingly clear that complex traits are exceedingly polygenic, with influences from many complex regulatory and molecular pathways, and even chromosomal three-dimensional structure^11-13^. While there has been debate over whether non-additive genetic variance is a major contributor to heritability^6,14-18^, such complexity makes gene interactions likely to exist and these interactions have been demonstrated using several systems and model organisms^7-9,19,20^. Identifying gene-gene interactions and the pathways and networks in which they occur will provide a critical context for understanding the biology of complex traits^7,10^. Ascertaining the prevalence and magnitude of epistasis would also clarify interpretation of family-based, and specifically twin-based, estimates of heritability, which may be inflated by non-additive variance in combination with maternal or environmental effects^14,18^.

Despite the likely importance of epistasis, genome-wide interaction tests remain rare. Computational burden, correlation among predictors (leading to false positive epistatic associations^21,22^), and interpretability are key challenges to genome-wide, exhaustive tests of epistasis^7,23-25^. Perhaps the greatest challenge is that the sheer number of variants available in imputation panels (10M+) leads to tens of billions of pairwise tests, which despite recent methodological advances^26,27^ remains prohibitive. Many address this through two-stage approaches, in which the predictors are filtered in some way prior to testing epistasis among the retained predictors^24,28^. Often, interactions are only tested between loci that are significant in single-locus GWAS or phenotypic variance test effects, or are based on hypothesized pathways or networks. While such methods improve feasibility by reducing the number of tests, they constrain the ability to detect novel epistatic effects or new pathways and networks involved in complex traits^8^, and in some cases do not indicate whether the interactor effect is an environment or a second gene^28,29^. Similarly, if a strong interaction between two loci exists, the main effects estimated in a single-variant GWAS could be muted^7^, reducing the likelihood of identifying such interactions in two-stage approaches. Thus, exhaustive approaches are preferable to two-stage or filtered approaches.

Here, we report a new approach using imputed transcriptomes to perform exhaustive transcriptome-wide interaction studies (TWISs) for multiple traits across all pairs of genes expressed in several tissue types (Fig. 1). Using imputed transcriptomes, we provide an approach to simultaneously reduce the computational challenge and improve interpretability, while also aggregating small interaction effects of individual variants via gene expression to improve statistical power to detect interaction associations. We begin by performing extensive simulations to validate the TWIS approach and develop standardized analytic procedures, including power analyses, multiple test correction thresholds, and pruning on linkage that can lead to false positives. Importantly, we find that unmodeled interactions can also produce false negatives for main effects such that TWIS both identifies epistatic effects and identifies previously unassociated loci. Finally, we develop and validate Enrichment TWIS (E-TWIS), a novel method for aggregating genome-wide gene-gene interactions with respect to *a priori*-defined gene sets to understand the specific functional networks enriched for epistatic effects. In an empirical application, we identify several replicated, significant interactions and numerous functional gene sets and brain cell types that are enriched in interaction associations. Epistasis is likely a major source of phenotypic variation in complex traits, and the analytic procedures and results presented here reflect a computationally and statistically tractable framework for beginning to unpack these interactive effects.

**Figure 1.**
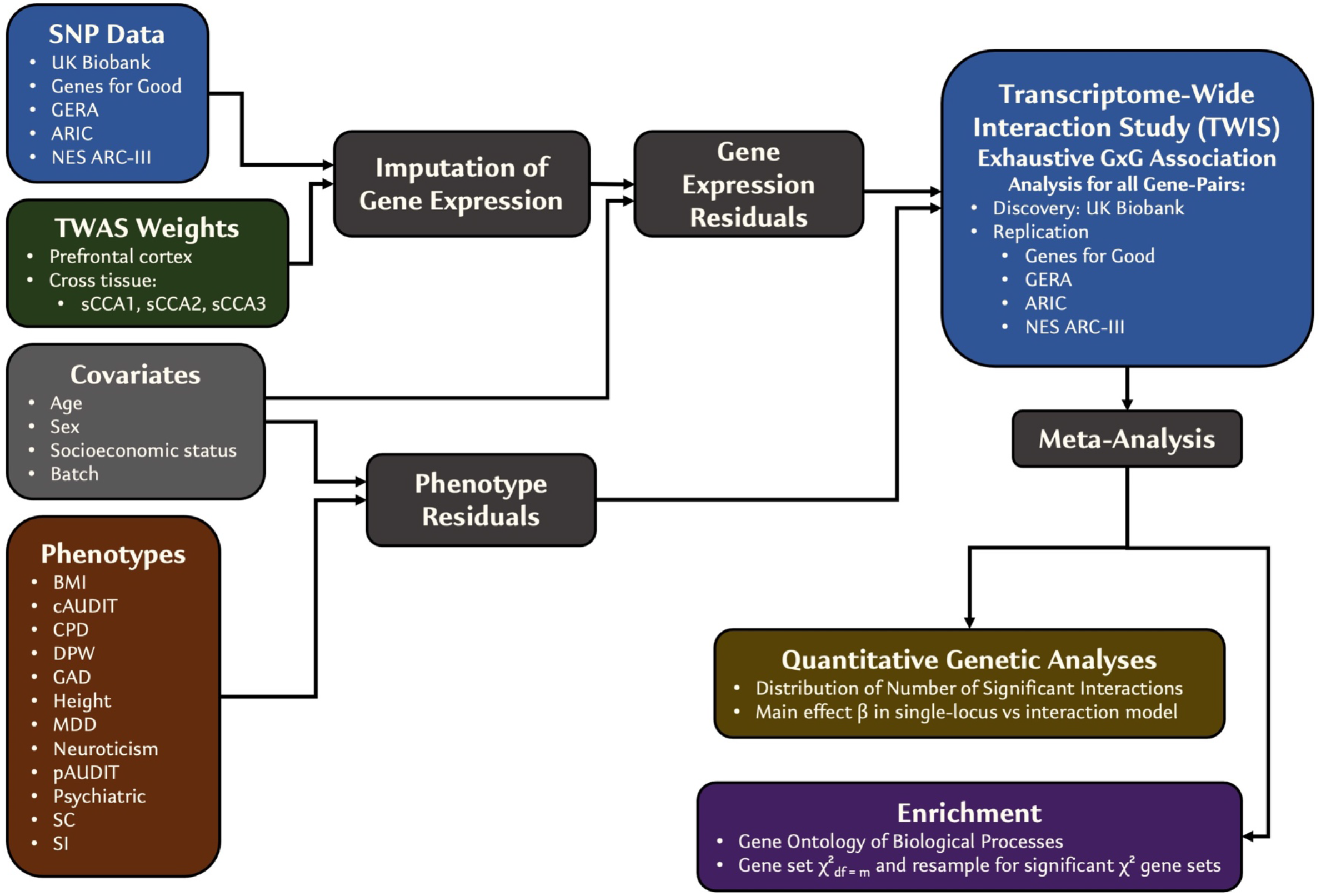
Overview of TWIS approach.

## Results

### TWIS Approach – Simulation and Validation

Figure 1 is a diagram of our overall transcriptome-wide interaction study (TWIS) approach. We leveraged a total of five cohorts to perform discovery and replication TWIS of 12 complex traits, including biometric, substance use, and psychiatric traits (defined in Supplementary Table S1 and Supplementary Information). We used the UK Biobank as the discovery cohort to identify significant interactions (N=53,880-329,705) and used the remaining 2-3 cohorts (depending on the trait) as an independent replication sample (N=8,718-61,531). Following standard quality control (see supplemental methods), we imputed gene expression in each cohort for the prefrontal cortex (PFC, *m*=14,729 genes) using FUSION^3,4^-generated TWAS weights from the PsychENCODE consortium^30^. The PFC was chosen because of the importance of neurocognitive functions in many of the traits we examined (e.g., psychiatric and substance use traits) and because it is currently the largest available brain reference panel with expression TWAS weights. Because of the large number of possible tissues relevant to complex traits, we also used cross-tissue expression weights from the first three sparse canonical correlation axes (sCCA1-3) of Feng et al.^31^ (*m*=13,242; 12,521; and 12,032).

Correctly accounting for covariates and possible confounding effects in interaction associations requires including all covariate-by-main effect interactions^32^, which quickly increases computational time with numerous covariates and categorical factor levels. Therefore, following QC and expression imputation, we residualized phenotype and imputed expression on covariates prior to performing the gene-gene interaction associations (see methods). The cohorts differed in the specific measures available, but included measures of age, sex, educational attainment, income or socioeconomic status, genotyping batch (where available), and the first 10 genomic principal components. When performing 10s of millions of tests, this residualization step substantially decreased the total computation time while estimating unbiased gene-gene interaction effects. Following this step, we used a parallelization procedure to divide all 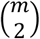 pairwise interactions across multiple compute nodes for each trait and each tissue, testing the simplified model,

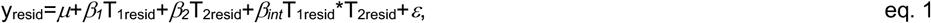

where y_resid_ is the phenotype residualized on the covariates; T_1resid_ and T_2resid_ are the imputed expression of genes 1 and 2, respectively, residualized on the covariates; *µ* is the intercept; *β*_*1*_ and *β*_*2*_ are the main effects of T_1resid_ and T_2resid_; *β*_*int*_ is the gene expression interaction effect on the phenotype; and *ε*∼N(0,1) is the error. We emphasize that this model does not require physical interaction of gene products, only that the association of expression of one gene is affected by that of another. Such interactions could include physical interaction, but also other mechanisms, such as stoichiometric relationships within molecular pathways.

*Power and Significance Thresholds*. We performed a series of simulations to estimate power to detect interactions in the context of imperfect expression imputation across a range of epistasis effect sizes, define the appropriate *σ* for genome-wide multiple test correction in the context of many millions of individual tests, and assess the role of linkage in influencing interaction tests (see Methods and Supplemental Figures S1-S8). Consistent with prior findings^21,22^, we find that pairs of genes with imputed expression correlations (|*r*|) > 0.1 or those physically nearby produce inflated type I error for identifying interaction effects.

Within each phenotype, we applied a significance threshold of *p*<5.86e-10 (see Methods) while also excluding from further analysis any pairs of genes whose imputed expression |*r*|>0.05 at the discovery stage (UK Biobank sample) or those within 1MB of each other. In independent replication, we applied, first, this correction within each phenotype and tissue to interactions identified within the discovery cohort, and second, a nominal p<0.05 as suggestive evidence of replication. Finally, we meta-analyzed^33^ all cohorts together (discovery + replication) for use in functional and pathway enrichment analyses.

### TWIS Associations – Empirical Results

Across 12 traits (height, BMI, cigarette smoking initiation [SI], smoking cessation [SC], heavy vs. light cigarettes per day [CPD], major depressive disorder [MDD], generalized anxiety disorder [GAD], neuroticism, cross-trait psychiatric disorders [PSYCH], problematic alcohol use [pAUDIT], alcohol consumption [cAUDIT], and drinks per week [DPW]; see Supplementary Materials for full phenotype and cohort descriptions), 16 pairwise interactions were significant (*p*<5.86e-10) at the discovery stage, with one replicating in independent replication datasets in the same direction and four remaining significant in the final (discovery + replication) meta-analysis (Fig. 2, Table 1, Supplementary Figures S9-S20). An additional two pairwise interactions were significant when all cohorts were meta-analyzed. Of the five significant in the final meta-analysis, three associations with pAUDIT involved *PRKCG* imputed prefrontal cortex expression, interacting with *WNT6, MAP7*, and *SEZ6L2*. Notably, WNT is known to modulate PKC localization and activity via G-protein- and Ca2+-dependent mechanisms^34,35^. MAP7 is known to directly interact with PKC signaling^36^ and has a role in axon collateral branching^37,38^. SEZ6L2 is a cell-surface protein that regulates neurogenesis and differentiation through adducin signal transduction^39^, which is a substrate for PKC.^40^ The fourth interaction associated with pAUDIT was *TFCP2L1*x*CENPN. TFCP2L1*, which is down regulated in cells exposed to alcohol^41^, regulates transcription involved in pluripotency and cell renewal and is also involved in the WNT pathway^42^. CENPN is a histone that forms a complex with other histones in the presence of DNA and locates at the centromere, forming kinetochores^43^; their interaction may reflect effects on neurogenesis or neural cell types from a brain stem cell.

**Figure 2.**
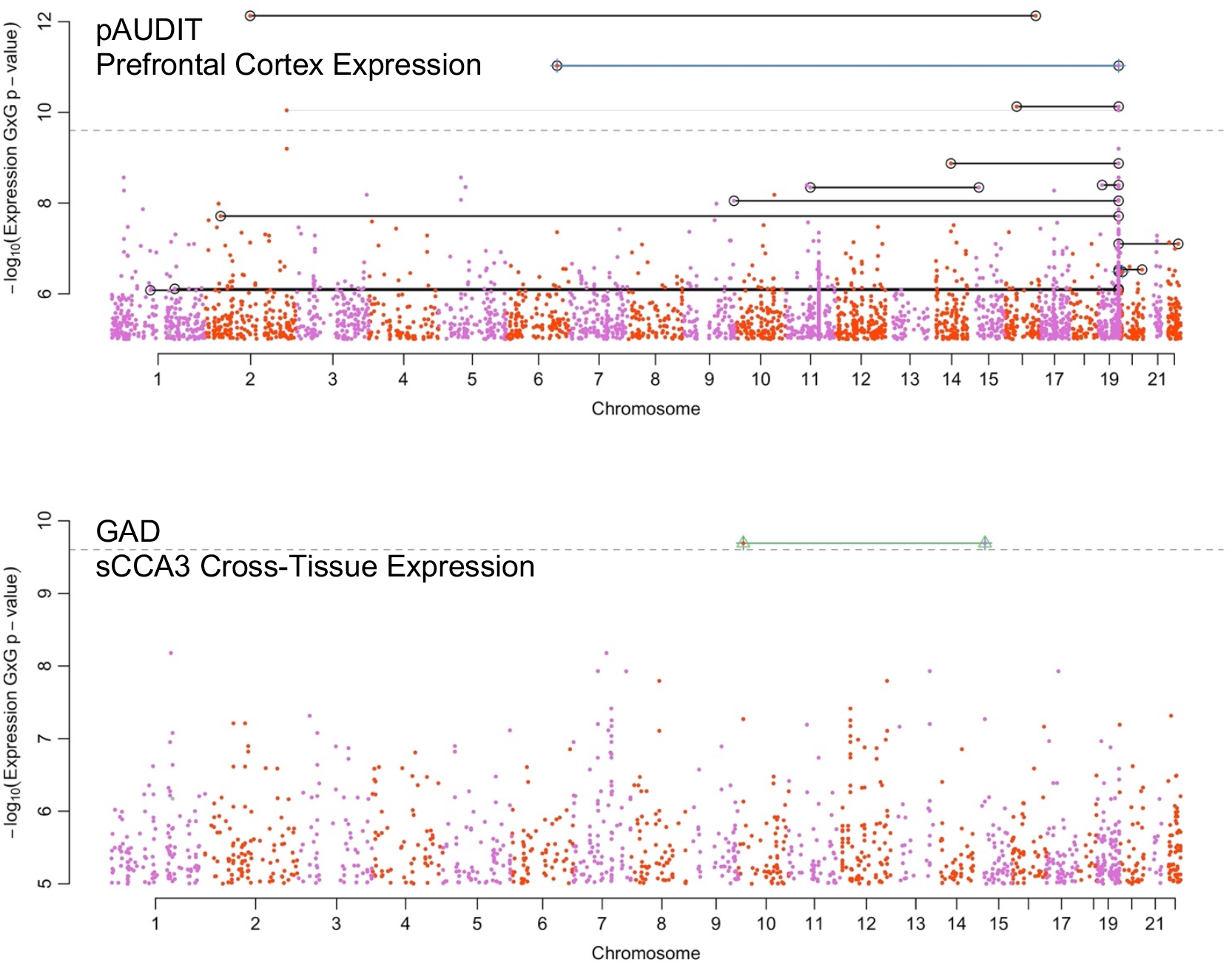
Boulder plot of pAUDIT (top) and GAD (bottom) interaction association p-values using imputed transcription. Shown are the results from the final meta-analysis of all data. Black lines connect pairs that surpassed p<5.86e-10 in the discovery cohort (UKB), green and blue lines connect pairs of loci with FDR q<0.05 or nominally significant interaction (p<0.05) in the replication cohort, and gray lines connect pairs of genes with p<2.5e-10 in the final meta-analysis. For clarity, only interaction associations with p<1e-5 are shown. Numerical results of genes reaching significance are presented in Table 1.

**Table 1.**
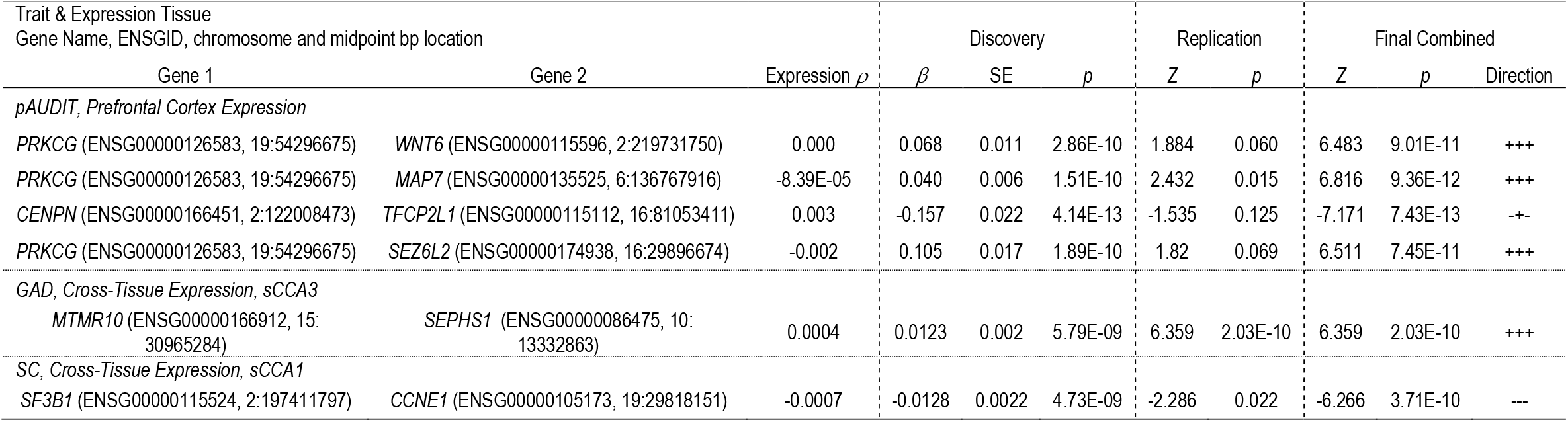
Interaction associations of pairs that reached *p*≤5.86e-10 in the final combined meta-analysis. Replication and final combined results indicate the meta-analyzed *Z* scores.

*MTMR10*x*SEPHS1* was significantly associated with GAD in the final meta-analysis. Both expressed in glial cells, *MTMR10* is in a locus associated with schizophrenia and dendritic growth deficiency^44,45^, substance use disorders and related behavioral traits^46^, while *SEPHS1* deregulation has been reported in rats under chronic stress^47^. SEPHS1 influences selenium metabolism pathways, deficiencies in which lead to oxidative stress^48^ and increased inflammation and degradation of extracellular matrix^49^. MTMR10 plays a role in the extracellular matrix, including in neurons and protects dendrites in response to oxidative stress^50^; their interaction may relate to regulation of inflammation and stress response. SC was associated (final meta-analysis *p*=3.71e-10, discovery *p*<5e-9, replication *p*=0.023) with an interaction between *CCNE1*, a cyclin family protein required for CDK kinase activity, and *SF3B1*, a component of the splicosome^51^, which may highlight the role of transcription regulation and modification with circadian cycles and the relationships of smoking cessation with sleep chronicity^52^ and disturbance^53^.

The limited number of significant interaction associations was not surprising given the low power to detect small effect sizes, particularly when expression imputation is imperfect and with stringent multiple test correction (Supplementary Figures S1-S4). As in single-locus GWAS, we anticipate additional, replicated loci to be identified with larger GWAS and expression reference panels, because imperfect expression imputation sharply reduces power.

For genes involved in at least one suggestive (*p*≤1e-5) interaction association, we found that across all traits, the number of interactions per gene followed a power-law distribution, with the majority of genes participating in only one or two interactions, but a few involved in many (“hub” genes, Fig. 3, Supplementary Fig. S21-32, Supplementary Table S2). Such highly connected genes (Table S2) represent logical targets for functional follow-up and characterization as hubs of interactions with many genes, integrating signals throughout pathways. While they may be poor drug targets as critical bottlenecks that impact multiple traits, identifying the genes they interact with could be a useful approach to find specific targets to modulate in developing therapeutics. The gene with the most interactions was, with pAUDIT using PFC expression, *FOLH1B*, an untranslated pseudogene previously associated with psychiatric disorders^54^ and BMI^55^. *PRKCG*, noted above, was the second most interacting gene, again with pAUDIT using PFC expression. The glutamate receptor *GRIK1* had the most interactions associated with CPD but was not identified by the GSCAN study^56^, demonstrating the novel associations to be found and the possible role of glutamate and excitatory neurotransmitters in smoking^57^. *RRAGA*, which regulates^58^ the mTOR signaling cascade^59^ that may have a role in the antidepressant effects of NMDA antagonists^60^, was the most interacting gene associated with GAD, highlighting the possible role of the mTOR pathway for internalizing disorder treatment. From a genetic architecture perspective, these findings support a long-standing hypothesis that while epistasis is common, most genes will interact with a limited number of other genes^7^. They also support an omnigenic model^61^ of architecture, where core or hub genes interact with and incorporate the regulatory effects of many peripheral genes. TWIS may identify such core or hub genes more directly than single gene models.

**Figure 3.**
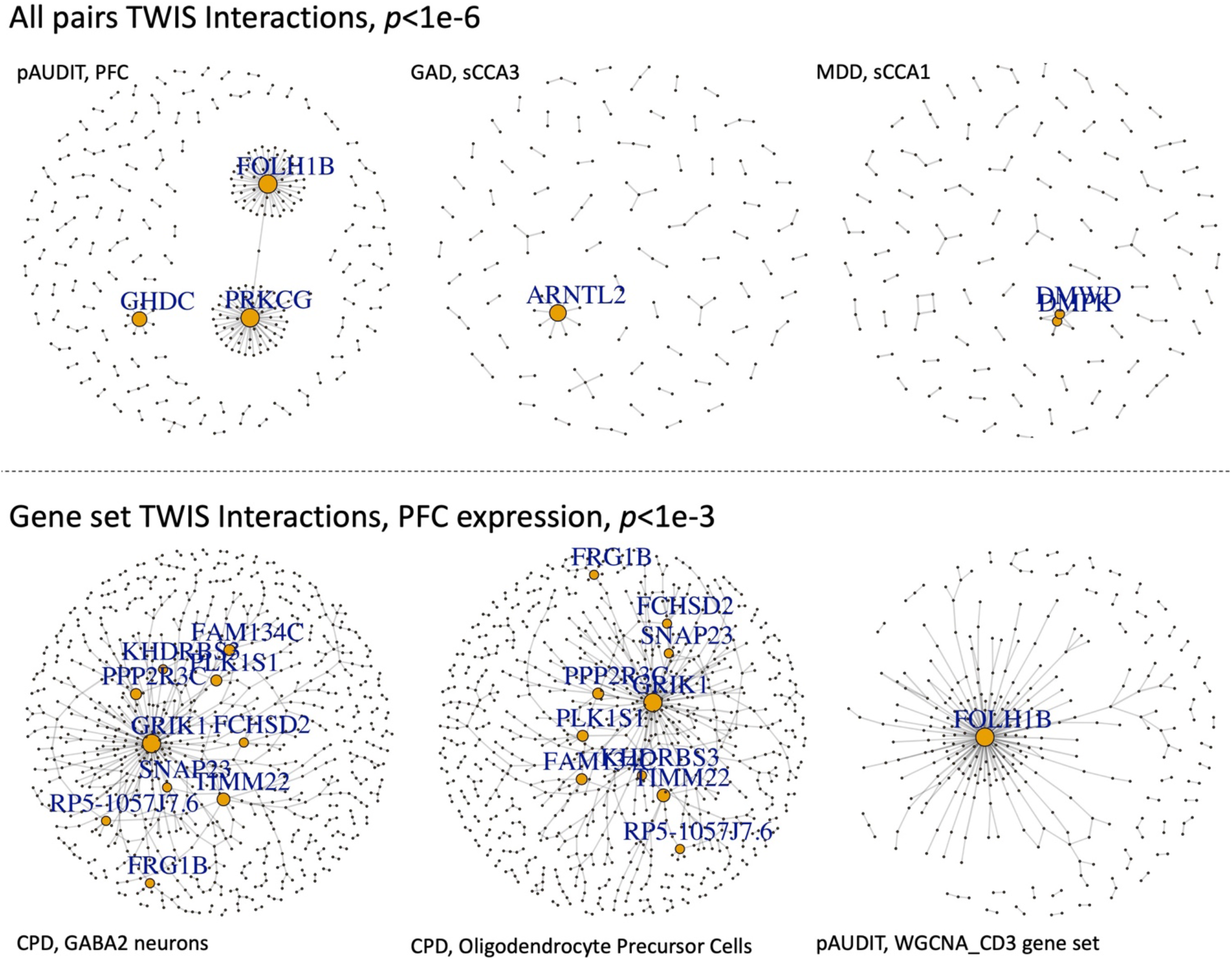
Networks of TWIS associations for selected traits and gene expression in specific tissues, either based on all pairs with *p*<1e-6 from the exhaustive, genome-wide TWIS (top), or within specific gene sets applying a nominal *p*<1e-3 threshold (bottom). Genes with degree≥5 are labeled, and size of points is proportional to node degree.

Given our exhaustive, all-pairs TWIS for multiple traits, we were also able to test whether genes with evidence of interaction association would have been identified in a single gene TWAS, as it is hypothesized the effect sizes of a locus could be diminished when analyzed individually if the gene’s effect depends on an interaction with another^7^. For example, the main effect of *GRIK1* on CPD was significant (*p*=2.93e-8) when the interaction with *PRRC2C* was modeled, but not in a single locus TWAS (*p*=0.35), nor was it identified in the largest CPD GWAS to date^56^. Using pairs of suggestive (*p*<1e-5) interaction associations in the combined meta-analysis, we estimated that, on average, only 3% (SD=6.8%) of the unique genes identified in TWIS would have been identified using a single gene TWAS (Fig. 4a, Table S3). As an example, of the 1106 unique genes in 655 pairs identified with GAD TWIS associations using PFC expression, none would have been associated in single locus models. Similarly, of the 981 unique genes in 547 interacting pairs associated with BMI using PFC expression, only 25 would have been identified in single-gene TWAS (Table S3). This results from reduced effect sizes in the single gene TWAS for genes with the largest interaction effects (Fig. 4b). This is consistent with the hypothesis that when a gene interacts with many others, its estimated effect in a single locus model may not be strong^7^, and it highlights the fact that novel loci may be identified using an exhaustive, all-pairs TWIS relative to single-locus TWAS or GWAS, with *GRIK1*, noted above, an example.

**Figure 4.**
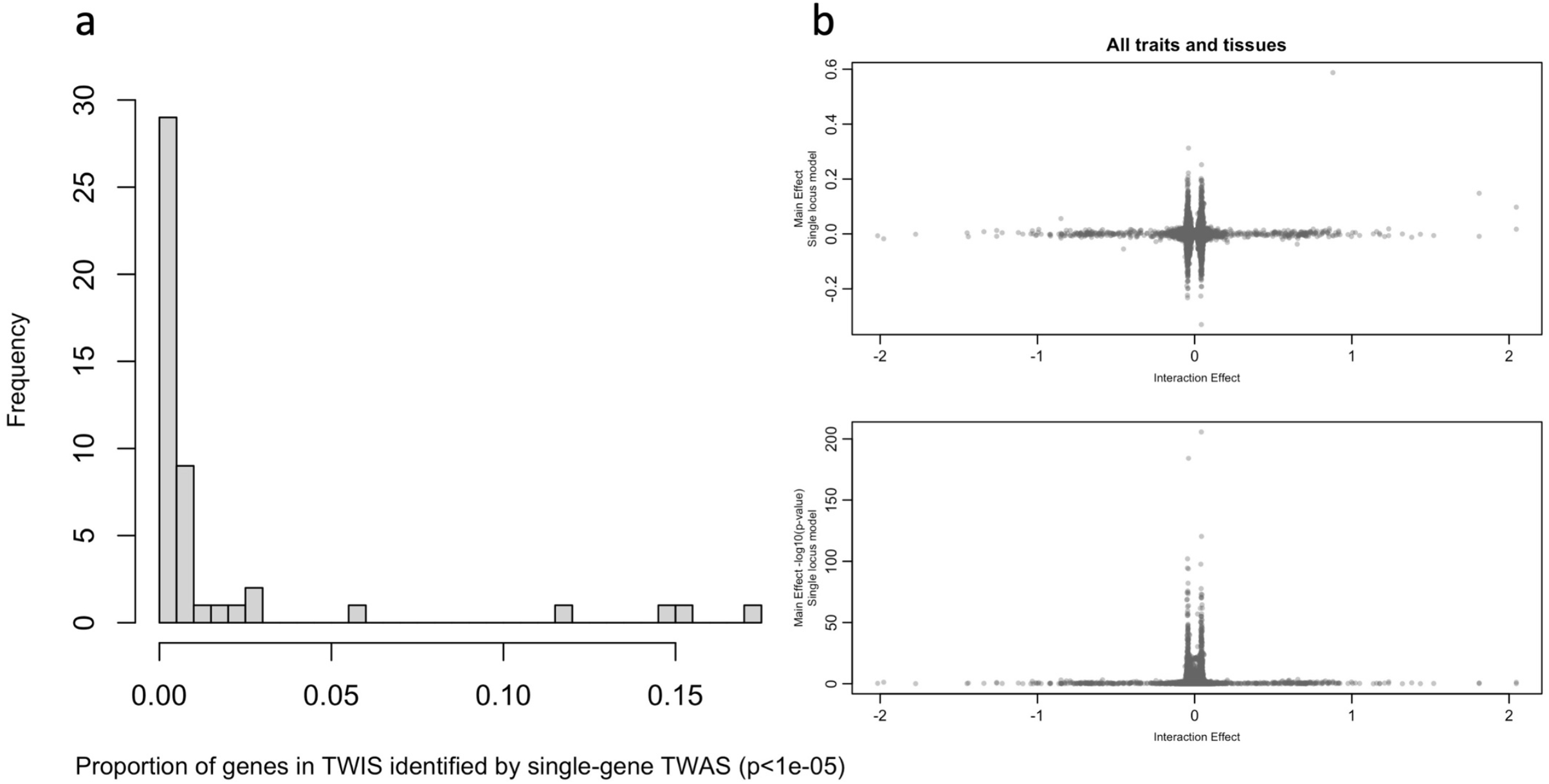
(a) Proportion of genes identified within suggestive interaction associations (p≤1e-5) that would have been identified using the same threshold in a single gene TWAS. Data in Supplementary Table 3. (b) Relationships of TWIS interaction effect sizes and main effect sizes of the same genes from TWAS (single locus model). Estimates of effects from all genes identified in TWIS included across traits and tissues, but each TWIS-identified gene is included only once per trait and tissue combination, even if a gene interacted with multiple other genes.

### Functional and Pathway Interaction Enrichment

We developed Enrichment-TWIS (E-TWIS) to assess the strength of interaction associations among genes within *a priori* defined gene sets of interest, rather than individual pairs of genes, including multiple functional pathways and networks. We first used a measure similar to network connectivity^62^ to use *χ*^*2*^ tests to efficiently test enrichment of approximately 8,000 gene sets. This was anti-conservative for large (n>150 genes) gene sets, where we used a random resampling approach to confirm enrichment (Supplementary Fig. S33-S35). The resampling represents a competitive test (*sensu* ref.^63^) of enrichment relative to background epistatic interactions, and in practice produced qualitatively similar results. We advocate an approach of efficiently testing many gene sets via *χ*^*2*^ tests and resampling to confirm large gene set enrichment.

Gene sets included the weighted gene coexpression network analysis (WGCNA) modules in PFC expression data^30,64^, many sets defined in the Molecular Signatures Database (MsigDB)^65^, and genes specifically expressed within individual cell types within multiple brain regions and subsets intolerant to protein-truncating mutations^66^.

We identified 50 significantly associated (FDR<5%) gene sets across all traits and expression tissues (Fig. 3, Fig. 5, Supplementary Table S4-S5). Among the associated gene sets, a common theme for several traits, notably GAD, PSYCH, neuroticism, CPD, and alcohol use, was enrichment of sets related to immune system and inflammation pathways. For neuroticism, we identified *STAT1* transcription factor binding sites as enriched, which regulates cellular responses to interferons, cytokines and other growth factors, and plays a role in immune response. Genes involved in immune system function (upregulated in T cells relative to B cells) were enriched in GAD, together suggesting the importance of immune system and inflammatory pathways for anxiety-related traits. Genes with expression influenced by *FOXP3*, which regulates immune system response including *IL2*, were enriched in psychiatric case epistatic interactions.

**Figure 5.**
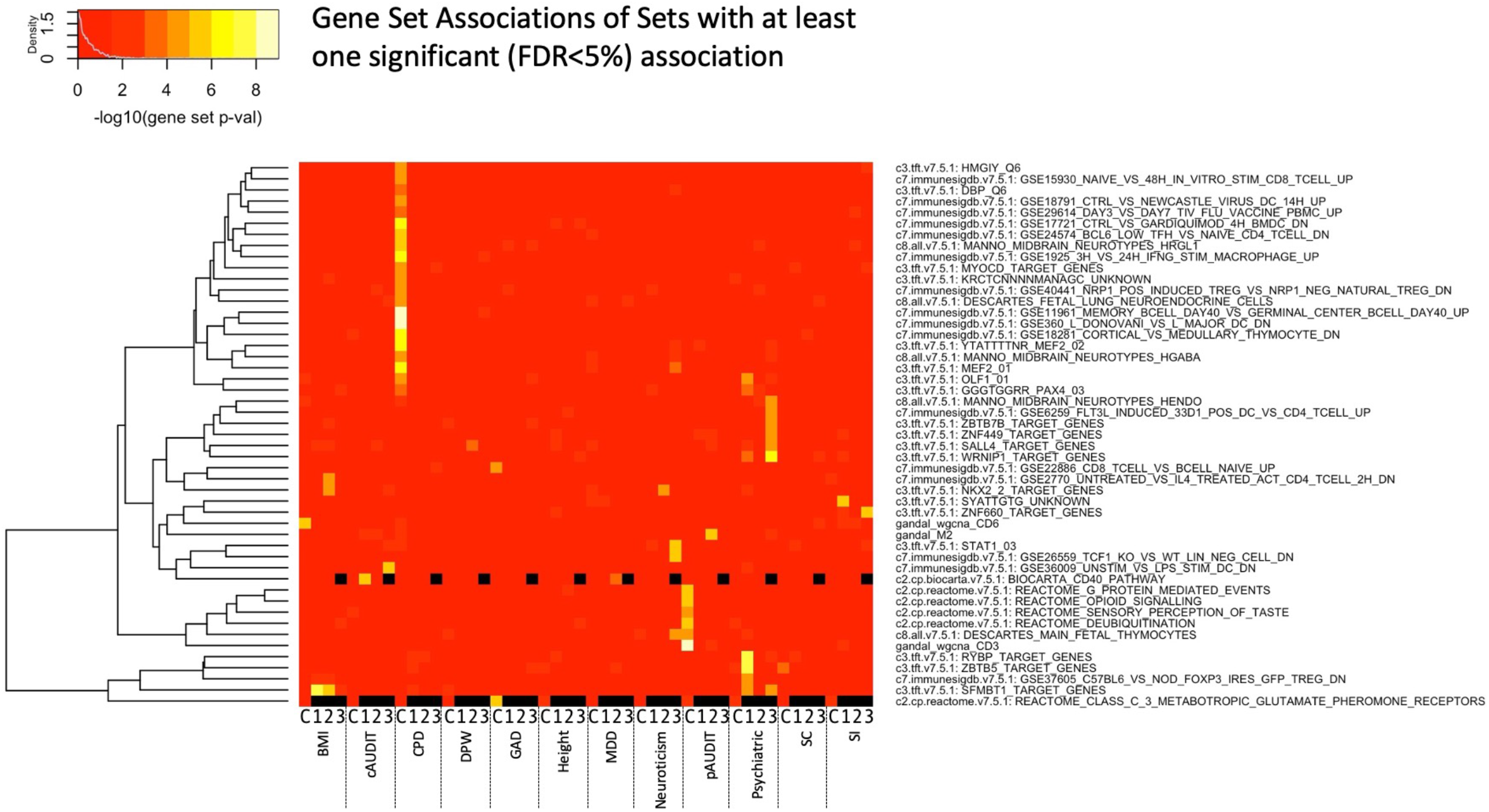
Gene set enrichment across all tissues and traits for those sets with at least one significant test (FDR<5%). Black indicates that the gene set association was not evaluated for that tissue and trait combination. X-axis shows the trait and tissue, where C indicates PFC and 1-3 represent the cross-tissue sparse canonical correlation axes 1-3. Phenotype details are in the Supplementary Materials.

Evidence of cell signaling pathway enrichment was also found, such as glutamate receptor genes for GAD (Supplementary Table S5). G-protein mediated event genes were enriched for pAUDIT, which includes signal transduction at the synapse, and is consistent with the *WNT6-PRKCG* interaction noted above (and possible immune function). Gene interactions within the deubiquitination REACTOME pathway were associated with pAUDIT, suggesting the importance of post-translational modification in alcohol use as has been hypothesized^67^, and highlighting the need for additional ‘omics integration into such analyses. Notably, three of the coexpression network modules identified by Gandal et al.^30,64^ were associated with BMI or pAUDIT. The gene M2 network (associated with pAUDIT) was found to be downregulated in oligodendrocytes in bipolar and schizophrenia cases^30^, while the CD3 module (also associated with pAUDIT) was found to be enriched in oligodendrocytes^64^, suggesting a role of glia.

Among gene sets specifically expressed in individual cell types^66^, we found enrichment of many traits for interactions in both excitatory and inhibitory neurons, with a number of GABAergic neuron enrichments (Fig. 3, Fig. 6, Supplementary Tables S6-S7). Notably, excitatory neurons were strongly enriched in CPD, supporting the individual strong interactions of *GRIK1* noted above. Oligodendrocytes and/or their precursor cells were enriched in BMI, CPD, height, MDD, and pAUDIT, highlighting a role of non-neuronal cells in several traits.

**Figure 6.**
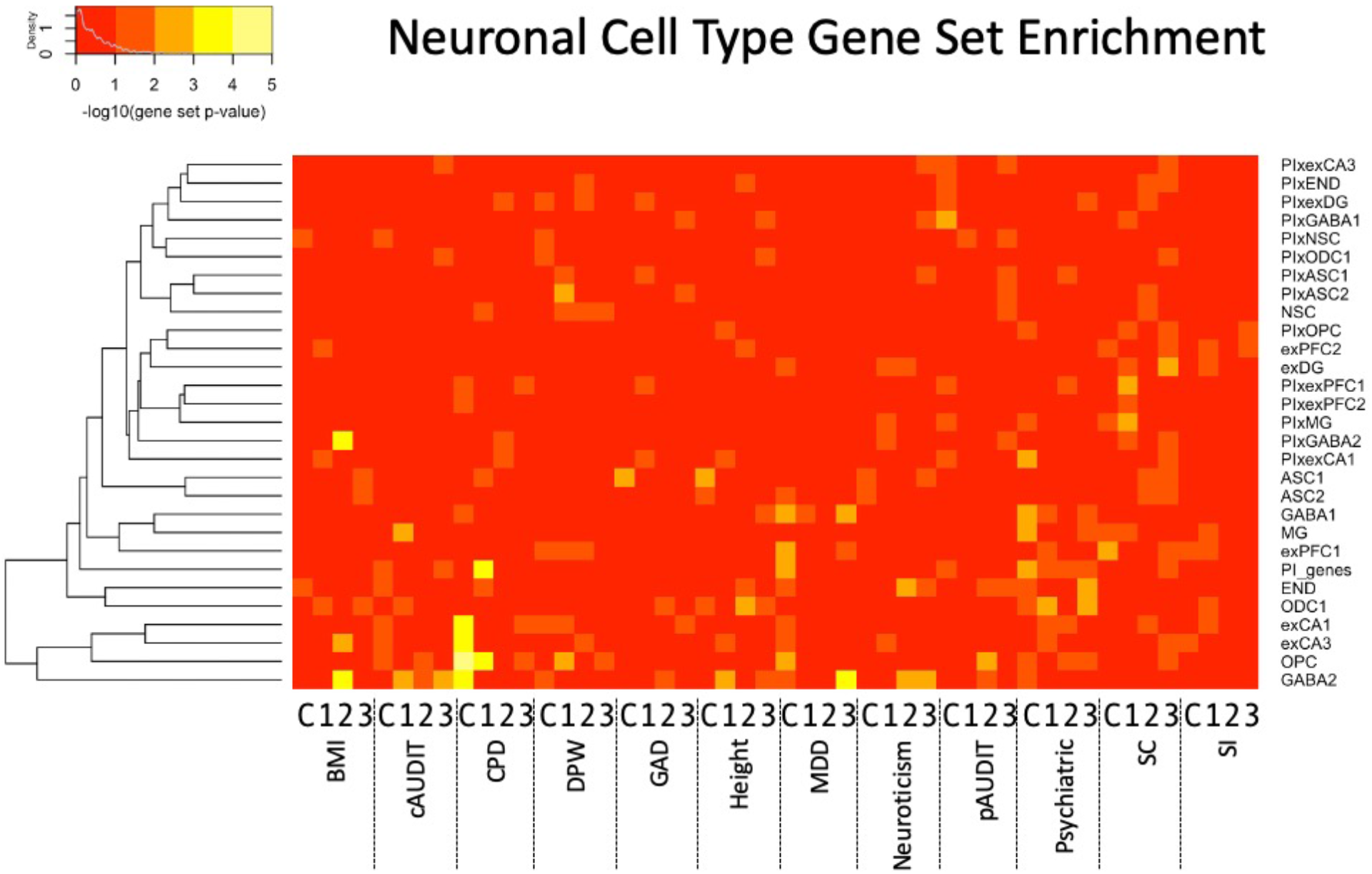
Neuronal cell type^66^ gene set interaction association enrichment across all tissues and traits. X-axis shows the trait and tissue, where C indicates PFC and 1-3 represent the cross-tissue sparse canonical correlation axes 1-3.

## Discussion

Here, we present the first, to our knowledge, fully exhaustive transcriptome-wide interaction study of all pairwise gene interaction associations. We confirmed several long-standing expectations of quantitative genetics, including that most genes have only a few interactions while a few ‘hub’ genes contain many, and that for genes with strong gene-gene interactions, estimated effects from a single-locus models are weaker. These two findings imply that epistasis is frequent, with key hub genes yet to be identified. These results also suggest that exhaustive interaction studies are needed rather than two-stage or variance models, which are efficient but may fail to detect real interactions. TWIS is an efficient way to both reduce the overall number of tests (on the order of 1e8 rather than 1e12 SNPxSNP tests) and improve power by integrating small individual SNP effects on expression. Although other approaches have been proposed^28,68,69^, we have built upon previous findings suggesting epistasis is important for complex traits, and provide a novel framework in which to exhaustively search all pairwise gene-gene interactions.

We also present findings of power analyses and type-I error, which verify both low power, as expected in interaction tests, as well as a need for stringent control of false positives. We confirmed that linkage disequilibrium (LD) and imperfect expression imputation and phenotype measurement can lead to false positive epistasis^21,22^. However, across extensive simulations, we were only able to inflate the type I error rate in the presence of LD; therefore, we apply a relatively simple yet robust approach to remove findings likely enriched for false positive interaction associations by excluding from analyses pairs of nearby genes and those with correlated imputed expression.

Despite these challenges, we identify genome-wide significant gene-gene interaction associations with problematic alcohol use, generalized anxiety disorder, and smoking cessation. This is proof-of-principle that the approach will identify novel interactions that can extend our biological understanding of complex traits, and as larger datasets and consortia become available, we anticipate additional epistatic associations will emerge.

Furthermore, when adopting a self-contained gene set-level approach^63^, we identified several significantly associated gene sets (Figures 4-5, Table S5-S7). We note that as a self-contained gene set analysis, this is testing a null hypothesis of no pairwise interaction association of genes within the gene set, rather than an enrichment of association signal relative to the background level of interaction associations (competitive gene set analysis^63^); computational constraints currently limit widespread E-TWIS competitive set analyses, but our follow-up resampling procedure performs such a competitive test, and we found qualitatively similar results.

Identified gene sets of interest include inflammatory and immune system pathways as relating to smoking, alcohol use, GAD and neuroticism; deubiquitination related to alcohol use suggesting the importance of epistasis for posttranslational modification; and multiple, notably glutamatergic, cell signaling pathways. Of particular interest, specific relevant cell populations can be identified using E-TWIS, and these include individual neuronal cells as well as glia.

### Limitations

Our exhaustive TWIS study has several notable limitations. First, we applied a linear regression-based statistical definition of epistasis. While computationally efficient, other models of epistasis could apply as well^23,24^, such as non-linear interactions among gene expression or dominance epistasis effects.

Second, LD leads to correlated tests and correlated predictors, which leads to complications in error control in interaction studies, increasing type I error and false associations of epistasis^21,22^. While standards for type I error correction have been generally accepted in single-SNP GWAS, there is no previous analogous standard for application to interactions. We have addressed this via extensive analyses of power and bias and have taken a conservative approach, removing any nearby pairs of loci and those with correlated imputed gene expression (|*r*|>0.05). This has likely removed true epistatic interactions, in which nearby, linked genes or intragenic loci interact^10,25,70^. While this prevents identification of physically proximate interactions, it removes a major source of LD-driven false positives^21,22^ which we view as necessary.

Third, the replication rate for epistasis tests is expected to be substantially lower than for additive tests, due to ascertainment of markers in LD with the causal variants and their chance resampling in independent datasets^10^. Nonetheless, we have applied rigorous replication thresholds, which we acknowledge likely result in higher rates of false negative replication. Combined with the stringent thresholds to remove LD-driven false positives, we are likely underestimating the extent of epistasis throughout the genome in complex traits; larger sample sizes will improve epistasis discovery.

Furthermore, scaling phenotypes in different ways (e.g., logarithmic) will impact the interaction estimates^9,71^. We residualized phenotypes and imputed expression, but the statistical epistasis identified here may be scale-dependent, and further mechanistic studies are required to determine the biological interactions at individual loci. Our analysis represents a computationally demanding, yet initial assessment of interactions throughout the genome.

Finally, assortative mating is expected to lead to correlation (i.e., LD) at functional loci even if they are physically separated^72,73^. We removed correlated loci, those in which assortative mating would be expected to lead to false positives. In this way, we expect assortative mating to not be a large driver of results here, but it is an area of future work worth exploring.

### Conclusions

Epistasis is likely widespread, but the computational challenges of so many pairwise tests have prevented its extensive examination. Here, we present a way forward using predicted gene expression, finding several significant interaction associations and multiple cell types and functional annotations enriched in epistasis affecting complex traits. We anticipate more to be identified as GWAS and expression reference panels continue to grow.

## WEBSITES

MsigDB: https://www.gsea-msigdb.org/gsea/msigdb/; FUSION: http://gusevlab.org/projects/fusion/; UKBiobank: https://www.ukbiobank.ac.uk/; dbGaP: https://www.ncbi.nlm.nih.gov/gap/

Code availability: code to be released publicly upon acceptance.

## DATA AVAILABILITY

All data used are from publicly available repositories, accessible via links above. All summary statistics will be deposited in a repository for public download.

## ACKNOWLEDGEMENTS

We thank the participants of the UK Biobank, NESARC-III, Genes for Good, ARIC, and GERA, and we thank the studies and their administrators. This research has been conducted using the UK Biobank Resource (application number 1665).

This work utilized the Summit supercomputer, which is supported by the National Science Foundation (awards ACI-1532235 and ACI-1532236), the University of Colorado Boulder, and Colorado State University. The Summit supercomputer is a joint effort of the University of Colorado Boulder and Colorado State University.

This work utilized the Blanca condo computing resource at the University of Colorado Boulder. Blanca is jointly funded by computing users and the University of Colorado Boulder. Data storage supported by the University of Colorado Boulder ‘PetaLibrary’. In particular, we thank Andrew Monaghan of CU Research Computing.

This work was supported by the National Institutes of Health (AG046938-06 to Chandra A. Reynolds, DA044283-01 to Scott I. Vrieze, MH100141-06 to Matthew C. Keller, DA017637 to John K. Hewitt, DA051937 and AA026733 to Marissa Ehringer, and AG064465 to Charles A. Hoeffer), the Linda Crnic Institute for Down Syndrome, and the University of Colorado Boulder Institute for Behavioral Genetics.

## ONLINE METHODS

### Description of TWIS Approach

We tested all pairs of gene-gene interactions using imputed gene expression after residualizing both the phenotype and expression on multiple covariates. This approach improved computation time while leading to unbiased estimates of the interaction effect. Details of each step are described below.

NB: Upon acceptance, scripts for each step will be made public.

#### Gene Expression Imputation in the Prefrontal Cortex (PFC) and Three Orthogonal Cross-Tissue Expression Measures

We imputed expression of genes in the PFC using the weights generated by PsychENCODE^30^ (14,729 genes) as well as three cross-tissue measures of expression^31^ (13,242; 12,521; and 12,032 genes for the three measures). We included the cross-tissue measures of expression (sparse canonical correlation analysis axes [sCCA] 1-3), as integration of data across multiple tissues increases reference sample sizes and improves power^31^.

TWAS weights were downloaded from the FUSION website for the PFC and cross-tissue expression measures (http://gusevlab.org/projects/fusion/). For each gene in each tissue, we first created score files of the best performing model weights using the make_score.R script (as outlined and available at the FUSION github site: https://github.com/gusevlab/fusion_twas). Following standard genotype QC (described below), we next extracted all SNPs in each cohort with non-zero expression weights using plink2^74^, followed by creating the individual-level expression prediction (plink2 --score command) for each gene’s expression.

### Residualization of Imputed Expression and Phenotypes

In interaction studies, proper control of covariates requires inclusion of all covariate-by-main effect terms^32^. This is critical when possible confounding variables exist. Therefore, we first examined a model for phenotype *y* in which, for imputed gene expression of two genes, *T*_*1*_ & *T*_*2*_, all main gene expression, expression interaction and covariate-by-gene expression terms were included:

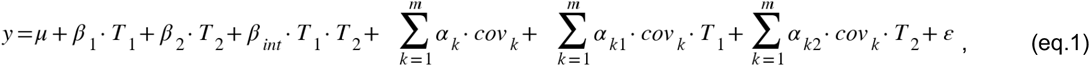

where *µ* is the intercept, *β*_*1*_ is the effect of expression of gene 1 (*T*_*1*_), *β*_*2*_ is the effect of expression of gene 2 (*T*_*2*_), *β*_*int*_ is their interaction effect, *α*_*k*_ is the effect of the *k*^th^ covariate (*cov*_*k*_), *α*_*k1*_ and *α*_*k2*_ are the interaction effects of the *k*^*th*^ covariate with *T*_*1*_ and *T*_*2*_, respectively, and *ε* is the error term.

Covariates include, depending on availability within each cohort (see methods), age, sex, genotyping batch, assessment center, socioeconomic variables such as income or education, and the first 10 genome-wide principal component axes. When many covariates are included, such as the large numbers of genotyping batches (106) and assessment centers (22) in the UK Biobank, all *m* covariates and their interactions with the main gene expression terms rapidly increases to hundreds of additional terms to estimate in the model for each pair of genes. This drastically increased computation time across many pairwise tests, particularly in samples of hundreds of thousands (e.g., the UK Biobank). Even with the reduced number of predictors (at the gene expression level) used here compared to all individual SNPs, all pairwise comparisons reach tens of millions of tests, e.g., ∼14,000 genes imputed using the PsychENCODE cortex expression weights^30^ results in ∼108M pairwise comparisons.

To improve speed, we therefore first residualized both the phenotype and genetically predicted gene expression on all covariates. This approach allowed us to remove covariate effects first, rather than repeatedly estimating them and their interactions for each pairwise test. Residualizing both predictor and response variables leads to unbiased estimates of the gene-gene interaction effect. We extracted the residuals from the following model:

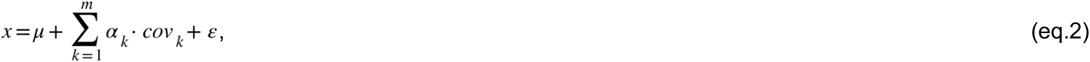

where *µ, α*_*k*_, *cov*_*k*_, and *ε* are as above, and *x* is either the phenotype (e.g., height) or the imputed gene expression (e.g., predicted *T*_*1*_). We used *fastLm* in the *RcppArmadillo*^*75*^ R package to fit the model efficiently for each imputed gene’s expression and continuously distributed phenotype, and the *glm* function to fit logistic regressions for each dichotomous phenotype. Residualized imputed expression and phenotype data were then merged into a single data frame.

*Exhaustive, All Pairs Gene-Gene Interaction TWIS*. Within each cohort, we then performed an exhaustive (all pairs) TWIS within each tissue for each trait using the following model:

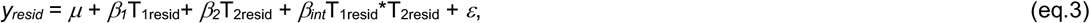

where *y*_*resid*_ indicates the residuals of phenotype *y* and T_1resid_ and T_2resid_ are the residuals of predicted gene expression of *T*_*1*_ and *T*_*2*_, respectively, from eq.2. We estimated *µ, β*_*1*_, *β*_*2*_, and *β*_*int*_ using fastLm in R. For each tissue and trait within each cohort, this amounted to *gp*=108,464,356; 87,668,661; 78,381,460; and 72,379,496 pairwise tests in the PFC and three cross-tissue expression measures, respectively, or 346,892,973 total pairwise tests for each trait in each cohort.

To expedite this step, we parallelized these tests across multiple compute nodes using the RMACC Summit Supercomputer at CU Boulder. For each combination of tissue, trait and cohort, we split the total tests into 1000 chunks, each of which was distributed to independent compute nodes. Each chunk therefore performed *gp*/1000 pairwise tests, which were indexed as the tests between pair (*a*[*k*], *a*[*i*+*k*+*d*\*(*d*+1)/2-*m*]), where *n*=number of total genes, *m=n*(n-1)/2, y=m-i, i* is the *i*^*th*^ chunk out of 1000, *d*=1+floor(((8\**y*+1)^0.5 -1) / 2), and *k=n-d*. This uniquely tested each pair only once, while distributing the computation to as many compute nodes as available on the supercomputer. Within each chunk, we further parallelized eq. 3 to multiple available CPUs using the *foreach* R library^76^.

### Discovery, Replication, and Meta-Analysis

We treated the UK Biobank as the discovery sample, and meta-analyzed results from the remaining cohorts for each phenotype as an independent replication sample. For meta-analysis, we applied the sample-size weighted approach of METAL^33^. We applied this rather than a traditional inverse-variance weighted meta-analysis because in several cases, the phenotypes in each cohort were approximate comparisons (e.g., “psychiatric disorder” based on ICD-9 & -10 codes (GERA, UK Biobank) vs. self-reported and DSM-V diagnoses of multiple disorders (ARIC, NESARC-III) and because the predictors and phenotypes were residualized on covariates prior to our TWIS, making SE-based meta-analysis inappropriate.

A full description of power and type-I error rates is in *Determining Alpha* and *Tests of Power and Biases* below. Based on those findings, we applied a significance threshold of *α*=5.86e-10. When pairs of genes are unlinked, this is the approximate 5^th^ percentile of minimum *p*-values from exhaustive genome-wide gene-gene tests under the null (see below). This is also very close to the Bonferroni correction threshold for all pairs of genes across the genome (i.e., ∼0.05/choose(20000,2)). Based on those findings and tests of biases described below, we restrict all subsequent analyses to pairs of genes whose physical position midpoints are greater than 1Mb apart and whose imputed expression is uncorrelated (|*r*| < 0.05), because linked and correlated pairs of genes lead to high rates of false positives. In our independent replication dataset, at pairs passing discovery significance, we applied a nominal *p*<0.05 as evidence of replication. Finally, we meta-analyzed all cohorts together (UKB+ replication cohorts). The complete meta-analysis results were utilized in subsequent gene set enrichment tests.

### Sample QC, stratification, PCA and relatedness

All cohorts (see supplemental note) included SNP array and/or imputed genome-wide SNP data. Genotype quality control (QC) of the array data included genotype missingness, Hardy-Weinberg Equilibrium tests, and minor allele frequency (MAF) using plink2 (command: -- geno 0.05 --hwe 0.00000001 --maf 0.01). For cohorts without imputed data, we utilized the Michigan Imputation Server to impute array data to the Haplotype Reference Panel^77,78^ after QC. Following imputation, then applied additional QC imputation metrics using plink2 (command: --extract-if-info R2 ‘>=’ 0.9 --maf 0.0001 --hwe 0.00000001 --geno 0.01 --mind 0.01).

Within each cohort, we identified a set of unrelated and relatively unstratified individuals matching (in terms of principal components analysis [PCA] axes) the expression reference panels, which are primarily European ancestry individuals. To reduce stratification effects and because expression imputation relies on sufficient matching of LD patterns between the target and reference panels^79^, we restricted our analyses to individuals of European ancestry, as that was both the largest relatively genetically homogeneous sample available across all cohorts and because the expression reference data were primarily derived from European ancestry individuals. We first used HapMap3 positions in the 1000 Genomes (1KGv3)^80^ reference panel to generate PCA loadings of the first 10 axes using flashpca^81^. We then extracted these same HapMap3 positions from each study cohort and projected them onto the 1KGv3 PC axes using flashpca. We then identified all individuals within +/-5 standard deviations of the 1KGv3 EUR population mean on each of the first four PCs, matching the approach applied by GSCAN^56^ across many cohorts, thus identifying a relatively unstratified set of individuals with LD patterns roughly matching those of the expression reference panels available.

We retained unrelated individuals using GCTA^82^ within each cohort after applying a pairwise relatedness cutoff of 0.05 using MAF- and LD-pruned SNPs (plink2 --maf 0.01 --indep-pairwise 50 5 0.2). See Supplemental Table S1 for final sample sizes for each cohort and each phenotype.

### Tests of Power and Biases

We performed a series of simulations to estimate power to detect interactions in the context of imperfect expression imputation across a range of epistasis effect sizes, define the appropriate alpha for genome-wide multiple test correction in the context of many millions of individual tests, and assess the role of linkage in influencing interaction tests.

#### Assessment of Power in the Context of Expression Prediction Error

Expression prediction is imperfect (Fig. S1). This is a function both of the heritability of the trait^83^ as well as sampling variance from finite (often small) expression reference panels^3-5^. To assess how such imperfect expression prediction impacted the power to detect gene-gene (GxG) expression interactions, we performed a set of Monte Carlo simulations (each 5,000 replicates) while varying the sample size (N=5000, 10000, 15000, 25000, 40000, 50000, 75000, 100000, 150000, 200000, 250000, 500000), the proportion of the phenotypic variance truly explained (PVE) by the interaction (PVE=0, 0.0001, 0.00025, 0.0005, 0.001, 0.005), and incorporating prediction error of the gene expression values by drawing randomly from the observed distribution of imputation accuracy (Fig. S1). We simulated gene expression values (the predictors in our model) from standard normal distributions, then generated phenotypes as a function of main and interaction expression effects and random noise, based on the set PVE. We then added error to the predictor expression values by drawing random noise from a ∼*N*(0, *σ*^2^_resid_), where *σ*^2^_resid_ was equal to one minus the observed prediction accuracy of a value randomly drawn from the distribution in Fig. S1. We performed these simulations with and without the added prediction error to assess its influence on bias and power.

As expected, decreased predicted expression accuracy decreased the power to detect significant interactions (Fig. S2). Note that when PVE=0, roughly 5% of tests were significant when using alpha=0.05 (and 0% with more stringent thresholds), indicating a well-calibrated interaction test statistic under these simulated conditions.

#### Assessment of Power Using Actual Predicted Gene Expression

We next used UK Biobank data, with genome-wide predicted gene expression, to incorporate real predicted expression data into our simulations. We used predicted sCCA1 expression data, and excluded individuals with relatedness > 0.05 (e.g., a sample similar to that used when testing epistasis effects on height). We randomly selected 5,000 pairs of genes from throughout the genome, and from the imputed expression data, simulated phenotypes as described above. We then added random noise to the imputed expression predictors, based on the estimated prediction accuracy (Fig. S1) for each gene in each pair. Again, power declined when error (due to imperfect expression prediction models) was added to the expression values used in the regressions (Fig. S3). Power was also decreased relative to the simulations described above (Fig. S2). Note that when PVE=0, roughly 5% of tests were significant when using alpha=0.05 (and 0% with other thresholds), indicating a well-calibrated interaction test statistic when incorporating data derived from real imputed expression data within a large biobank sample.

#### Assessment of Power Using Pairs of Physically Proximate Genes in LD

In the presence of imputation error, LD leads to an inflated false positive rate. We confirmed that, similar to recent reports^21,22^, this is due to binomially distributed predictors (i.e., true expression abundance when genetically based) with normally distributed error added (either from imperfect expression imputation or from random error) through a series of simulations varying LD, physical proximity and the distribution of the predictors (binomially distributed or normally distributed gene expression levels). We found evidence for this inflation only in the presence of LD. We next describe the two analyses we performed to conclude this.

Variation within nearby genes is expected to be correlated due to LD, and we expected that this could inflate test statistics, leading to false positives when comparing physically proximate genes based on other studies^21,22^. To understand how linkage impacts the test statistics, we therefore performed tests identical to those described above, but randomly chose only pairs of genes that were physically, immediately next to one another, thereby building into the simulations the desired physical proximity and underlying linkage. In these simulations (Fig. S4), power to detect true effects was slightly reduced relative to when pairs were randomly selected throughout the genome, but when prediction error was added to the expression values, we observed inflation of the Type I Error rate. When PVE=0, at the largest samples simulated, ∼40%, 7.5%, and 5.5% of tests were significant at alpha=0.05, 5e-8, and 2.5e-10.

To confirm LD as the cause of this, we simulated pairs of gene expression data from either a standard normal distribution (∼*N*(0,1)) or from a simple PGS (the sum of the minor alleles) of varying polygenicity (2, 10, 20, 50 or 100 SNPs per gene) derived from binomially distributed genotype data. We then generated phenotypes from the main effects of the simulated gene expression but without a true interaction. For each simulation, we tested the regression model interaction term, either using the simulated PGS (representing the simulated expression of each gene), or simulating imperfect expression prediction by adding normally distributed noise to the PGS (Fig. S5). When using the true PGS as the predictor (no predictor error), the interaction tests are well calibrated (Type I error rate ∼0.05 when applying alpha=0.05) regardless of trait architecture or linkage. When genes were unlinked, the interaction tests are also well calibrated. However, if the genes are linked (such as would occur for perfectly linked PGSs of nearby genes), type I error rates can become strongly inflated in the presence of imperfect expression. When using expression with added error (to mimic imperfectly predicted expression data), the false positive rate becomes much greater if the expression is predicted from a PGS generated from simulated, binomially distributed SNPs. The effect is greatest for a PGS derived from a few SNPs with poor prediction accuracy (high error variance added to the predictor), and declines as the expression polygenicity increases or the prediction accuracy improves. When estimated expression was derived from a standard normal distribution, the type I error rates were never inflated. This appears to be due to the combination of a binomially distributed predictor with added error variance, a situation that has been observed previously^21,22^.

This suggests that tests of nearby genes (those with linked SNPs as predictors) have inflated type I error rate and should be treated with caution. Unlinked genes (e.g., far away or on different chromosomes) are unaffected, and the type I error rate is well calibrated.

#### Assessment of Expression-based Interaction Tests When Causal Variant Effects Do Not Operate Via Expression

We assessed the impact of true genetic interactions that are not mediated via expression effects on the phenotypes. Predicted cross-tissue or tissue-specific expression data are essentially local PGSs, built from SNPs within localized physical windows. If there are true causal variants (CVs) that impact the phenotype directly (*not* through effects on gene expression) and are linked to SNPs that predict gene expression, it is possible that one could identify significant GxG expression PGS-based associations due to linkage, when in fact no *expression-based* interactions truly influence the phenotype.

We tested this by simulating SNP-by-SNP interaction effects on phenotypes, then testing models of either SNP-SNP interactions or expression PGS gene-gene interactions. In these simulations, there is a true genetic interaction effect via SNPs, but the phenotype is unaffected by genetic-based expression. We included two different scenarios to confirm that LD between the truly functional SNPs and the rest of the SNPs that contribute to the genetically predicted expression is what drives the TWIS associations, using either the SNPs with the locally maximal LD score or the SNPs with the locally minimal LD score as the truly interacting SNPs.

Consistent with expectations, we found this results in false positive associations of gene expression epistasis, which reflects the expression PGS tagging of true causal interactions (Supplementary Figure S6). Furthermore, the larger the LD scores of the interacting SNPs, the higher the false positive rate of TWIS associations. We note that this is a false positive in the sense that there are no *expression-mediated* interactions, but there is a true genetic interaction in these scenarios, so such false positives may still be of interest.

### Determining Alpha

The study-wide alpha based on a Bonferroni correction is approximately 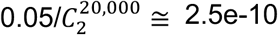 for a single trait and tissue expression combination assuming 20,000 genes in the genome, but these tests are not independent due to linkage and their pairwise nature. Furthermore, the influence of linkage, described above, clearly leads to inflated false positive rates. We estimated an appropriate genome-wide multiple test correction threshold by applying a similar simulation-based approach as has been used for univariate GWAS^84^. We simulated 100 independent genome-wide TWIS studies, each with 13,224 genes and 87,430,476 pairwise tests of epistasis (8,743,047,600 total tests) using the imputed sCCA1 expression in unrelated individuals from the UK Biobank, matching the sample size with height data (N=328,745). In each of the 100 datasets, we simulated phenotypes for each pair of gene-gene interaction tests, in which the genes had true main effects but no interaction effects, then estimated the interaction effect *p*-value using the approach described above. We identified in each of the 100 simulated TWIS studies the minimum *p*-value, then identified the 5^th^ percentile of these 100 minimum *p*-values as the appropriate genome-wide alpha. However, because LD varies throughout the genome and is expected to inflate false positive rates, we split this analysis into tests in which both genes are found on the same vs. different chromosomes, as a proxy for pairs possibly in LD vs. those not in physical LD. For 60 of these simulated TWIS studies, we further assessed whether the distribution of interaction *p*-values are drawn from an approximate cumulative *t* distribution across a range of pairwise expression correlations using Kolmogorov-Smirnov (K-S) tests, implemented in R. We found that the 5^th^ percentile of minimum *p*-values across the 100 simulated TWIS datasets from gene pairs on the same chromosome is 1.22e-20, reflecting the test statistic inflation due to LD between nearby genes noted above, while the 5^th^ percentile of minimum p-values from gene pairs on different chromosome is 5.86e-10, very similar to the alpha when using a Bonferroni correction (Supplementary Fig. S7). While predicted expression of all pairs of genes on different chromosomes were generally uncorrelated (most |*r*|<0.1) and the *p*-value distribution was not different from the expected cumulative *t* distribution, pairs on the same chromosomes had a range of pairwise expression correlation, and the distribution of *p*-values was increasingly dissimilar from expected at stronger pairwise expression correlations (Supplementary Fig. S8). Notably, across the 60 simulated datasets, the K-S test was not significant (almost all *p*>0.05) when, on the same chromosome, pairwise gene expression |*r*|<0.1, giving a threshold of pairwise expression correlation due to local LD, above which false positives are likely, but below which test statistics are reasonably well-calibrated. We therefore use a genome-wide, exhaustive TWIS corrected significance threshold of *p*<5.86e-10, while conservatively also excluding any pairs of genes whose imputed expression |*r*|>0.05 in the discovery sample (UK Biobank sample) and those pairs within 1MB.

### Enrichment-TWIS (E-TWIS)

We estimated enrichment of interaction associations within gene sets, rather than individual pairs of genes. We applied two separate approaches, first assessing enrichment of gene ontology^85,86^ (GO) categories of genes within pairs of suggestively significant interaction associations, which depended on statistical power to detect associations and yielded little information (data not shown). Second, we assessed the strength of interaction associations among genes within gene sets using a network analysis approach to determine the connectedness of all pairs of genes within *a priori* defined gene sets of interest, including multiple functional pathways and networks. Similar to network connectivity^62^, our measure summed the squared, meta-analyzed, pairwise interaction association *Z*-scores of all *m* pairs of *n* genes within each pathway or gene set, which was *χ*^*2*^_df=*m*_-distributed. To confirm that this approach produces appropriate *p*-values of gene set TWIS association enrichment, we performed simulations to estimate the distribution of gene set *χ*^*2*^ statistics under the null of no interaction association for several gene sets of varying size. These confirmed that our test statistic was *χ*^*2*^_df=*m*_-distributed for small (*n*<150) gene sets but was anti-conservative for very large gene sets (Supplementary Fig. S33). In these cases, we employed a secondary strategy, in which we randomly resampled *n* genes 1000 times, approximating the length and number of variants per gene in the target dataset, and averaged their *m* pairwise, squared TWIS *Z*-scores to estimate an empirical enrichment *p*-value. We confirmed similar findings to the *χ*^*2*^_*m*_ test (Supplementary Fig. S34-S35), noting that resampling represents a competitive test (*sensu* ref.^63^) accounting for background heritability throughout the genome via resampling random genes; therefore, annotated gene sets identified via random resampling are concluded to be enriched relative to background epistatic interactions. We advocate an approach of efficiently testing many gene sets via *χ*^*2*^ tests and resampling to confirm large gene set enrichment.

We tested a broad range of gene sets, including the weighted gene coexpression network analysis (WGCNA) modules in the PFC identified by Gandal et al.^30,64^ and multiple sets from the Molecular Signatures Database (MsigDB)^65^. The latter included hallmark gene sets; c2 canonical curated genesets from Biocarta, KEGG, and Reactome pathways; c3 regulatory target gene sets; c7 ImmuneSigDB gene sets; and c8 cell type signature gene sets. After excluding sets with fewer than 10 genes, we tested a total of 7,911-8,012 sets per trait and expression tissue and applied FDR≤0.05 multiple test correction. We then used the same approach to assess interaction association enrichment in genes specifically expressed within individual cell types within multiple brain regions and subsets of those genes that are intolerant to protein-truncating mutations (defined in ref.^66^).

### Number of Interactions Per Gene

To examine the distribution of interaction frequency per gene, we applied a nominal significance threshold of interaction *p*≤1e-5. We then evaluated the number of interactions each gene was involved in by plotting the distribution. As demonstrated by our power simulations, we are underpowered to detect strict Bonferroni-significant interactions, but as demonstrated by our gene set enrichment analyses, there is a signal of interaction associations within tests that do not reach strict significance, which is why we used a nominal *p*≤1e-5 threshold.

### TWIS vs. TWAS Comparison Using UK Biobank Data

We assessed whether genes identified in TWIS would have been identified using single gene TWAS^3^, as it has been hypothesized the effect sizes of a locus could be muted when analyzed individually if the gene’s effect depends on an interaction with another gene^7^. We restricted our analysis to genes within pairs of suggestive (*p*≤1e-5) interaction associations in the combined meta-analysis, across any phenotype and trait combination. We used the residualized UK Biobank data and applied a *p*≤1e-5 suggestive significance criteria. Using these results, we compared the interaction effect sizes from TWIS for each gene with its TWAS-estimated effect size to test whether genes with larger interactions have smaller effect sizes estimated in a single-locus model.

