## Supplemental Material for "Transcriptome-Wide Gene-Gene Interaction Association Study Elucidates Pathways and Functional Enrichment of Complex Traits"

#### Table of Contents

1. Supplementary Note: Cohort and phenotype descriptions: pages 2-5
2. Supplementary Figures 1-35: pages 6-38.

### Supplementary Note: Cohorts and Phenotypes Description

**Atherosclerosis Risk in Communities Study (ARIC)**

We accessed the Atherosclerosis Risk in Communities Study (ARIC) data in dbGaP (accession number phs000280.v7.p1), including genotype and phenotype data (approved dbGaP project number 20774). Sample sizes reported below are the final N after QC, removing related individuals, and merging with genetic/imputed expression data as described above.

Height (measured as standing height in cm) was from the Visit 1 ANTA dataset in field ANTA01 (N=7638). Body mass index (BMI) was obtained from the Visit 1 DERIVE13 dataset in field BMI01 (N=7655). Drinks per week (DPW) was the sum of the total number of beer, wine and liquor drinks per week. We first used the Dietary Intake dataset from Visit 1 (DTIA), using fields DTIA96, DTIA97, and DTIA98. For those with missing data in DTIA, we checked the Visit 3 PHXA04 and Visit 4 ALC datasets, using fields PHXA44A, PHXA45A, PHXA46A, ALC5, ALC6, and ALC7 (N=5747). Frequency of alcohol consumption was only available within the PHXA04 and ALC datasets. Thus, for cAUDIT, we used visit 3 PHXA04 and visit 4 ALC datasets, which contained data for cAUDIT items 1 and 2 in the above fields as well as fields PHXA44B, PHXA45B, PHXA46B, ALC5A, ALC6A, ALC7A. cAUDIT item 3 (how often one consumes 6+ drinks) was not directly asked within AIRC; therefore, we substituted DPW/days per week drinking of 6+ and frequency to obtain a similar binned variable of item 3. cAUDIT was the sum of these three derived variables (N=5747). There were no relevant AUD or pAUDIT fields in ARIC. Smoking initiation (SI) data were drawn from the Visit 1DERIVE13 dataset field CIGT01, which assessed whether an individual had ever smoked 400+ cigarettes in their lifetime and whether they currently, formerly or never smoked. Current or former smokers were considered to have initiated smoking (N=4915). Smoking cessation (SC) was derived from the same field, with never smokers coded as missing, and former smokers as cases (=1) and current smokers as controls (=0) (N=4915). Visit 1 Home Interview (HOM) dataset contained quantity of use in terms of cigarettes per day (CPD), either currently (field HOM32) or on average over the course of the entire time one smoked (field HOM35). We used the maximum of these two fields as a discretized, heavy/light smoker phenotype, cpdL10H20, indicating those whose CPD was either <=10 (=0) or >=20 (=1) (N=4455), as this is phenotype has a higher heritability[1] than the raw CPD count and likely reflects those addicted to nicotine vs. those who smoke but are not addicted. Depression (mdd) cases were identified as “Y” in the Visit 5 Neuropsychiatric Inventory dataset (field NPI4A) or “Y” in the Visit 5 Neurologic History (field NHX6) (N=1609), while anxiety (gad) was defined as “Y” in field NPI5A. Any psychiatric case was identified as responding “Y” to depression, anxiety, delusions, or hallucinations (NPI4A, NPI5A, NHX6, NPI1A or NPI2A) (N=1609). Covariates included categorical education (DERIVE13 field ELEVEL01), age at visit 1 (DERIVE13 field V1AGE01), and sex (DERIVE13 field GENDER), as well as total combined family income from the Home Interview (field HOM62), as well as the first 10 genetic principal components, as previously described. PCA of the covariate matrix was performed and the axes that explained ≥0.5% of the variance were retained as covariates.

**GENES FOR GOOD (G4G)**

We utilized the Genes for Good (G4G) study[2], obtaining approved access through the University of Michigan (https://genesforgood.sph.umich.edu/) in 2019. Covariates included age (field ID: age_range), sex (gender), education (education), household income (income), and the first 10 axes from the genomic principle components analysis (described above), and were included as fixed effects covariates. All were treated as categorical variables, with missing data coded as a separate category. PCA of the covariate matrix was performed and the axes that explained ≥0.5% of the variance were retained as covariates. Drinks per week (DPW) was derived from field drinks_per_day_in_last_30 and days_with_drink_in_last_30 and log-transformed as in GSCAN. As in other cohorts, AUDIT items were not directly asked. cAUDIT was indexed by days_with_drink_in_last_30 and drinks_per_day_in_last_third. As no question directly addressed item 6, we calculated a proxy variable by indexing those reporting 6 or more drinks per day with their frequency of drinking. pAUDIT questions were not addressed in the G4G survey data, with the exception of one relevant question: “Did you ever become tolerant to alcohol; that is, you drank a great deal more in order to get an effect, or found you could no longer get high on the amount you used to drink?" We used this single question as an index of problematic drinking in G4G. Height was obtained from height and BMI from height and weight. The data freeze we accessed asked “Have you ever been depressed (regardless of whether you received a formal diagnosis)?”, but the endorsement rate of this was ~79%, far different from endorsement rates or prevalence rates in the other datasets, and were therefore chose not to utilize G4G for the MDD phenotype.

Similarly, no appropriate survey questions for GAD were available. However, eight questions relevant to neuroticism, as defined by[3] were asked ("is_depressed", "is_relaxed", "can_be_tense", "worries", "is_emotionally_stable", "is_moody", "remains_calm", "gets_nervous"), and we indexed these to obtain a score of neuroticism.

**Genetic Epidemiology Research on Aging (GERA)**
We accessed The Resource for Genetic Epidemiology Research on Aging (GERA) Cohort genotypes and phenotypes through dbGaP, Study Accession: phs000674.v3.p3. GERA is part of the Kaiser Permanente Research Program on Genes, Environment, and Health (RPGEH) and the UCSF Institute of Human Genetics (AG036607; Schaefer/Risch, PIs).
We used two available “conditions” phenotypes from GERA – Depression (DEPRESS) and Psychiatric: any (PSYCHIATRIC). These two phenotypes were the only “condition” phenotypes similar to phenotypes available in the other cohorts. These were defined by GERA and released as case/control data based on ICD-9-CM codes. Per GERA data descriptions, these were, “Coded with disease if participant had at least two diagnoses in a disease category recorded on separate days. Diagnoses obtained from patient encounters at Kaiser Permanente Northern California facilities from January 1, 1995 to March 15, 2013.” Please see dbGaP phs000674.v3.p3 for details on the structure of the GERA study. Income, age, sex, education (all from the dbGaP-released “survey” dataset), and the first 10 axes from the genomic principle components analysis (described above) were included as fixed effects covariates. Income, age, and education were treated as categorical variables, with missing data coded as a separate category. PCA of the covariate matrix was performed and the axes that explained ≥0.5% of the variance were retained as covariates.

**National Epidemiologic Survey on Alcohol and Related Conditions-III (NESARC-III)**
We accessed the National Epidemiologic Survey on Alcohol and Related Conditions-III (NESARC-III) genotype and phenotype data through dbGaP, Study Accession: phs001590.v2.p1. See dbGaP phs001590.v2.p1 for details on the structure of the NESARC-III study. Covariates included age (field ID: NAGE), sex (SEX), education (NEDUC4), household income (NINCFAM3), and the first 10 axes from the genomic principle components analysis (described above), and were included as fixed effects covariates. Income, age, and education were treated as categorical variables, with missing data coded as a separate category. PCA of the covariate matrix was performed and the axes that explained ≥0.5% of the variance were retained as covariates. Height and BMI were obtained from variables NBMI, NFEET, and NINCHES. Smoking Initiation (SI) was defined following GSCAN definitions, from items N3AQ1A, N3AQ1B, and N3AQ1C; cigarettes per day (CPD) from fields N3AQ3C11 and N3AQ6A1; and smoking cessation (SC) from fields N3AQ3A1, N3AQ3A2, and N3AQ3A3. Drinks per week (DPW) was derived from field N2AQ4A (how often did you drink), restricted to those reporting a drink in the last year, therefore excluding former drinkers, and field N2AQ4B (how many on days that you drink). AUDIT items were not asked directly in the NESARC-III, but all items were approximated by questions within the NESARC-III. cAUDIT was derived from N2A4Q4A (cAUDIT item 1), N2AQ4B (cAUDIT item 2), and N2AQ4F and N2AQ4H (cAUDIT item 3). Item 3 in NESARC-III was separated between females & males aged 65+ or males aged below 65, and used a 4 vs 5 drink threshold rather than the AUDIT 6+ threshold. pAUDIT scores were generated by matching as best as possible to AUDIT questions. Item 4 was approximated by N2BQ1B6 & N2BQ1B7; item 5 by N2BQ1B30, N2BQ1B31, and N2BQ1B32; item 6 by N2BQ1B19; item 7 by N2BQ1B25 & N2BQ1B26; item 8 by N2BQ1B27; item 9 by N2BQ3A3 & N2BQ3A4; item 10 by N2CQ4A and N2CQ1. Thus, pAUDIT and cAUDIT scores are approximate, but index the degree of problematic drinking and alcohol consumption by participants. MDD cases were identified by item lmddisorder, and GAD cases from item lgadind. We identified NESARC-III individuals as ‘psychiatric’ cases matching the GERA psychiatric case definitions as closely as possible. The list of NESARC-III codes and descriptions included as cases are in the table below.

**Table.** NESARC-III Phenotype Codes included in psychiatric case phenotype.

| code | description |
|---|---|
| anndx | Past year DSM-5 anorexia nervosa |
| antisoc | DSM-5 antisocial personality disorder |
| antisocso | DSM-5 antisocial personality disorder with soc/occ |
| beddx | Past year DSM-5 binge-eating disorder |
| bipolar1 | Lifetime DSM-5 bipolar 1 disorder (hierarchical) |
| bpddx1 | DSM-5 borderline personality disorder (at least 1 criterion soc/occ) |
| bpddx2 | DSM-5 borderline personality disorder (at least 2 criteria soc/occ) |
| bulnerdx | Past year DSM-5 bulimia nervosa |
| conduct | DSM-5 conduct disorder without soc/occ |
| conductso | DSM-5 conduct disorder with soc/occ |
| lagoraind | Lifetime DSM-5 agoraphobia |

| | |
|---|---|
| lanndx | Lifetime DSM-5 anorexia nervosa |
| lbeddx | Lifetime DSM-5 binge-eating disorder |
| lbulnerdx | Lifetime DSM-5 bulimia nervosa |
| ldysind | Lifetime DSM-5 dysthymia (nonhierarchical) |
| ldysthymia | Lifetime DSM-5 dysthymia (hierarchical) |
| lgadind | Lifetime DSM-5 generalized anxiety disorder |
| lhypoind | Lifetime DSM-5 hypomanic episode (nonhierarchical) |
| lmanicind | Lifetime DSM-5 manic episode (nonhierarchical) |
| lmddisorder | Lifetime DSM-5 major depressive disorder (hierarchical) |
| lmdepind | Lifetime DSM-5 major depressive episode (nonhiercharical) |
| lpanicind | Lifetime DSM-5 panic disorder |
| lptsd | Lifetime DSM-5 posttraumatic stress disorder |
| lsocind | Lifetime DSM-5 social phobia |
| lspeind | Lifetime DSM-5 specific phobia |
| panndx | Prior to past year DSM-5 anorexia nervosa |
| pbeddx | Prior to past year DSM-5 binge-eating disorder |
| pbulnerdx | Prior to past year DSM-5 bulimia nervosa |
| ppyagoraind | Prior to past year DSM-5 agoraphobia |
| ppydysind | Prior to past year DSM-5 dysthymia (nonhierarchical) |
| ppygadind | Prior to past year DSM-5 generalized anxiety disorder |
| ppyhypoind | Prior to past year DSM-5 hypomanic episode (nonhierarchical) |
| ppymanicind | Prior to past year DSM-5 manic episode (nonhierarchical) |
| ppymdepind | Prior to past year DSM-5 major depressive episode (nonhiercharical) |
| ppypanicind | Prior to past year DSM-5 panic disorder |
| ppyptsd | Prior to past year DSM-5 posttraumatic stress disorder |
| ppysocind | Prior to past year DSM-5 social phobia |
| ppyspeind | Prior to past year DSM-5 specific phobia |
| pyagoraind | Past year DSM-5 agoraphobia |
| pybipolar1 | Past year DSM-5 bipolar 1 disorder (hierarchical) |
| pydysind | Past year DSM-5 dysthymia (nonhierarchical) |
| pydysthymia | Past year DSM-5 dysthymia (hierarchical) |
| pygadind | Past year DSM-5 generalized anxiety disorder |
| pyhypoind | Past year DSM-5 hypomanic episode (nonhierarchical) |
| pymanicind | Past year DSM-5 manic episode (nonhierarchical) |
| pymddisorder | Past year DSM-5 major depressive disorder (hierarchical) |
| pymdepind | Past year DSM-5 major depressive episode (nonhiercharical) |
| pypanicind | Past year DSM-5 panic disorder |
| pyptsd | Past year DSM-5 posttraumatic stress disorder |
| pysocind | Past year DSM-5 social phobia |
| pyspeind | Past year DSM-5 specific phobia |
| spddx1 | DSM-5 schizotypal personality disorder (at least 1 criterion soc/occ) |
| spddx2 | DSM-5 schizotypal personality disorder (at least 2 criteria soc/occ) |

**UK BIOBANK:**

UK Biobank[4] data was accessed via approved application number 1665.

Covariates included sex (UK Biobank field ID 31), age (21003), age$^2$, Townsend deprivation index (189), educational attainment (6138), genotyping batch (22000), scores of the first 10 worldwide principal components (22009), and scores of the first 10 genomic principal components. PCA of the covariate matrix was performed and the axes that explained ≥0.5% of the variance were retained as covariates.

Height was obtained from field 50 and BMI was obtained from field 21001. Drinks per week (DPW) was defined as the average number of weekly drinks for current or former drinkers, as in GSCAN[5]. cAUDIT was the sum of AUDIT items 1-3, corresponding to fields 20414, 20403, 20416. pAUDIT was the sum of AUDIT items 4-10, corresponding to fields 20413, 20407, 20412, 20409, 20408, 20411, and 20405.

Smoking initiation (SI) represented those who had smoked at least 100 cigarettes over their lifetimes. Smoking cessation (SC) was defined from fields 1239 and 1249. Heavy vs. light cigarettes per day (CPD) was defined from fields 2887, 3456, and 6183 and dichotomized as individuals who smoke more than 20 cigarettes per day vs individuals who smoke 10 or less, excluding intermediate individuals.

DSM-V-like MDD and GAD were defined from the UK Biobank mental health questionnaire: GAD DSMV-like cases required endorsement of either Field IDs 20425 or 20542, and endorsement of 20421 with 20420 reported as ≥56 months, and endorsement of 20540 or 20543≥2, and endorsement of 20541 or 20537 or 20539, as well as three or more “Yes” responses to the following symptom Field IDs: 20426 or 20423, 20429, 20419, 20422, 20417, 20427, and endorsement of ‘a little’ or more of field 20418 (impairment or impact). Similarly, MDD DSMV-like cases required “Yes” responses to 5 or more of the following symptom Field IDs: 20446, 20441, 20533, 20534, 20535, 20449, 20536, 20450, 20435, and 20437, as well as “somewhat” or more response to field 20440, a “almost every day” or more response to field 20439, and a “about half of the day” or more response to 20436. Neuroticism was obtained from field 20127.

We identified individuals as ‘psychiatric’ cases matching the GERA psychiatric case definitions as closely as possible. For this, we extracted individuals with ICD9 or ICD10 codes (fields 41270 and 41271, codes F20, F21, F22, F23, F24, F25, F28, F29, F30, F31, F32, F33, F34, F39, F40, F41, F43, F45, F50, 292, 295, 296,

297, 298, 300, 3071, 3075, or 3098) or those who self-reported having been diagnosed by a doctor any of related disorders (field 20544, codes 1,2,3,4,5,6,10,11,12,13,14,15,16, or 17). To these we added individuals identified as DSM-5-like MDD or GAD cases (see above). All individuals with any record of any of the above codes were considered cases. All others with non-missing data in at least one of the fields and with no endorsement across any field were considered controls. In the entire UK Biobank, this resulted in 43,459 cases and 459,052 controls (8.6% prevalence).

### Supplementary Figures

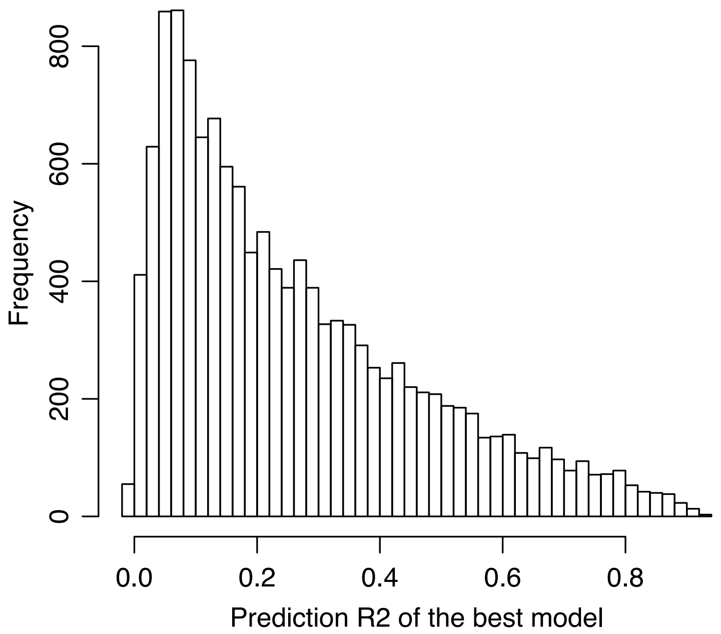


**Figure S1.** Histogram of the best model (i.e., lowest p-value) from FUSION output of the cross-tissue expression prediction models (first sCCA axis).

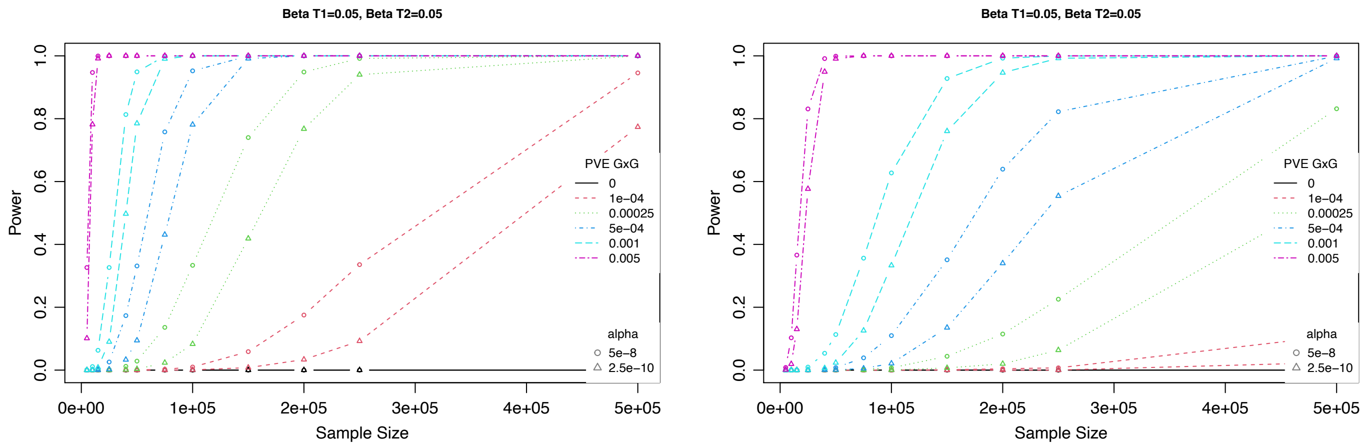


**Figure S2.** Power to detect significant interactions at two significance thresholds across varying sample sizes and true proportions of phenotypic variance explained (PVE) by the interaction. Left, without incorporating expression prediction error into the simulation. Right, incorporating random error for each predicted gene expression based on the distribution of observed prediction accuracies of the best model in Figure S1. Main effect sizes for the two expression predictors (T1 & T2) are shown above each plot; varying these had minimal effect on the interaction test power (data not shown).

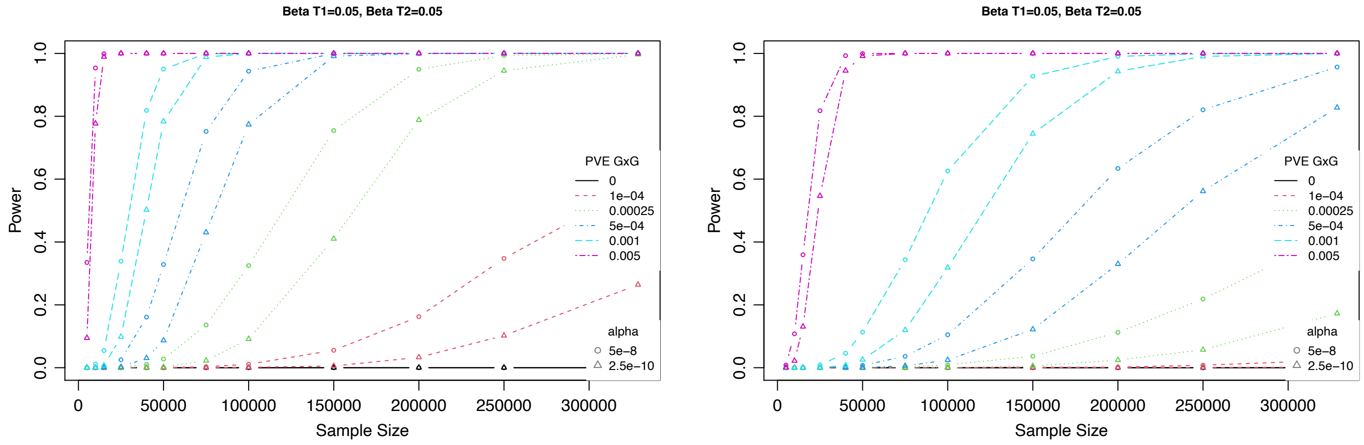


**Figure S3.** Power to detect significant interactions at two significance thresholds across varying sample sizes and true proportions of phenotypic variance explained (PVE) by the interaction when using predicted sCCA1 expression within the UK Biobank in unrelated individuals using pairs of genes randomly selected throughout the genome. Left, without incorporating expression prediction error into the simulation. Right, incorporating random error for each predicted gene expression based on the observed prediction accuracies of the best model in Figure S1. Main effect sizes for the two expression predictors (T1 & T2) are shown above each plot; varying these had minimal effect on the interaction test power (data not shown).

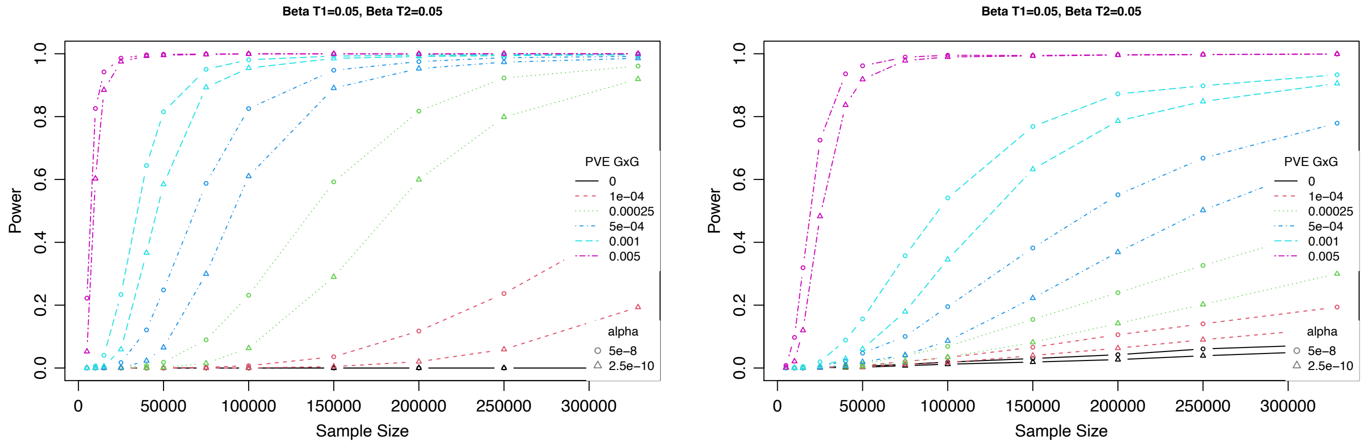


**Figure S4.** Power to detect significant interactions at two significance thresholds across varying sample sizes and true proportions of phenotypic variance explained (PVE) by the interaction when using predicted sCCA1 expression within the UK Biobank in unrelated individuals using pairs of genes that were immediately next to one another in the genome. Left, without incorporating expression prediction error into the simulation. Right, incorporating random error for each predicted gene expression based on the observed prediction accuracies of the best model in Figure S1.

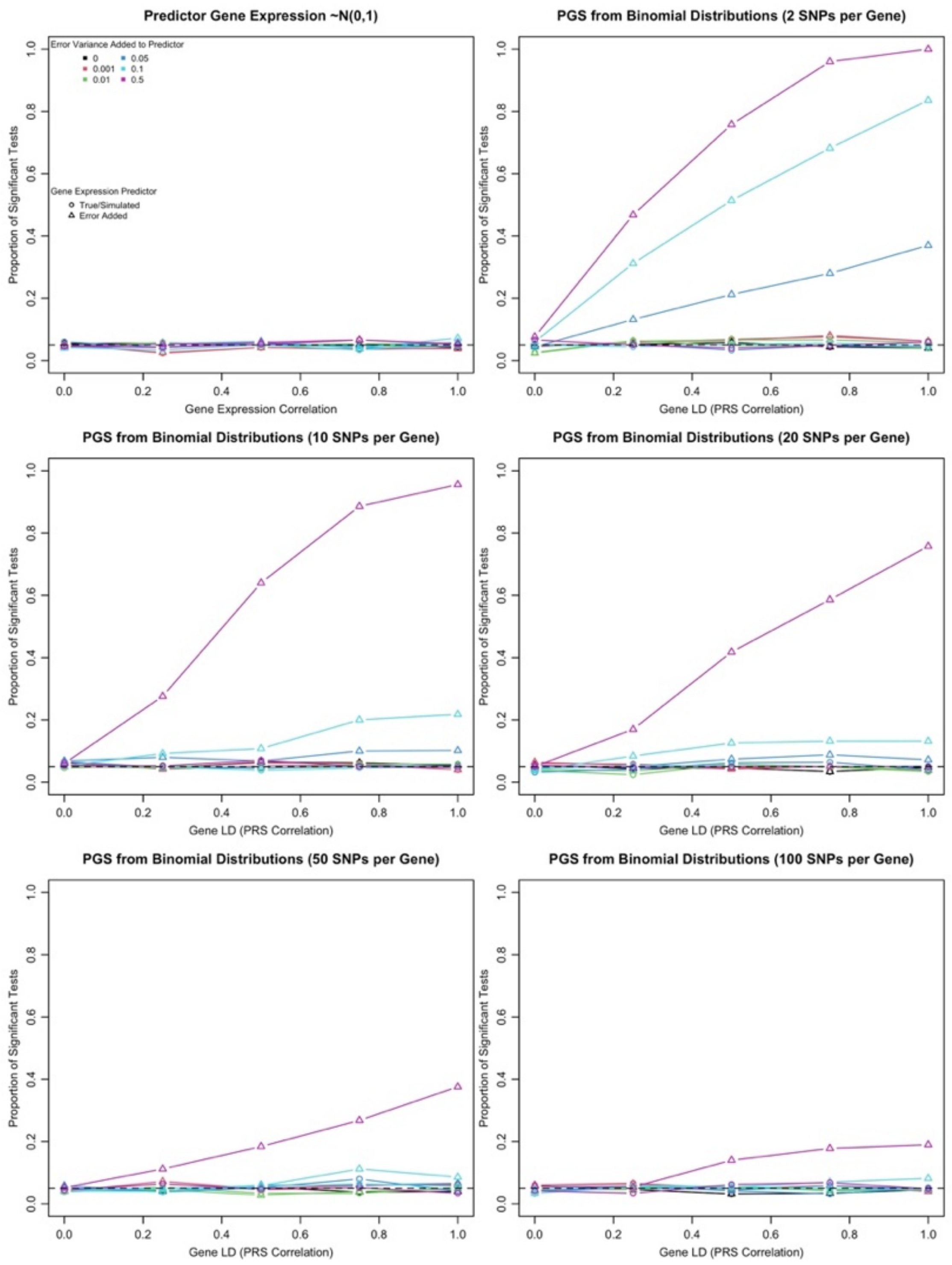


**Figure S5.** The proportion of significant tests (alpha=0.05) when simulating gene expression from linked (correlation of expression or gene LD = 1) or unlinked (=0) data, either from a standard normal distribution (~N(0,1)) or from a simple PRS of varying polygenicity. Simulated phenotypes included main effects of gene expression (based on varying polygenicity), but did not include gene expression interaction effects. When using truly normally distributed gene expression values in the regression (top), the test statistic is well calibrated (i.e., Type I error rate ~alpha), regardless of whether additional variance is added and whether estimated (i.e., imperfectly predicted) expression data are used. However, when the true expression data is generated from binomially distributed SNPs, using an imperfectly predicted PGS results in inflation of the Type I error rate, proportional to how poorly the PGS predicts expression, i.e., with increasing error variance added to the predictor. Note that this does not occur when the true observed expression is used, even if binomially distributed. The effect is greatest for a PGS using a single SNP, and weakens as the expression becomes more polygenic.

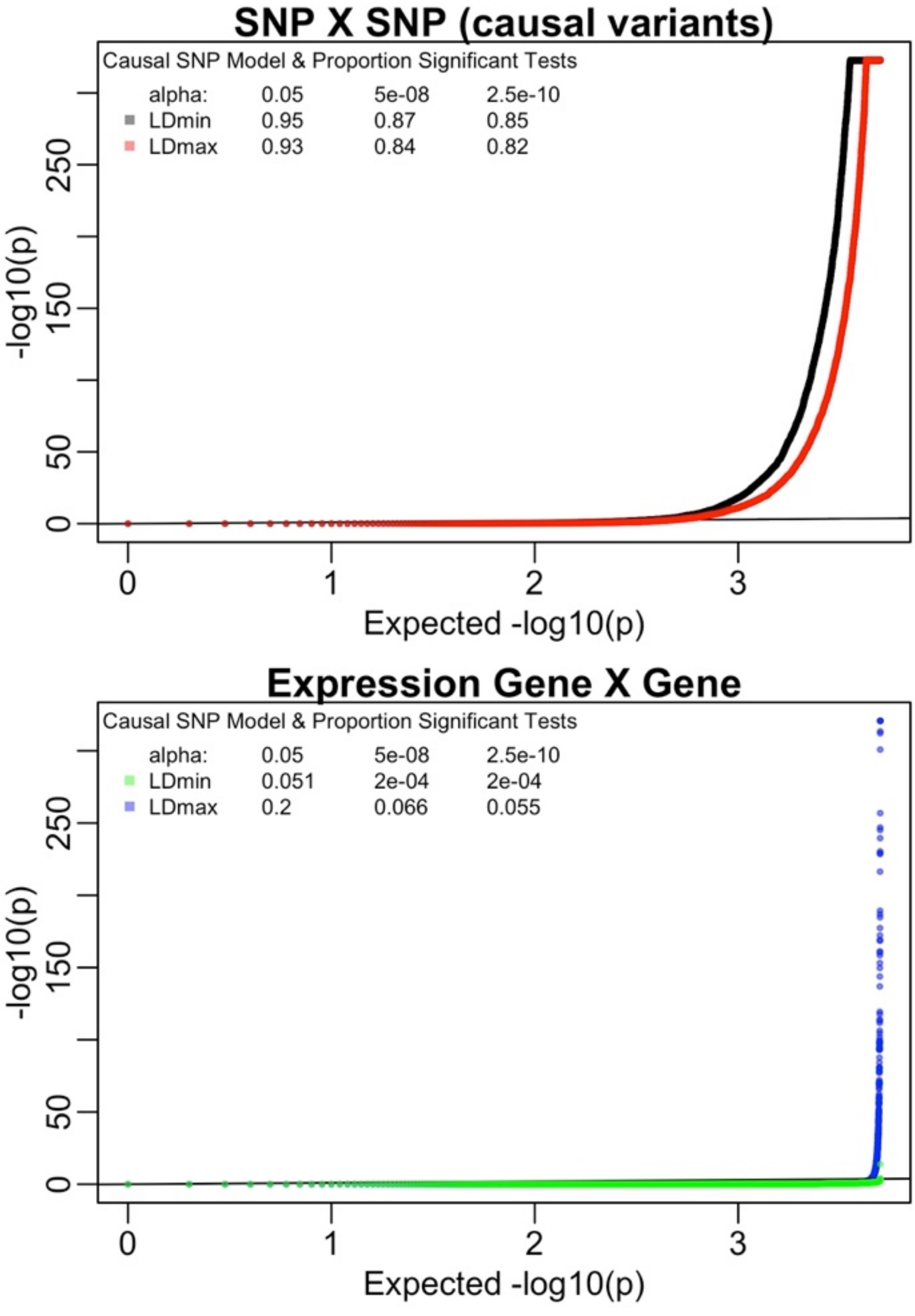


**Figure S6.** Simulations of causal SNPxSNP effects on the phenotype, tested either using SNPxSNP interactions (top) or imputed expression gene-gene interactions (bottom), when varying the LD (based on the LD score) of the causal SNPs. False positives increase when the SNPxSNP CVs have high LD scores than low LD scores, to the extent that the true effect is driven by SNP-SNP interactions, not expression-expression interactions. This results from LD between the causal SNPs and those used in the expression imputation.

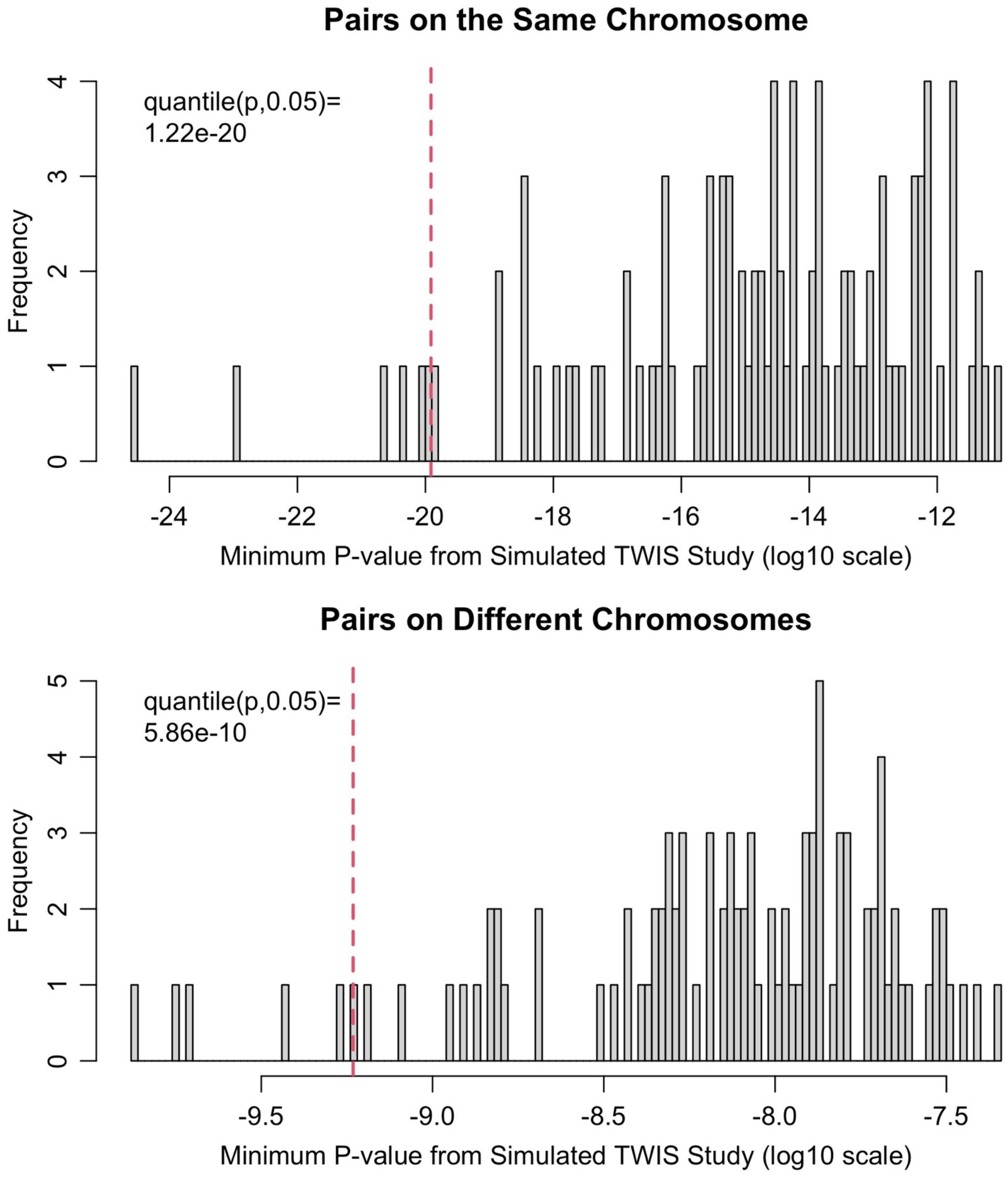


**Figure S7.** Distribution of the minimum interaction p-value for each of 100 simulated TWIS studies (each observation represents the minimum p-value of ~87M pairwise interaction tests across the genome), separated by whether the pair of genes is on the same (top) or different chromosomes (bottom). Red dashed line represents the 5th percentile of the minimum p-values. Note the x-axis scale differs between the two panels.

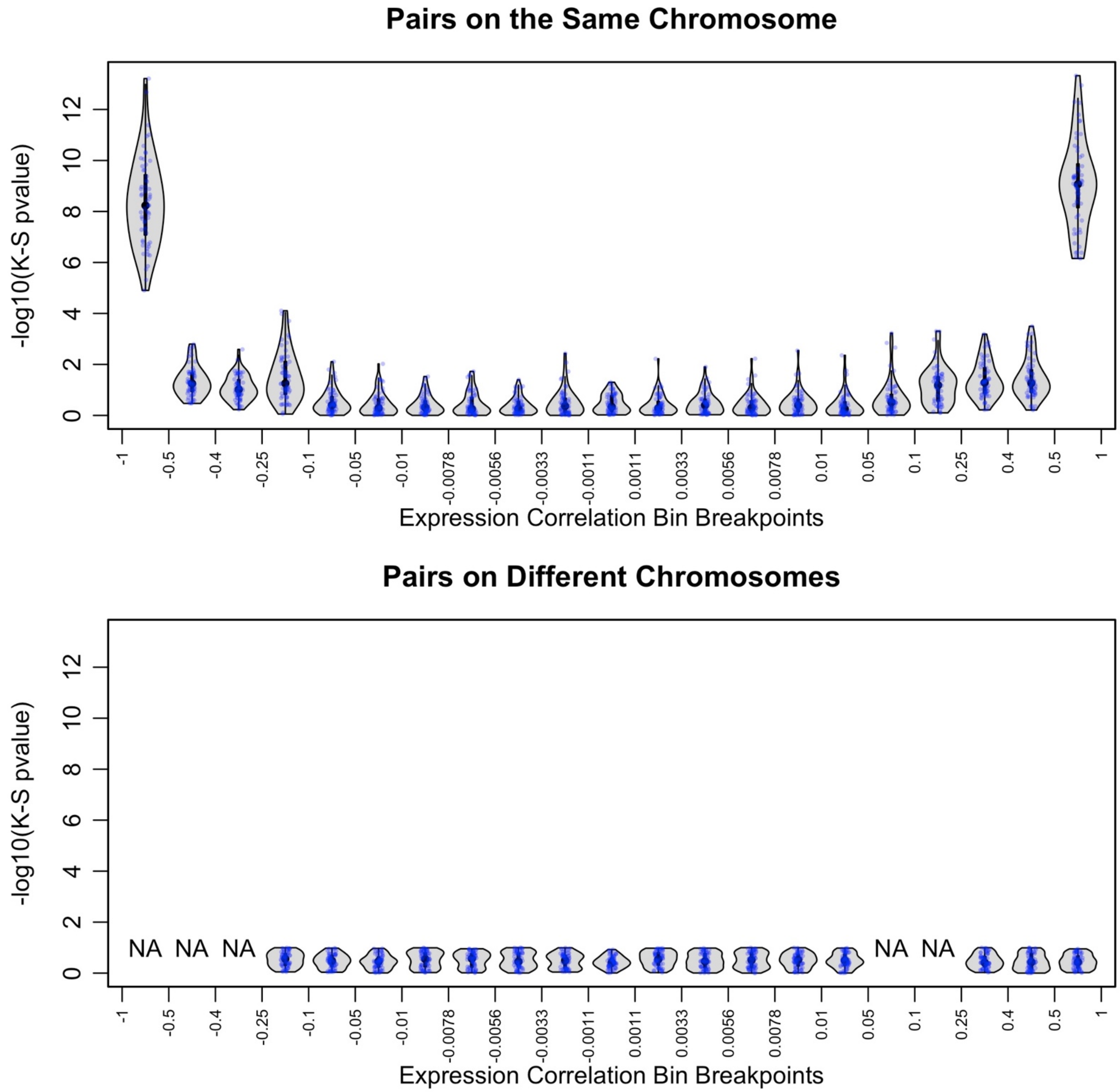


**Figure S8.** Violin plots of K-S test p-value testing whether the distribution of interaction test statistics is t-distributed across 40 whole genome TWIS study replications, depending on the pairwise imputed expression correlation between the gene pairs, and separated by whether the pair of genes is on the same (top) or different chromosomes (bottom). NA indicates no pairs of genes were found within that bin of pairwise imputed expression correlation. Blue dots are the (jittered) individual K-S test p-values for an entire simulated TWIS study.

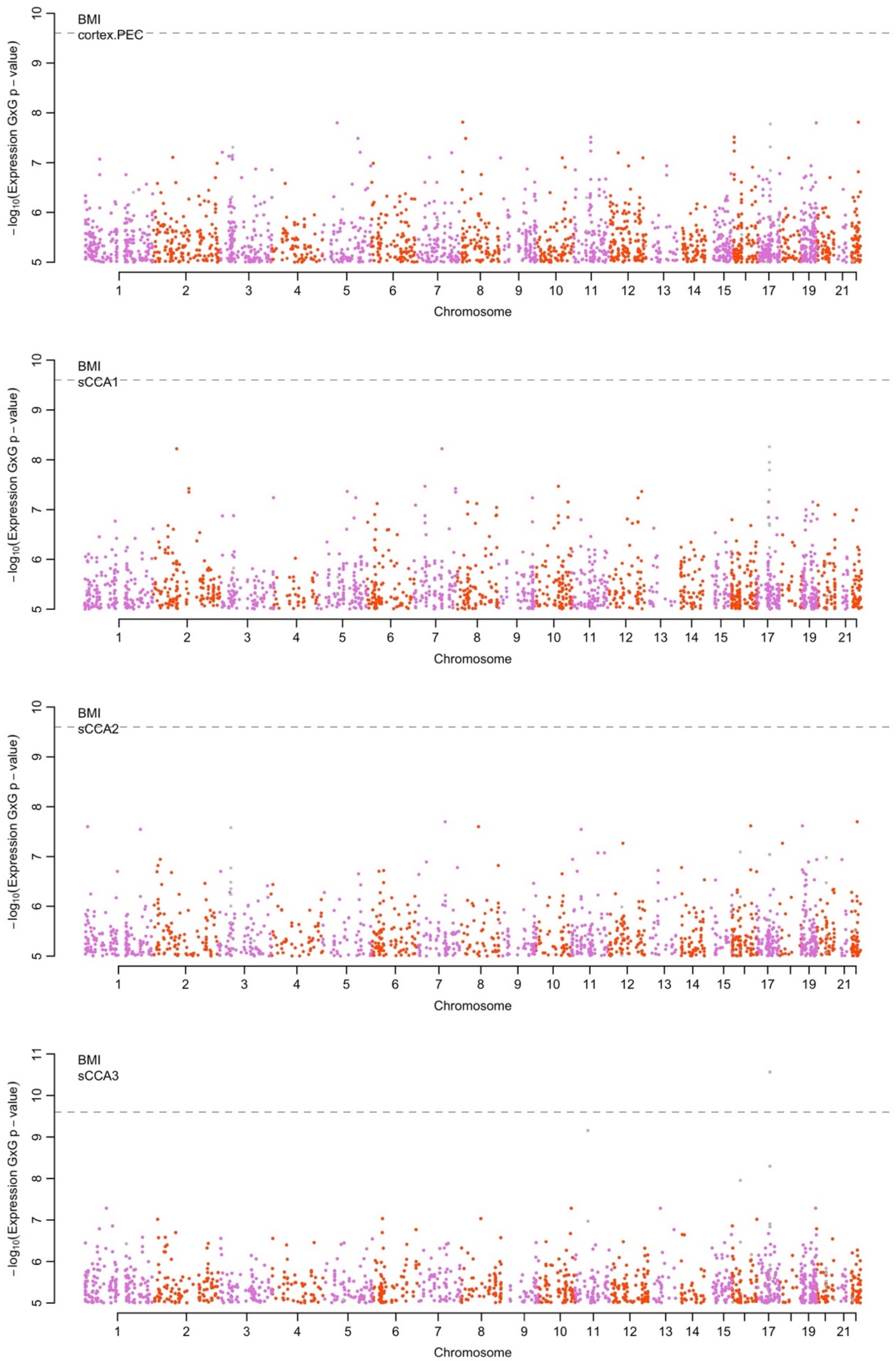


**Figure S9.** Boulder plot of BMI interaction association p-values using imputed transcription. Shown are the results from the final meta-analysis of all data. Black lines connect pairs that surpassed p<2.5e-10 in the discovery cohort (UKB), blue lines connect pairs of loci with nominally significant interaction (p<0.05) in the replication cohort, and gray lines connect pairs of genes with p<2.5e-10 in the final meta-analysis.

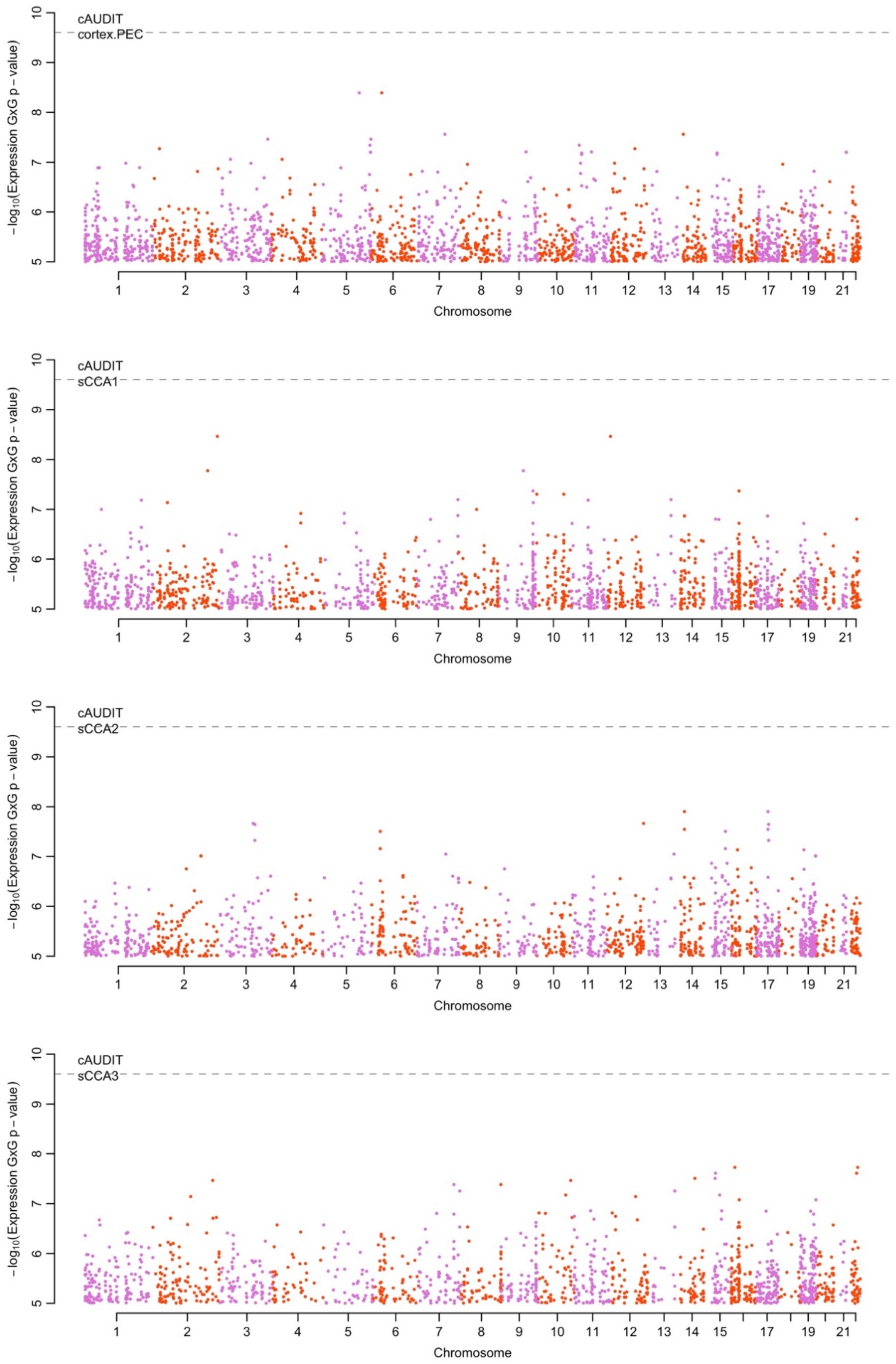


**Figure S10.** Boulder plot of cAUDIT interaction association p-values using imputed transcription. Shown are the results from the final meta-analysis of all data. Black lines connect pairs that surpassed p<2.5e-10 in the discovery cohort (UKB), blue lines connect pairs of loci with nominally significant interaction (p<0.05) in the replication cohort, and gray lines connect pairs of genes with p<2.5e-10 in the final meta-analysis.

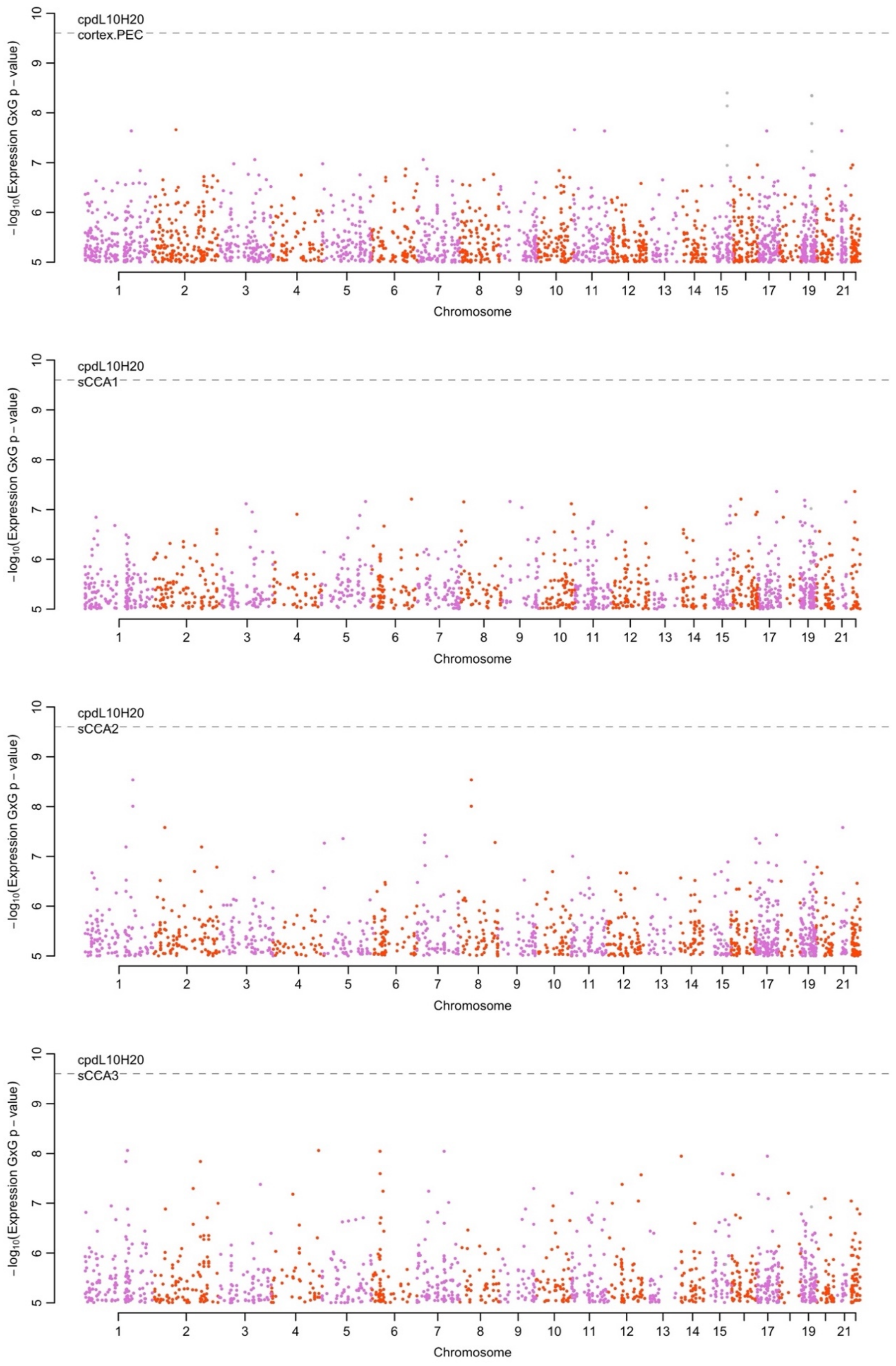


**Figure S11.** Boulder plot of CPD (heavy CPD>=20 vs light CPD<=10) interaction association p-values using imputed transcription. Shown are the results from the final meta-analysis of all data. Black lines connect pairs that surpassed p<2.5e-10 in the discovery cohort (UKB), blue lines connect pairs of loci with nominally significant interaction (p<0.05) in the replication cohort, and gray lines connect pairs of genes with p<2.5e-10 in the final meta-analysis.

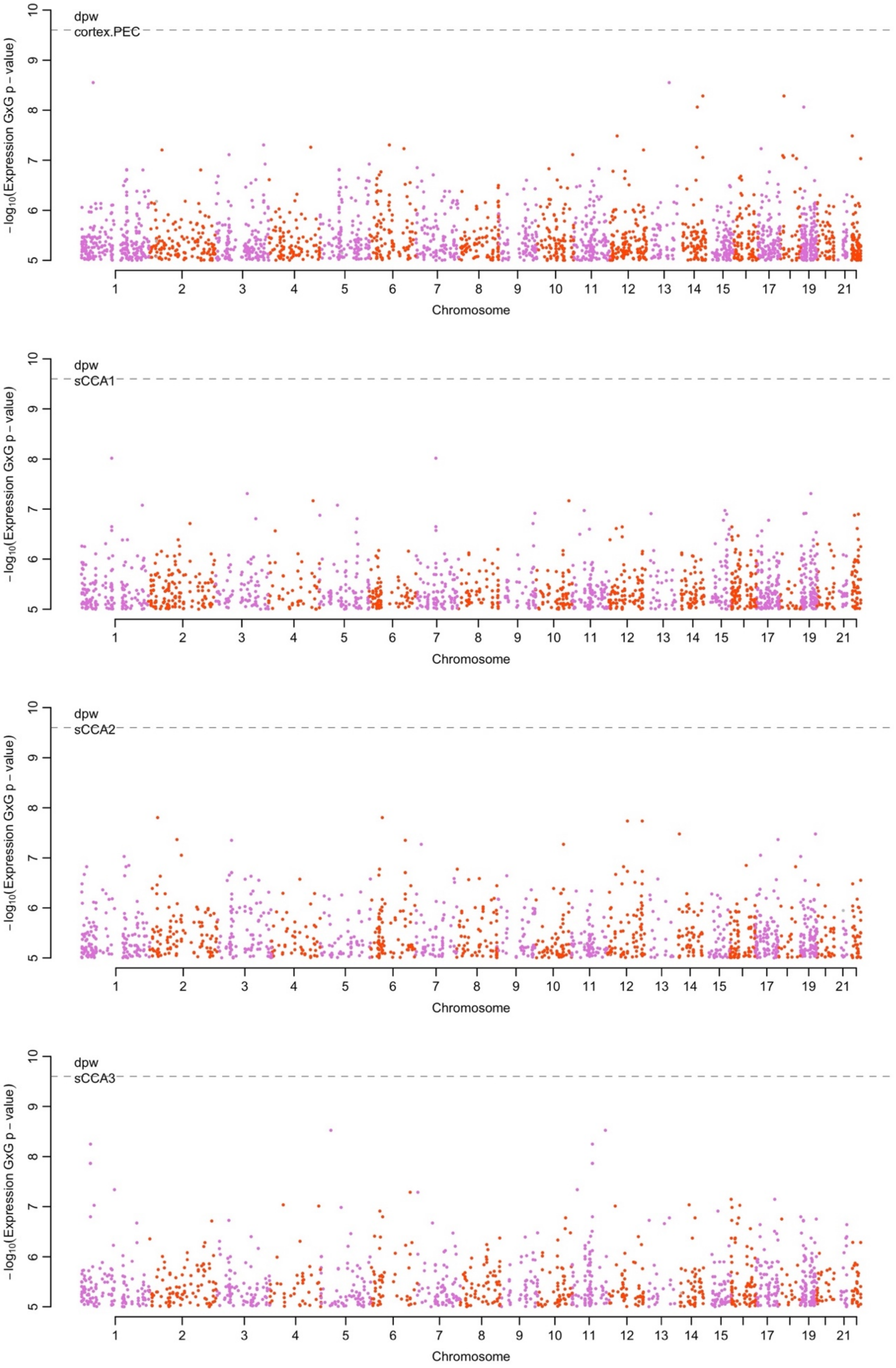


**Figure S12.** Boulder plot of DPW interaction association p-values using imputed transcription. Shown are the results from the final meta-analysis of all data. Black lines connect pairs that surpassed p<2.5e-10 in the discovery cohort (UKB), blue lines connect pairs of loci with nominally significant interaction (p<0.05) in the replication cohort, and gray lines connect pairs of genes with p<2.5e-10 in the final meta-analysis.

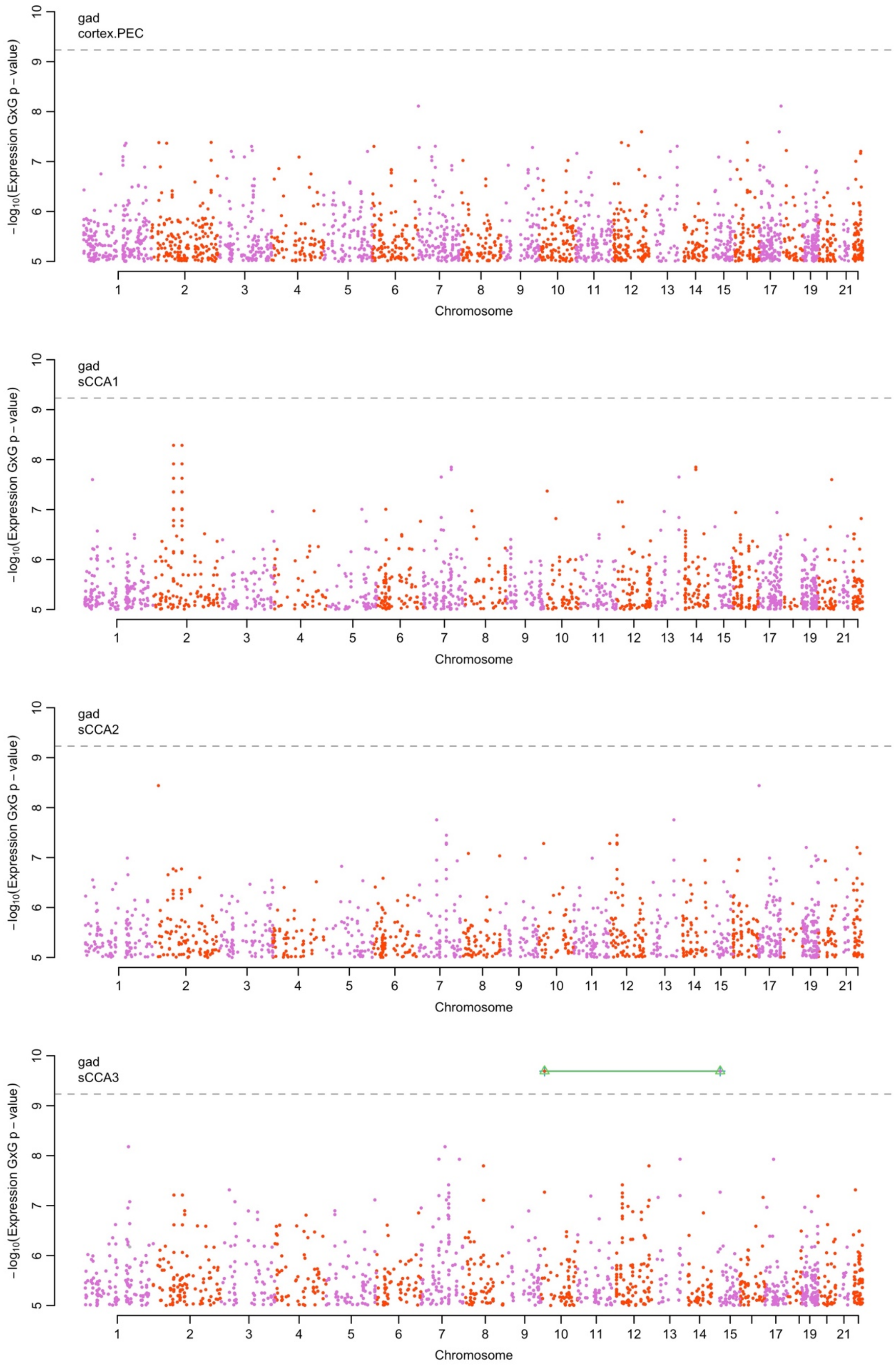


**Figure S13.** Boulder plot of GAD interaction association p-values using imputed transcription. Shown are the results from the final meta-analysis of all data. Black lines connect pairs that surpassed p<2.5e-10 in the discovery cohort (UKB), green lines connect pairs of loci with significant (q<0.05) in the replication cohort, and gray lines connect pairs of genes with p<2.5e-10 in the final meta-analysis.

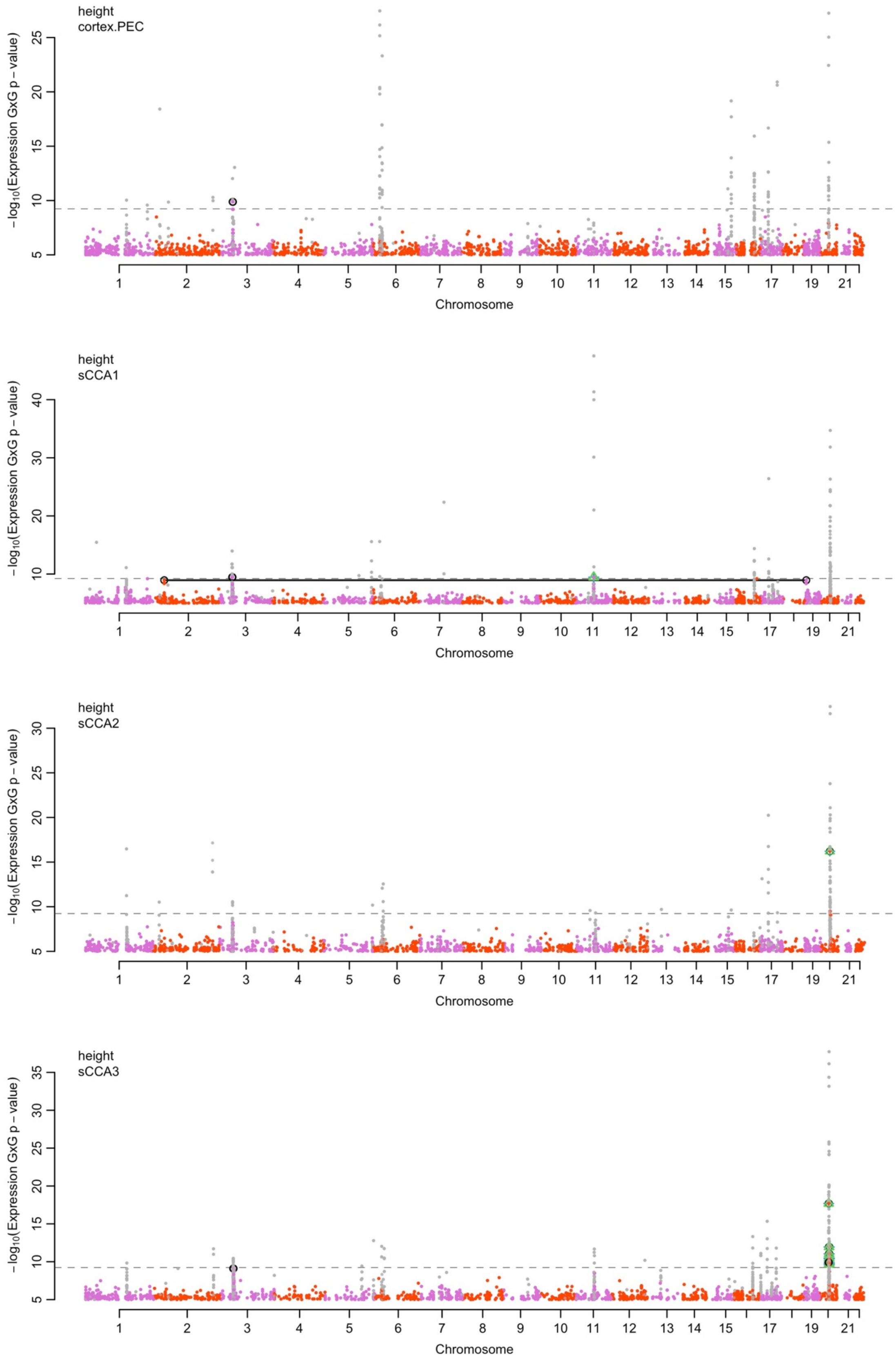


**Figure S14.** Boulder plot of height interaction association p-values using imputed transcription. Shown are the results from the final meta-analysis of all data. Black lines connect pairs that surpassed p<2.5e-10 in the discovery cohort (UKB), blue lines connect pairs of loci with nominally significant interaction (p<0.05) in the replication cohort, and gray lines connect pairs of genes with p<2.5e-10 in the final meta-analysis.

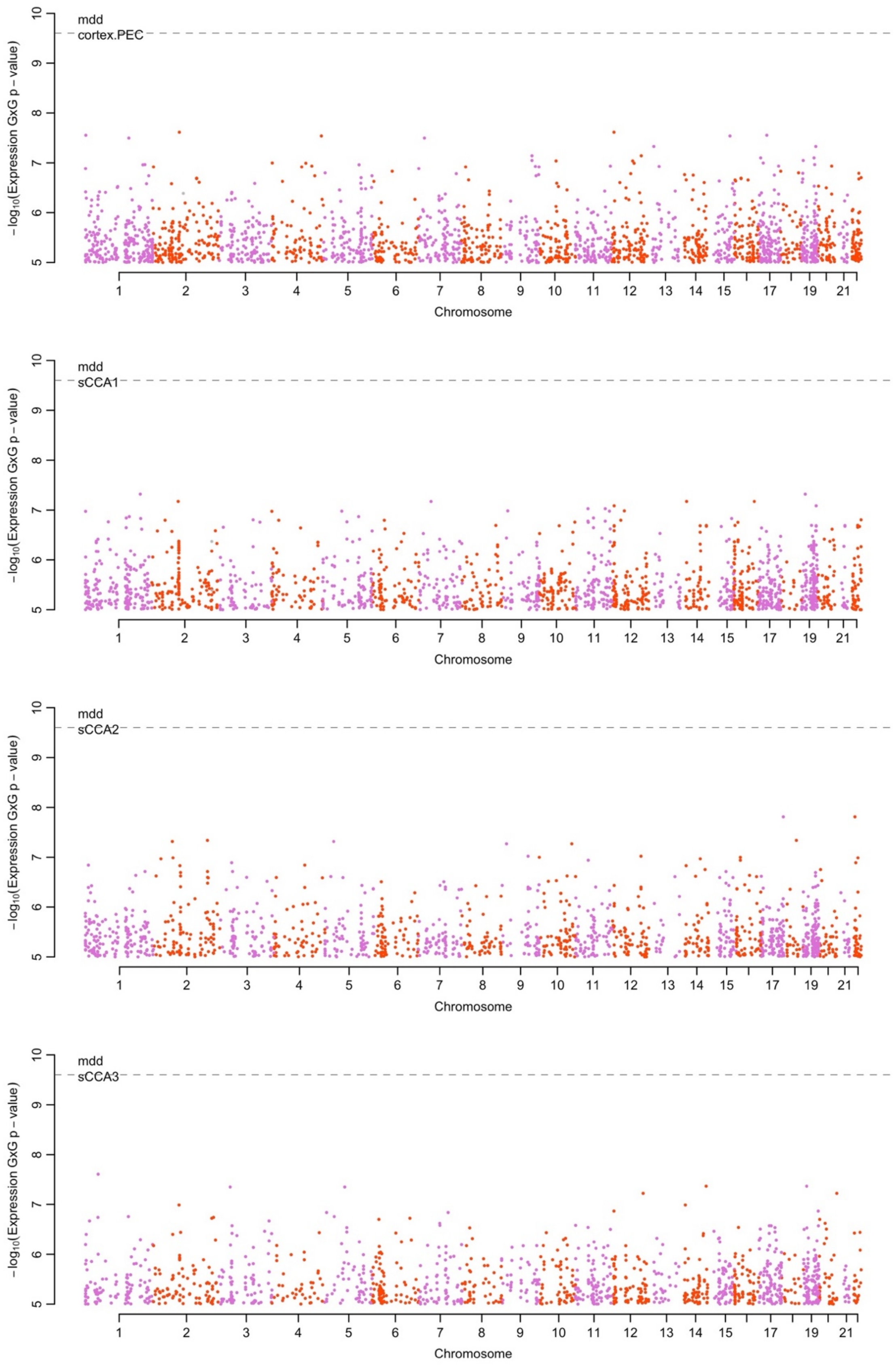


**Figure S15.** Boulder plot of MDD interaction association p-values using imputed transcription. Shown are the results from the final meta-analysis of all data. Black lines connect pairs that surpassed p<2.5e-10 in the discovery cohort (UKB), blue lines connect pairs of loci with nominally significant interaction (p<0.05) in the replication cohort, and gray lines connect pairs of genes with p<2.5e-10 in the final meta-analysis.

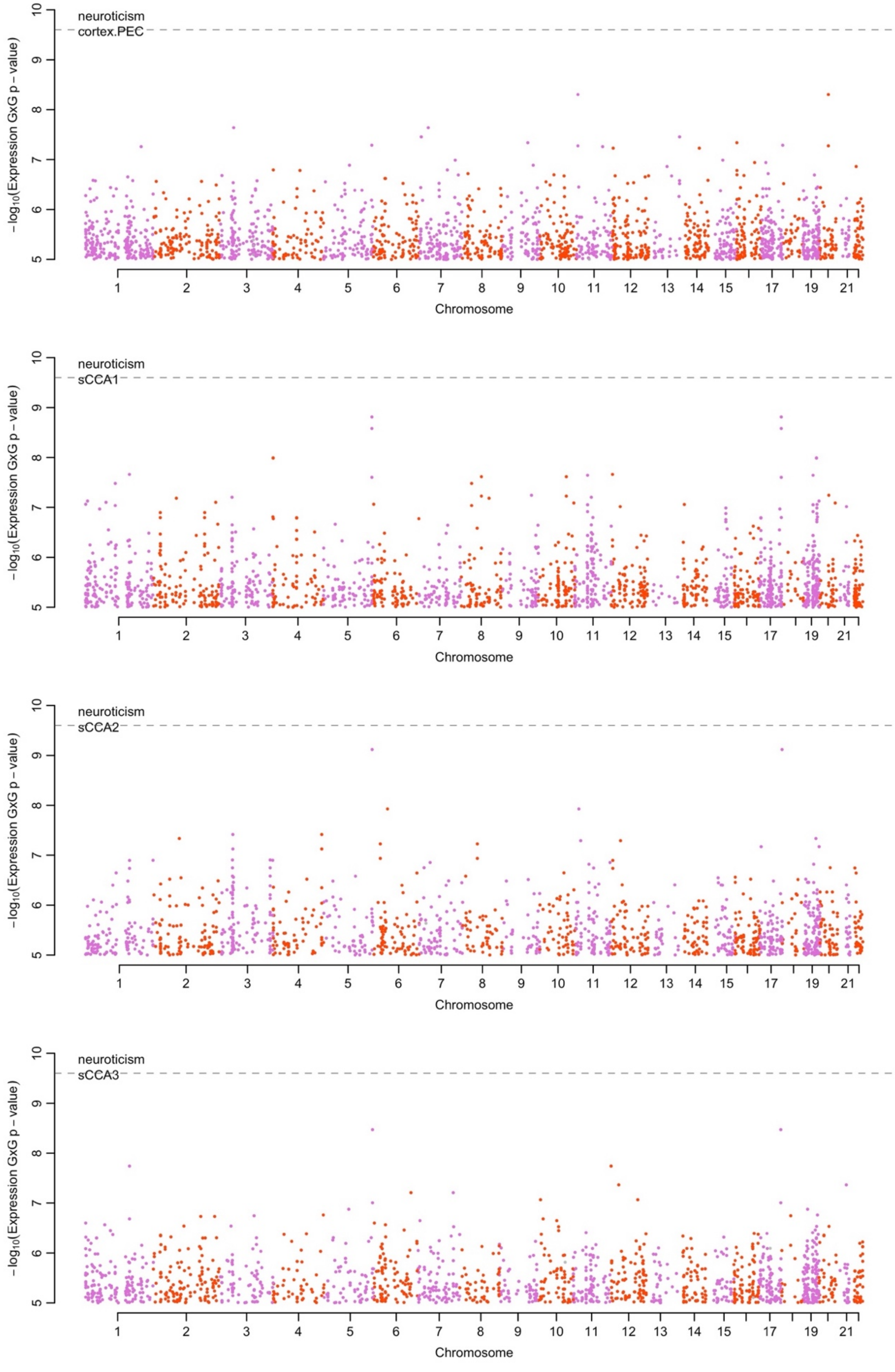


**Figure S16.** Boulder plot of neuroticism interaction association p-values using imputed transcription. Shown are the results from the final meta-analysis of all data. Black lines connect pairs that surpassed p<2.5e-10 in the discovery cohort (UKB), blue lines connect pairs of loci with nominally significant interaction (p<0.05) in the replication cohort, and gray lines connect pairs of genes with p<2.5e-10 in the final meta-analysis.

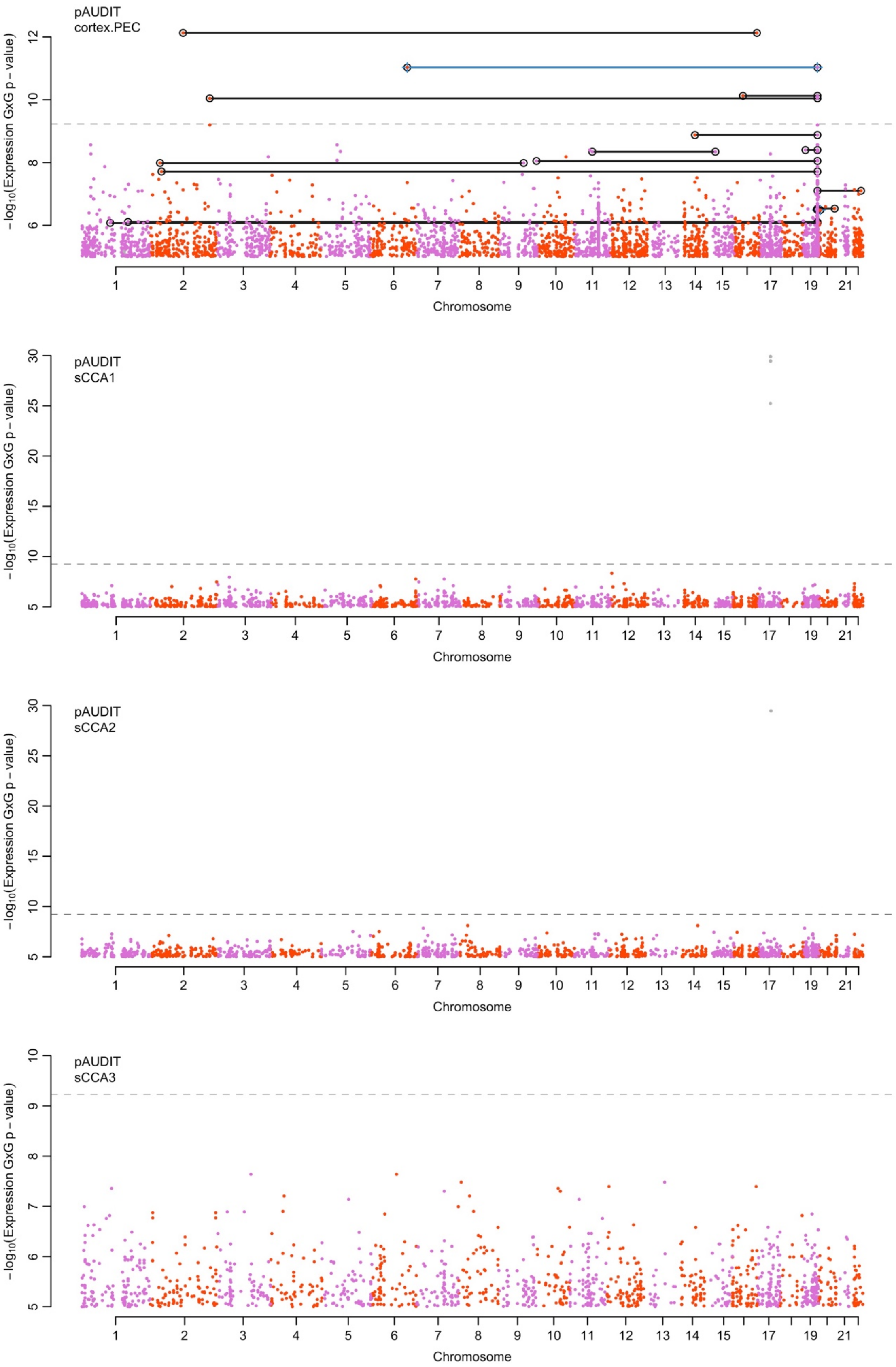


**Figure S17.** Boulder plot of pAUDIT interaction association p-values using imputed transcription. Shown are the results from the final meta-analysis of all data. Black lines connect pairs that surpassed p<2.5e-10 in the discovery cohort (UKB), blue lines connect pairs of loci with nominally significant interaction (p<0.05) in the replication cohort, and gray lines connect pairs of genes with p<2.5e-10 in the final meta-analysis.

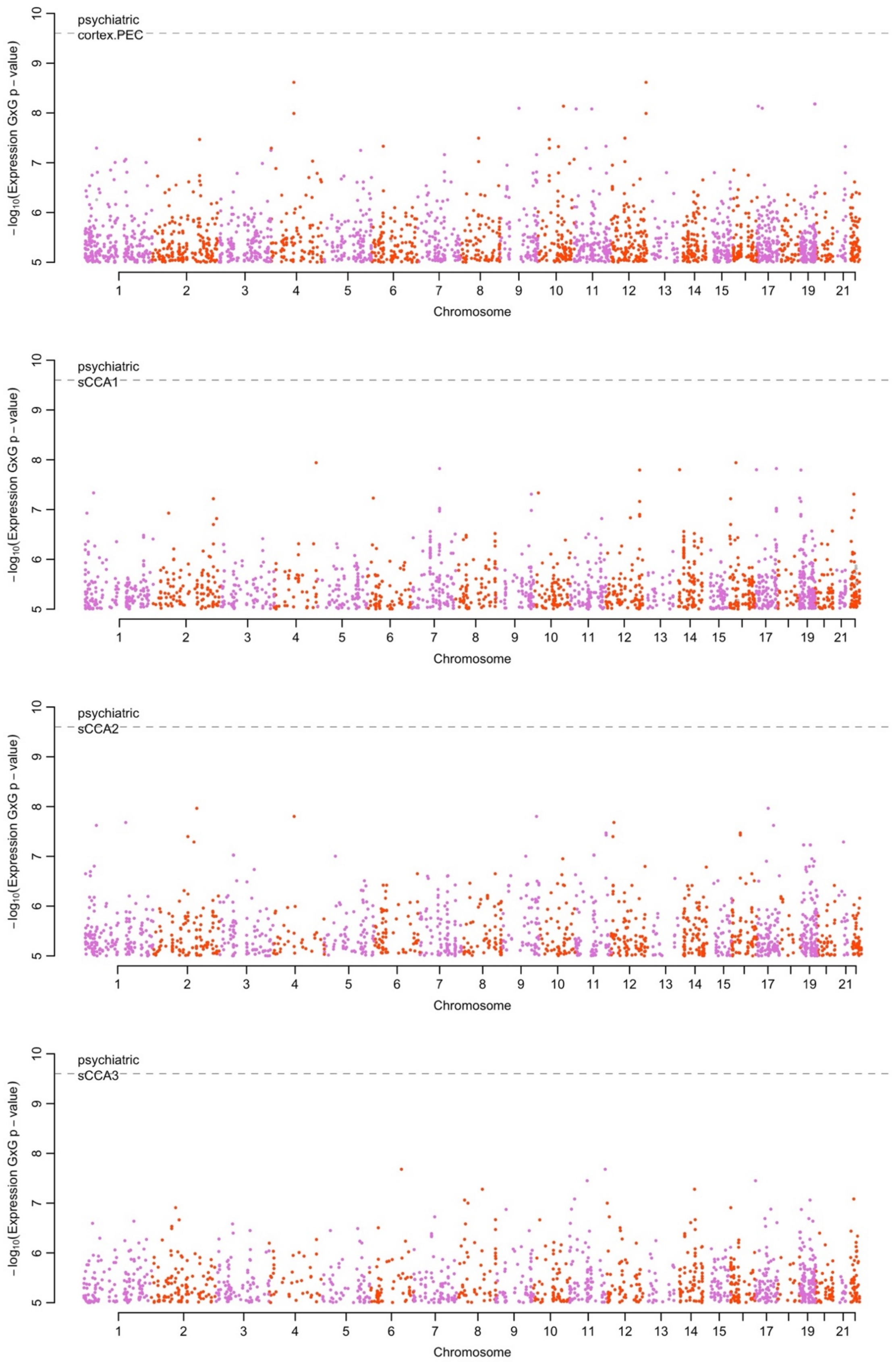


**Figure S18.** Boulder plot of psychiatric interaction association p-values using imputed transcription. Shown are the results from the final meta-analysis of all data. Black lines connect pairs that surpassed p<2.5e-10 in the discovery cohort (UKB), blue lines connect pairs of loci with nominally significant interaction (p<0.05) in the replication cohort, and gray lines connect pairs of genes with p<2.5e-10 in the final meta-analysis.

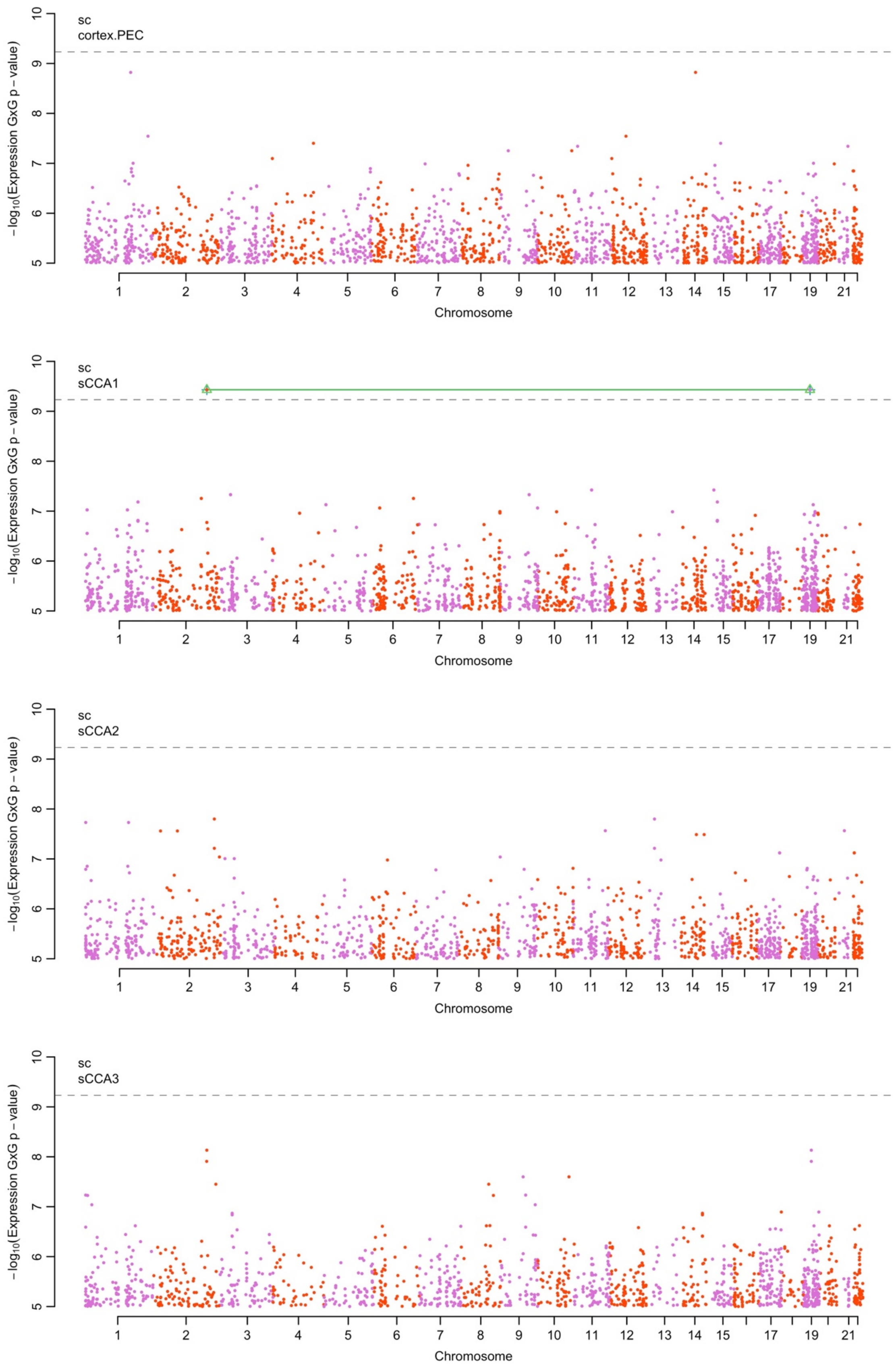


**Figure S19.** Boulder plot of smoking cessation (SC) interaction association p-values using imputed transcription. Shown are the results from the final meta-analysis of all data. Black lines connect pairs that surpassed p<2.5e-10 in the discovery cohort (UKB), blue lines connect pairs of loci with nominally significant interaction (p<0.05) in the replication cohort, and gray lines connect pairs of genes with p<2.5e-10 in the final meta-analysis.

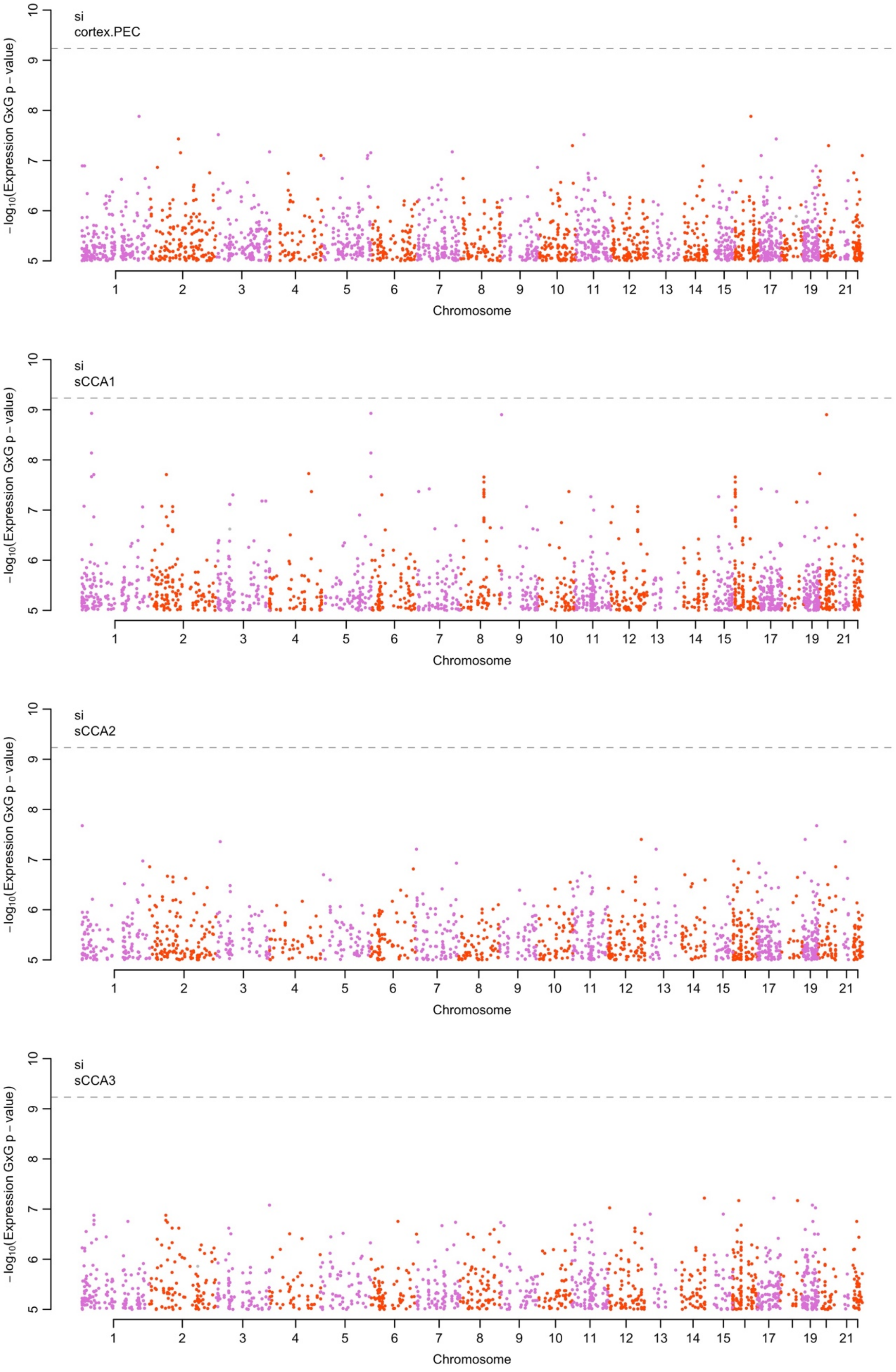


**Figure S20.** Boulder plot of smoking interaction (SI) interaction association p-values using imputed transcription. Shown are the results from the final meta-analysis of all data. Black lines connect pairs that surpassed p<2.5e-10 in the discovery cohort (UKB), blue lines connect pairs of loci with nominally significant interaction (p<0.05) in the replication cohort, and gray lines connect pairs of genes with p<2.5e-10 in the final meta-analysis.

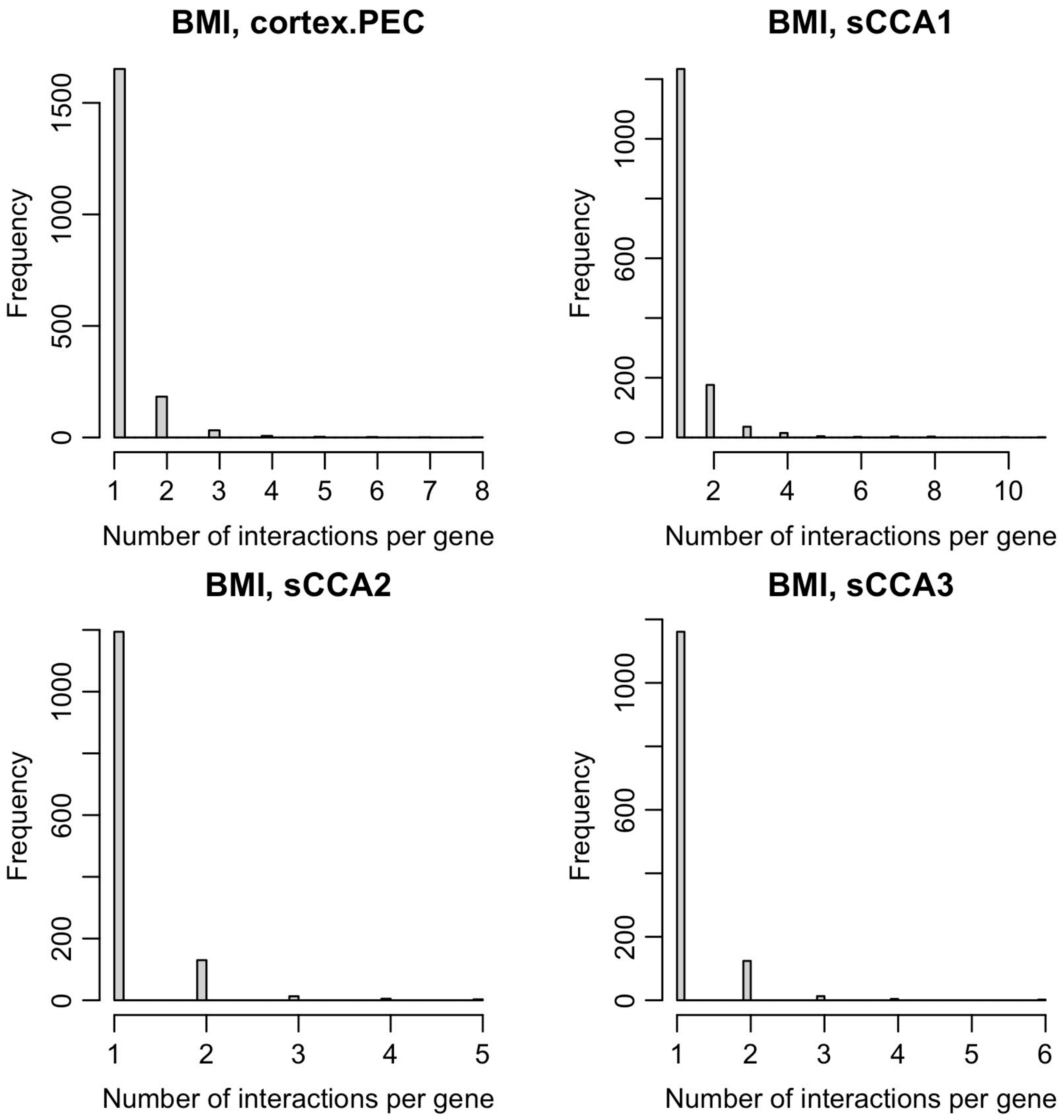


**Figure S21.** Distribution of number of interaction associations for each gene with at least one suggestive (p<1e-5) interaction with BMI.

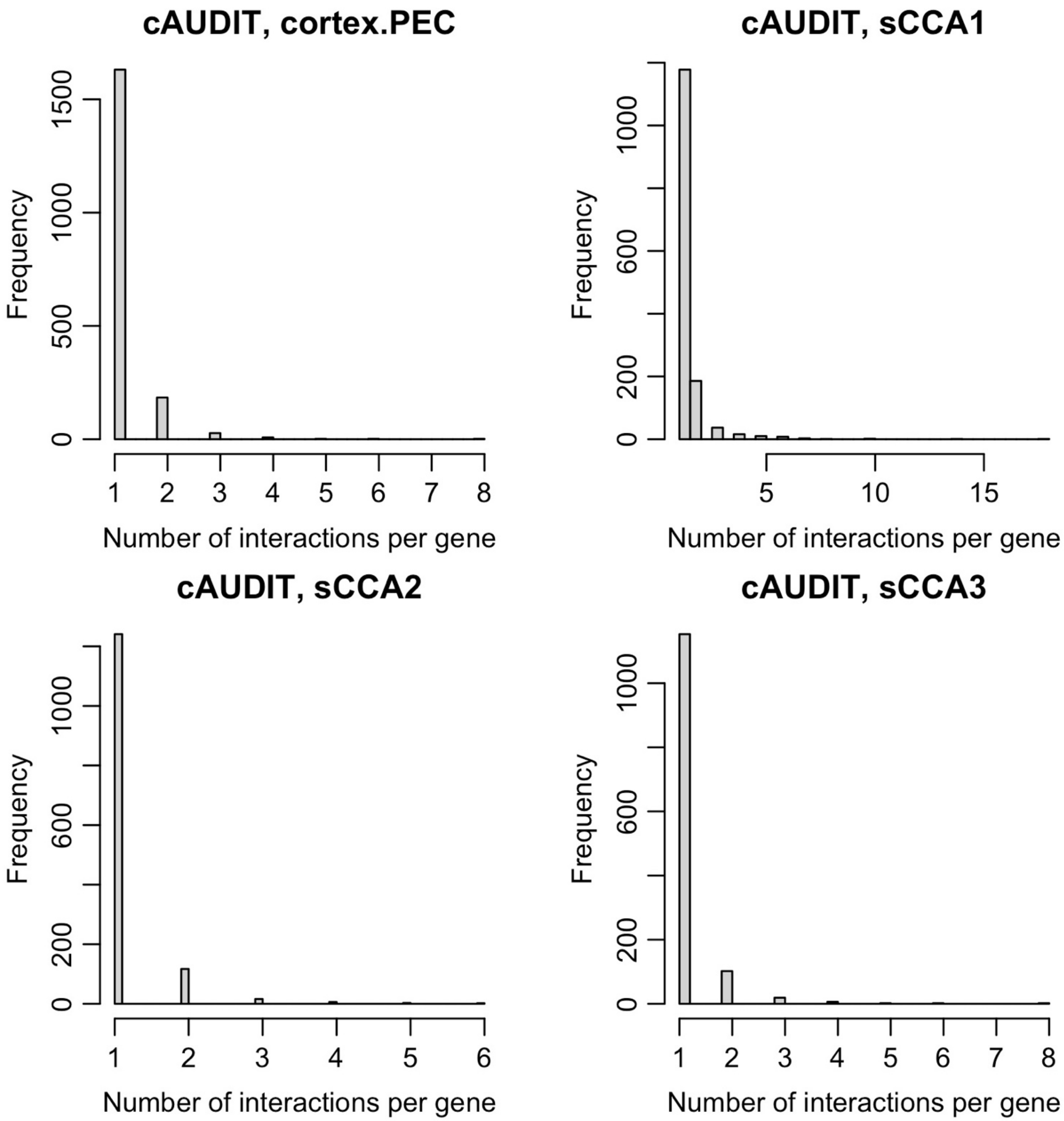


**Figure S22.** Distribution of number of interaction associations for each gene with at least one suggestive (p<1e-5) interaction with cAUDIT.

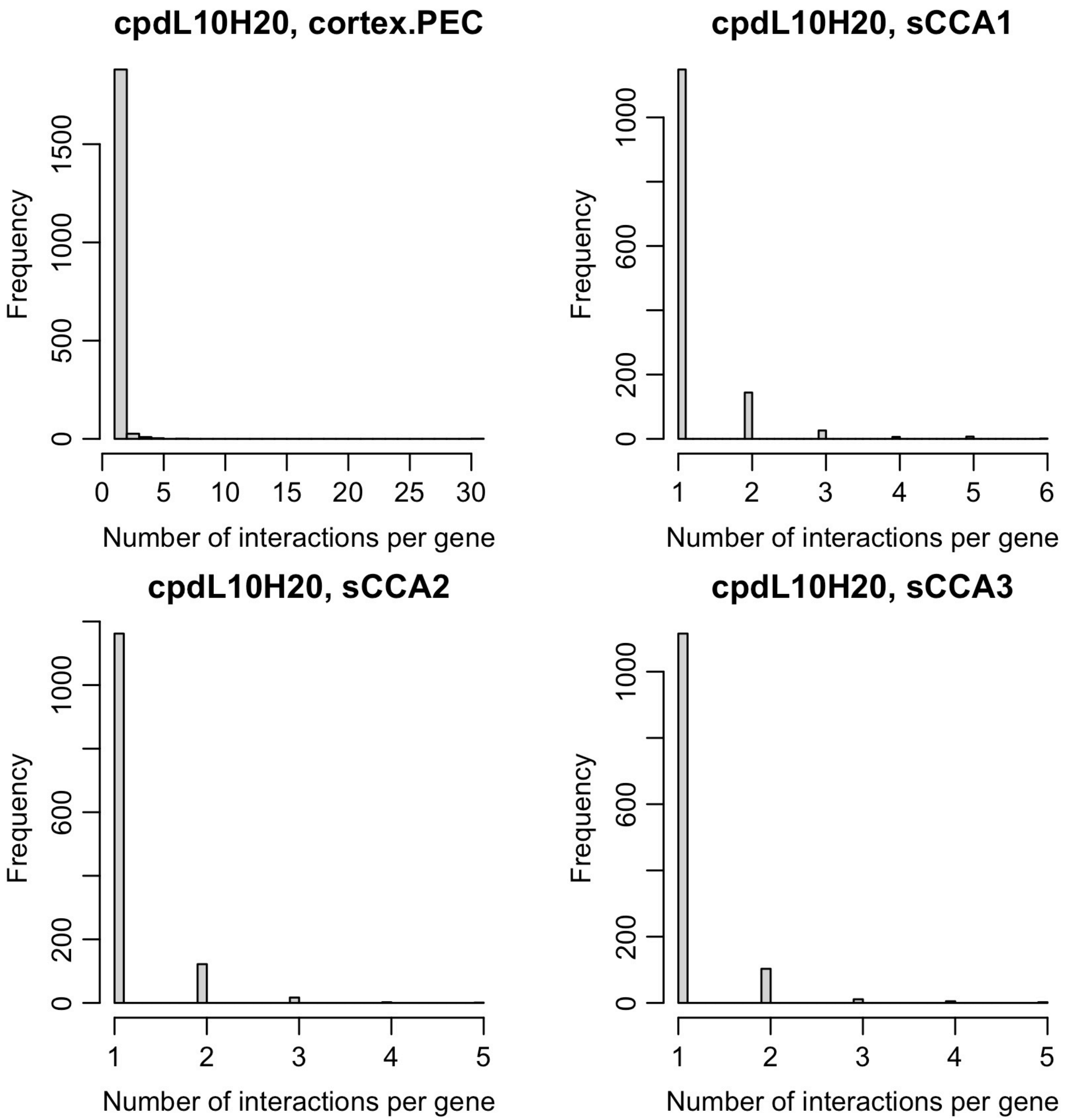


**Figure S23.** Distribution of number of interaction associations for each gene with at least one suggestive (p<1e-5) interaction with CPD (high vs. low use).

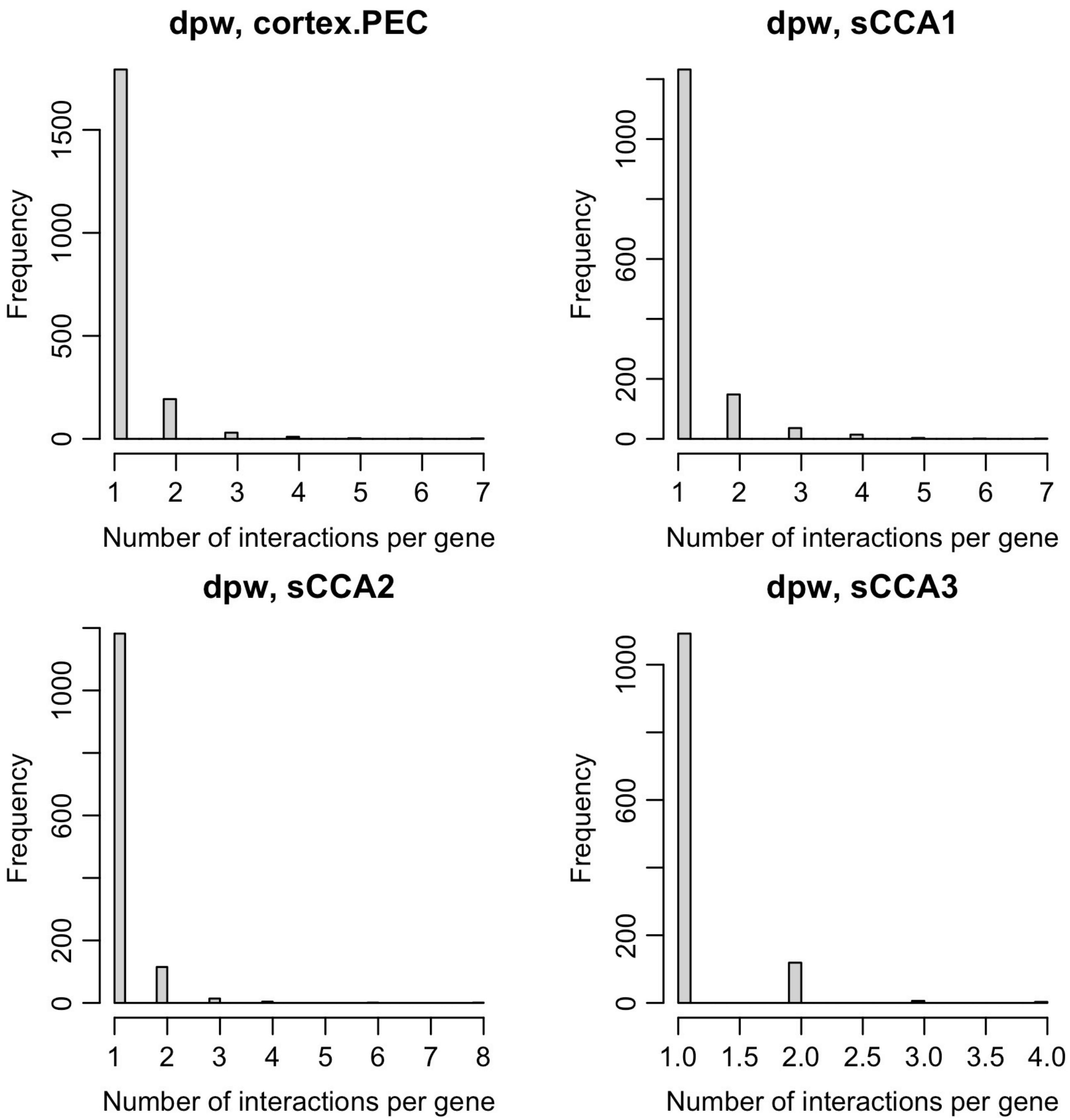


**Figure S24.** Distribution of number of interaction associations for each gene with at least one suggestive (p<1e-5) interaction with DPW.

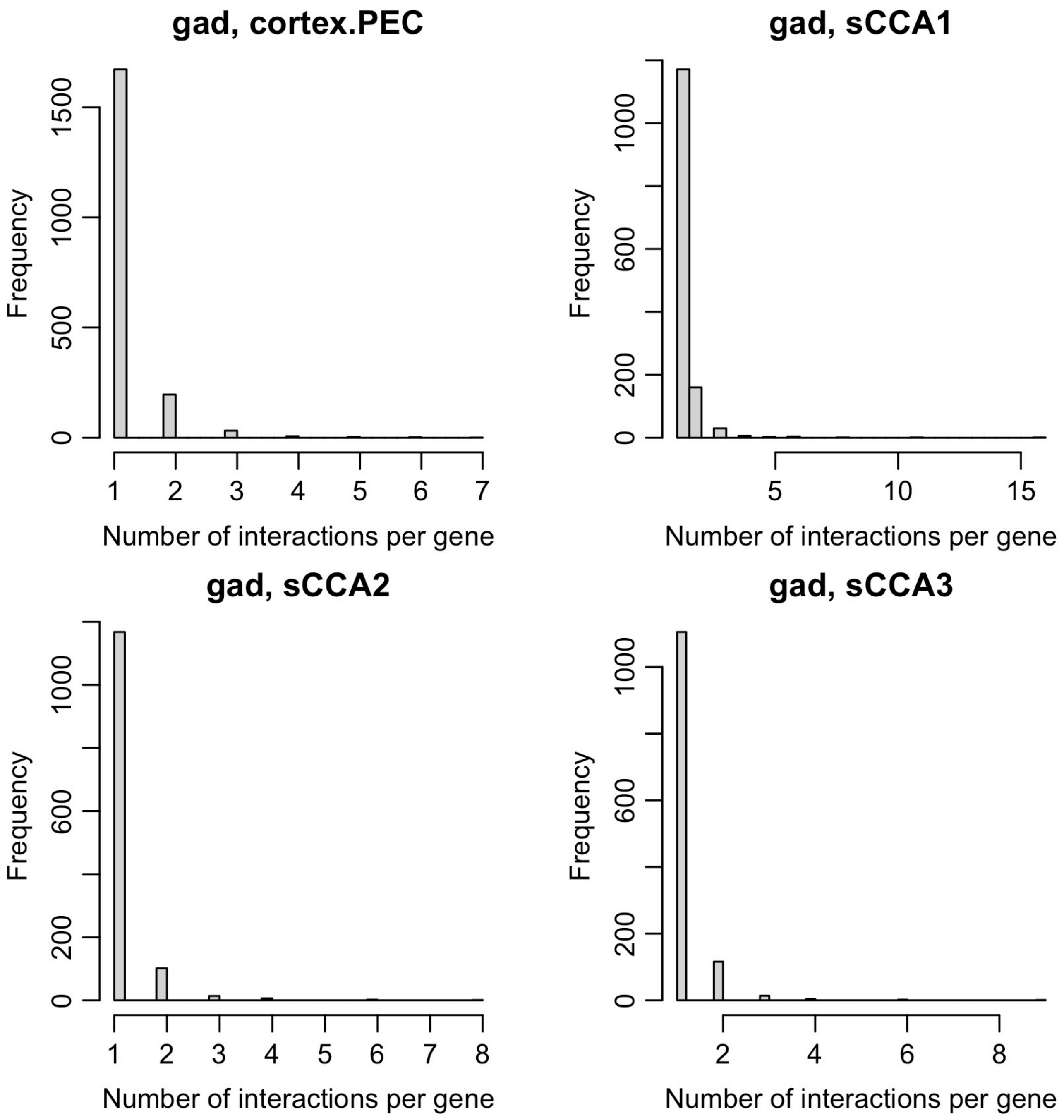


**Figure S25.** Distribution of number of interaction associations for each gene with at least one suggestive (p<1e-5) interaction with GAD.

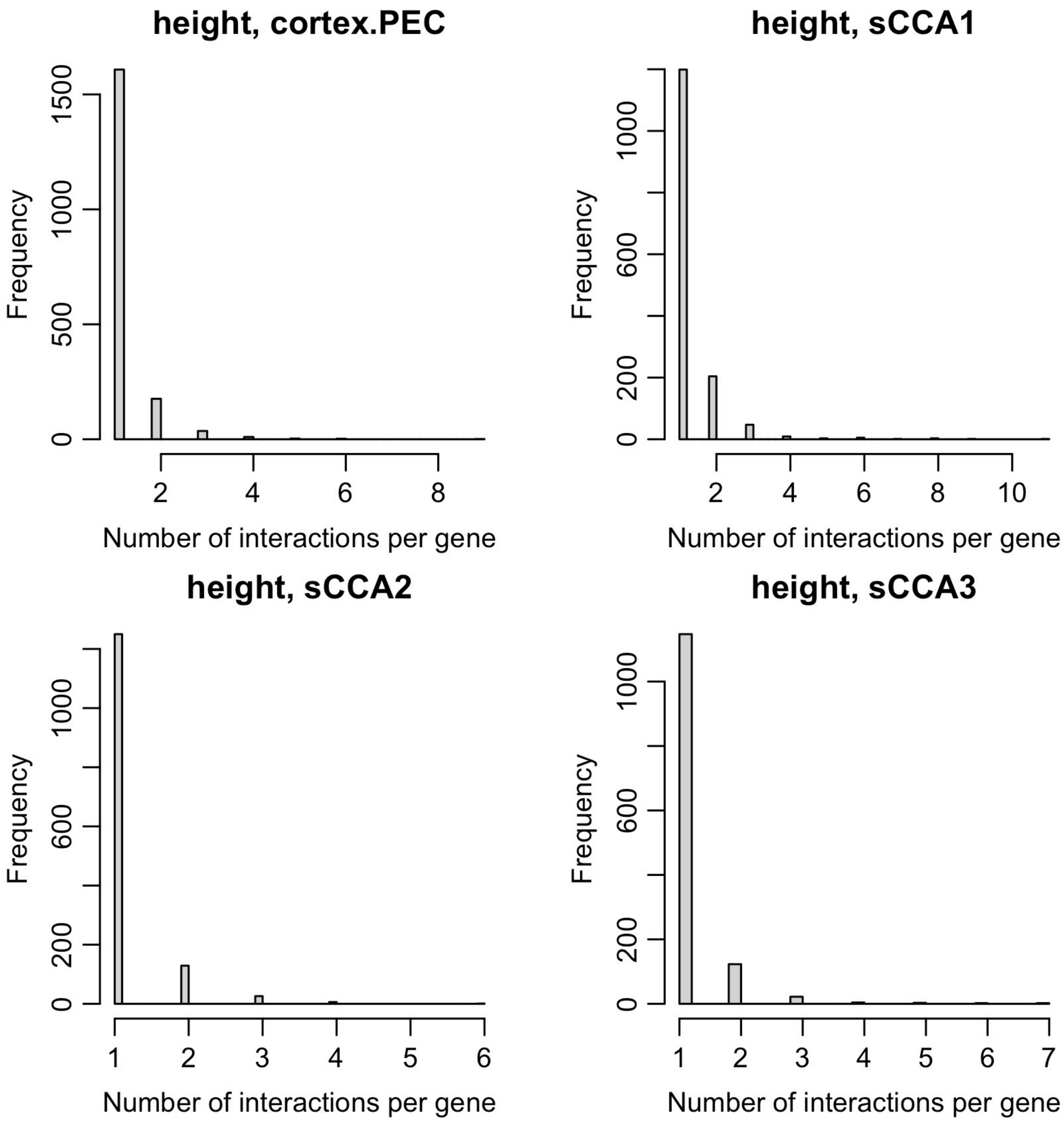


**Figure S26.** Distribution of number of interaction associations for each gene with at least one suggestive (p<1e-5) interaction with height.

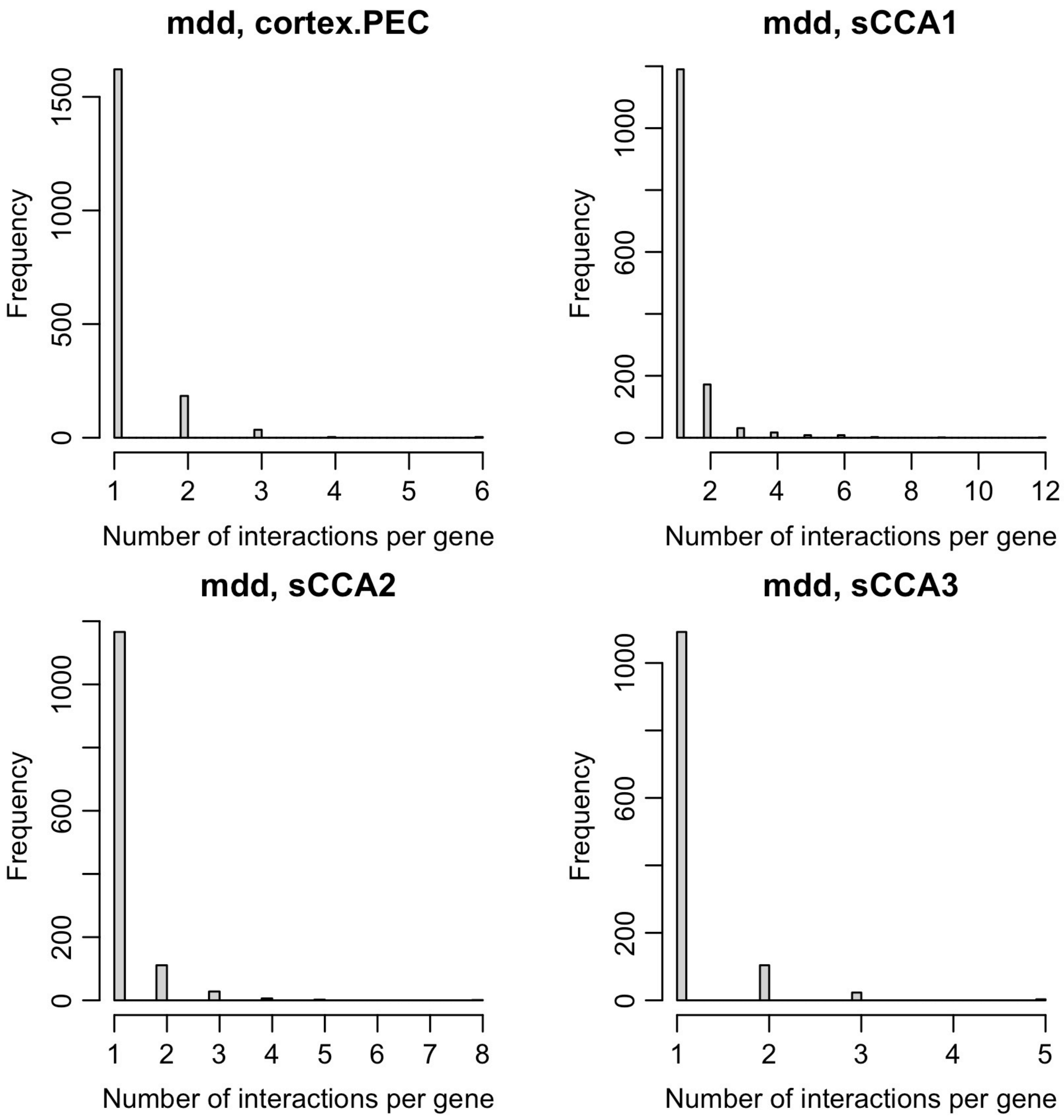


**Figure S27.** Distribution of number of interaction associations for each gene with at least one suggestive (p<1e-5) interaction with MDD

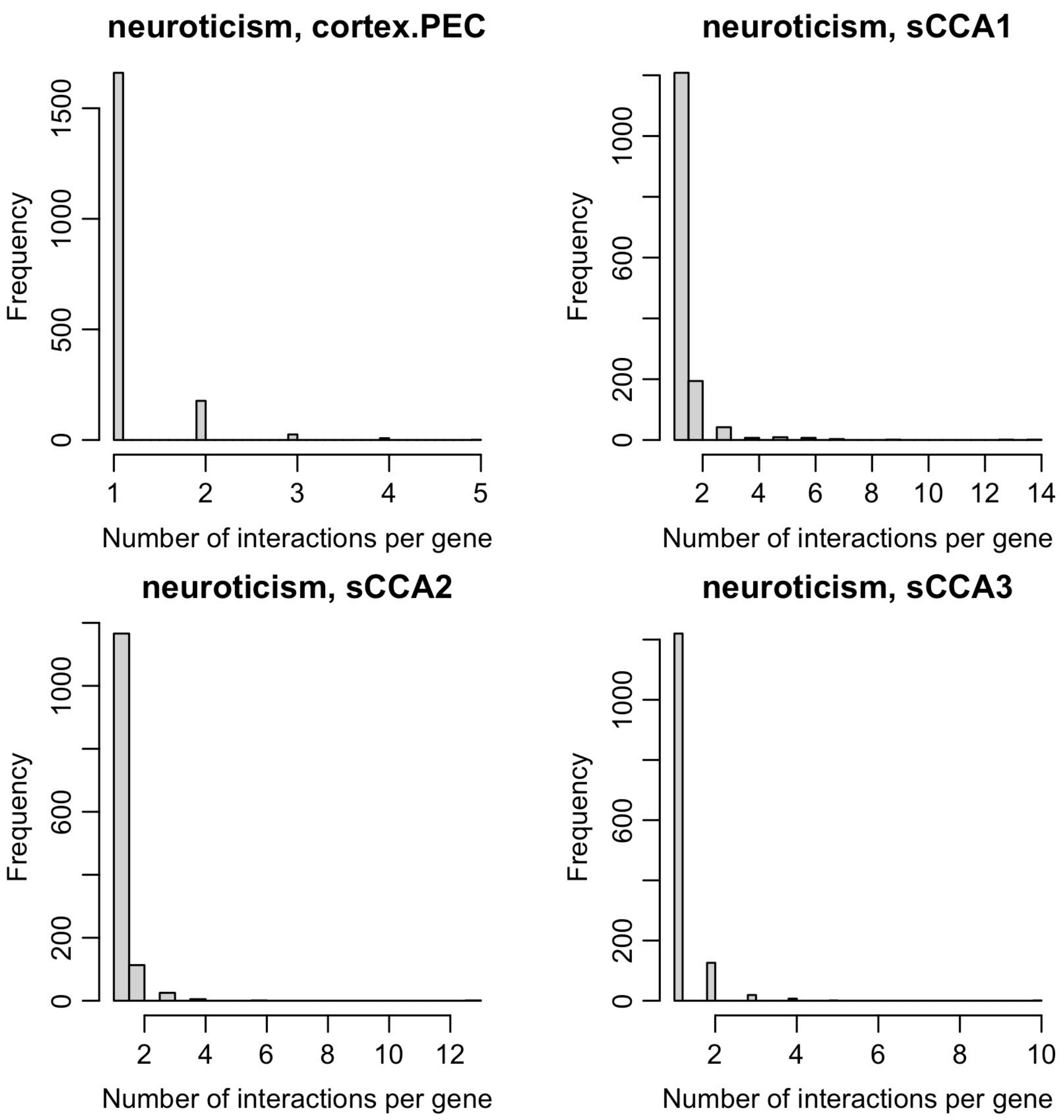


**Figure S28.** Distribution of number of interaction associations for each gene with at least one suggestive (p<1e-5) interaction with neuroticism.

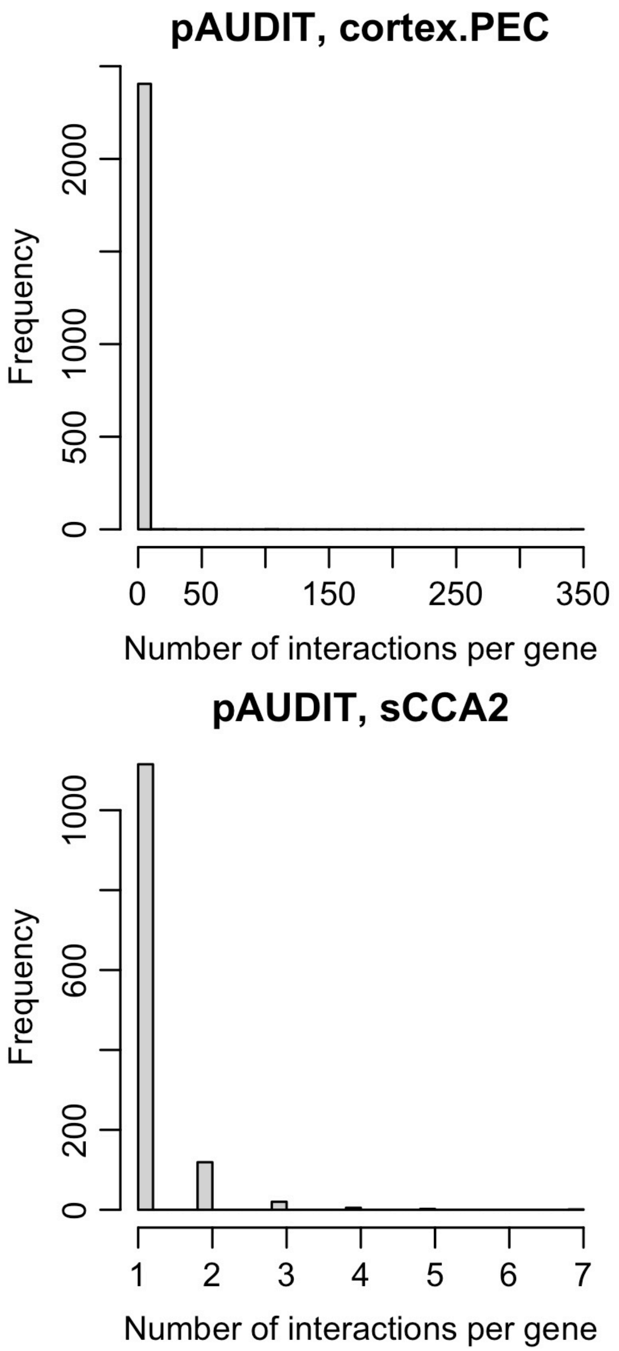


**Figure S29.** Distribution of number of interaction associations for each gene with at least one suggestive (p<1e-5) interaction with pAUDIT.

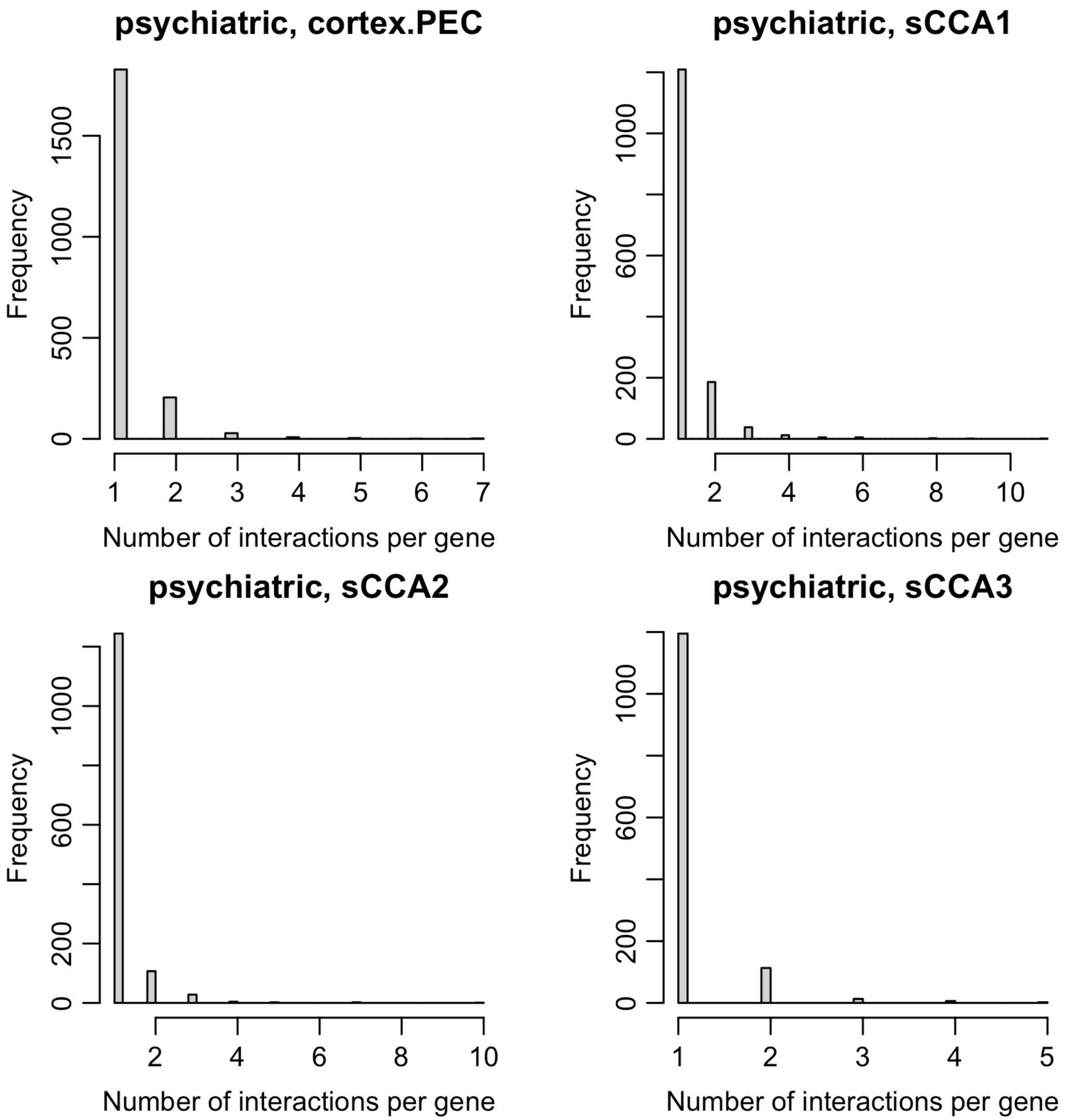


**Figure S30.** Distribution of number of interaction associations for each gene with at least one suggestive (p<1e-5) interaction with psychiatric.

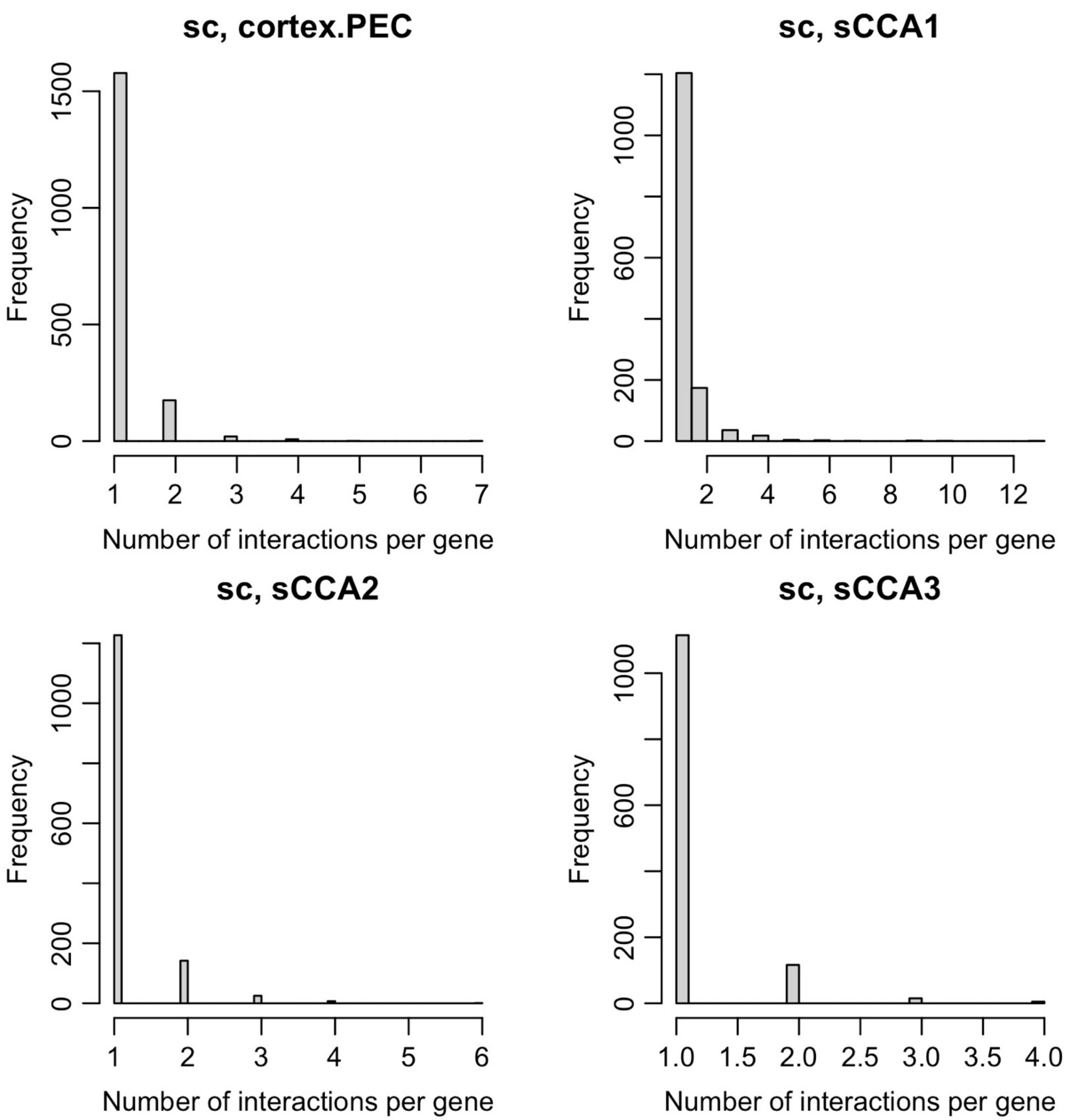


**Figure S31.** Distribution of number of interaction associations for each gene with at least one suggestive (p<1e-5) interaction with smoking cessation (SC).

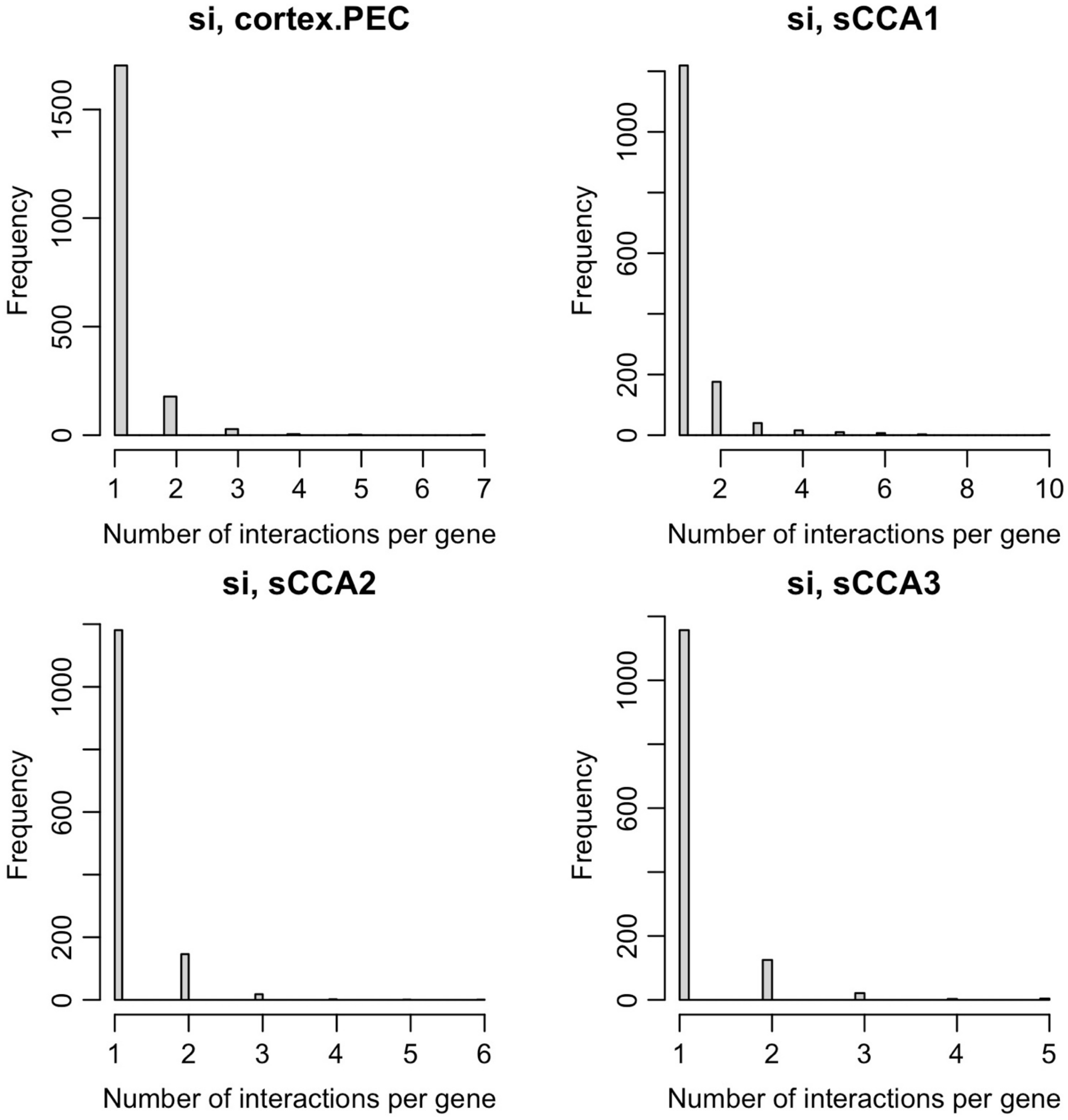


**Figure S32.** Distribution of number of interaction associations for each gene with at least one suggestive (p<1e-5) interaction with smoking initiation (SI).

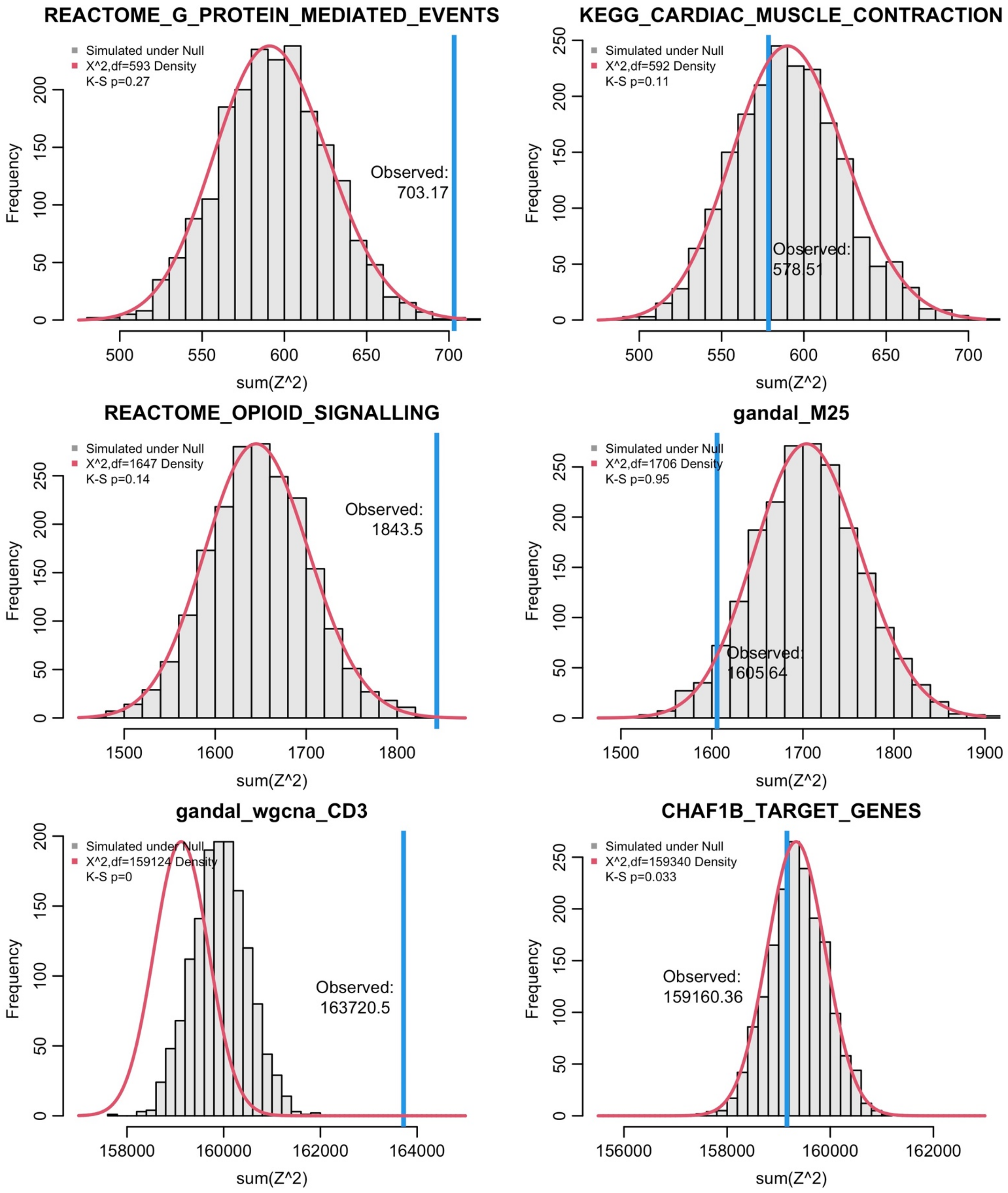


**Figure S33.** Distribution of summed, squared interaction Z-scores for 1000 simulated phenotypes under a null of no epistasis but including main effects for pAUDIT using cortex imputed expression for 6 different gene sets of varying size. Observed value shown by blue line, while the red line represents the $X^2$ density for the same df. These simulations show that for most gene sets, a standard $X^2$ test is appropriate, but can be anti-conservative for large gene sets, likely when there is a true signal (e.g., gandal_wgcna_CD3 set).

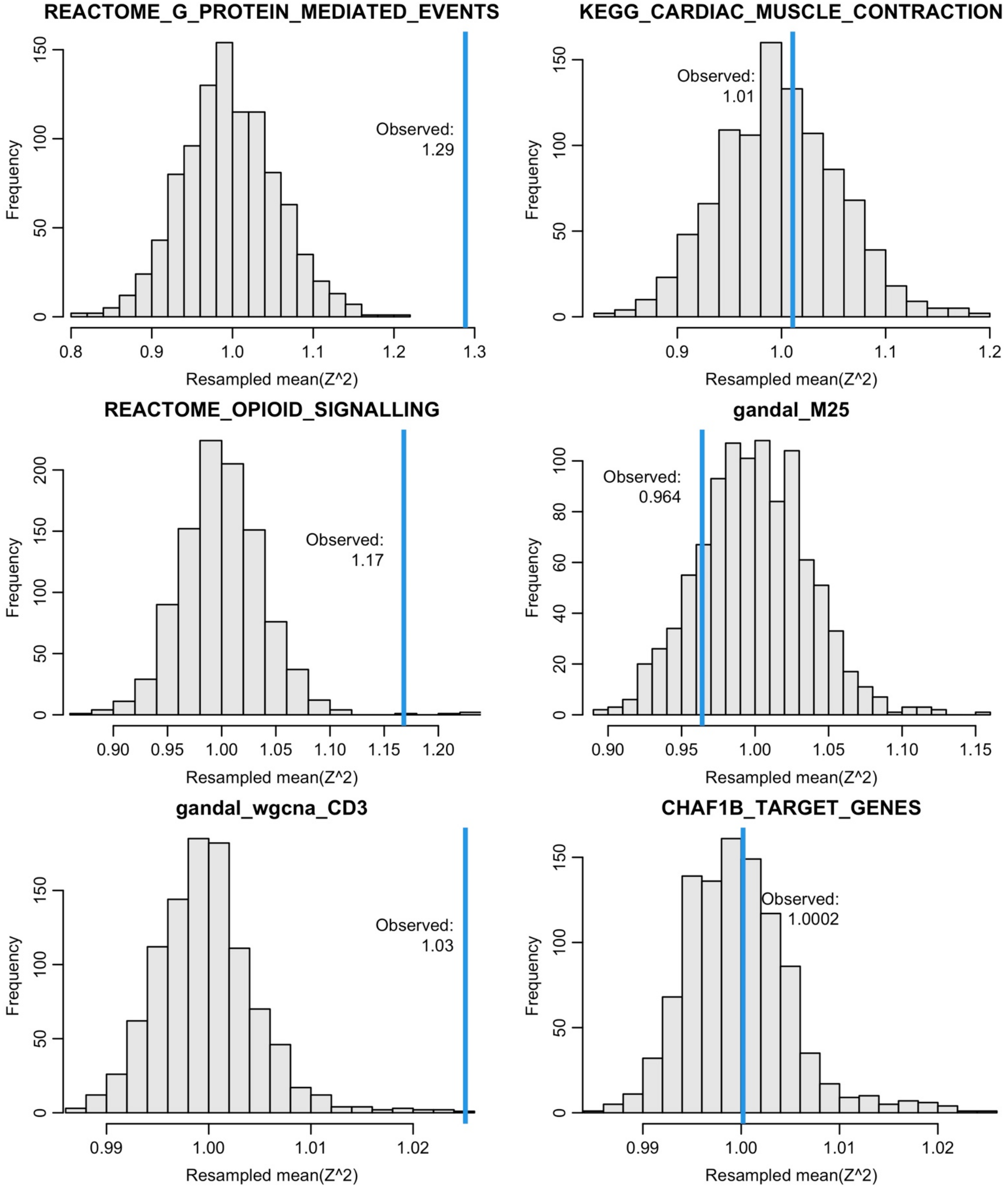


**Figure S34.** Distribution of mean squared interaction Z-scores for 1000 resampled gene sets for pAUDIT using cortex imputed expression for the same 6 different gene sets of varying size in Fig. S32. Observed value shown by blue line. These simulations show that for most gene sets, a random resampling approach recapitulates the results of a standard $X^2$ test.

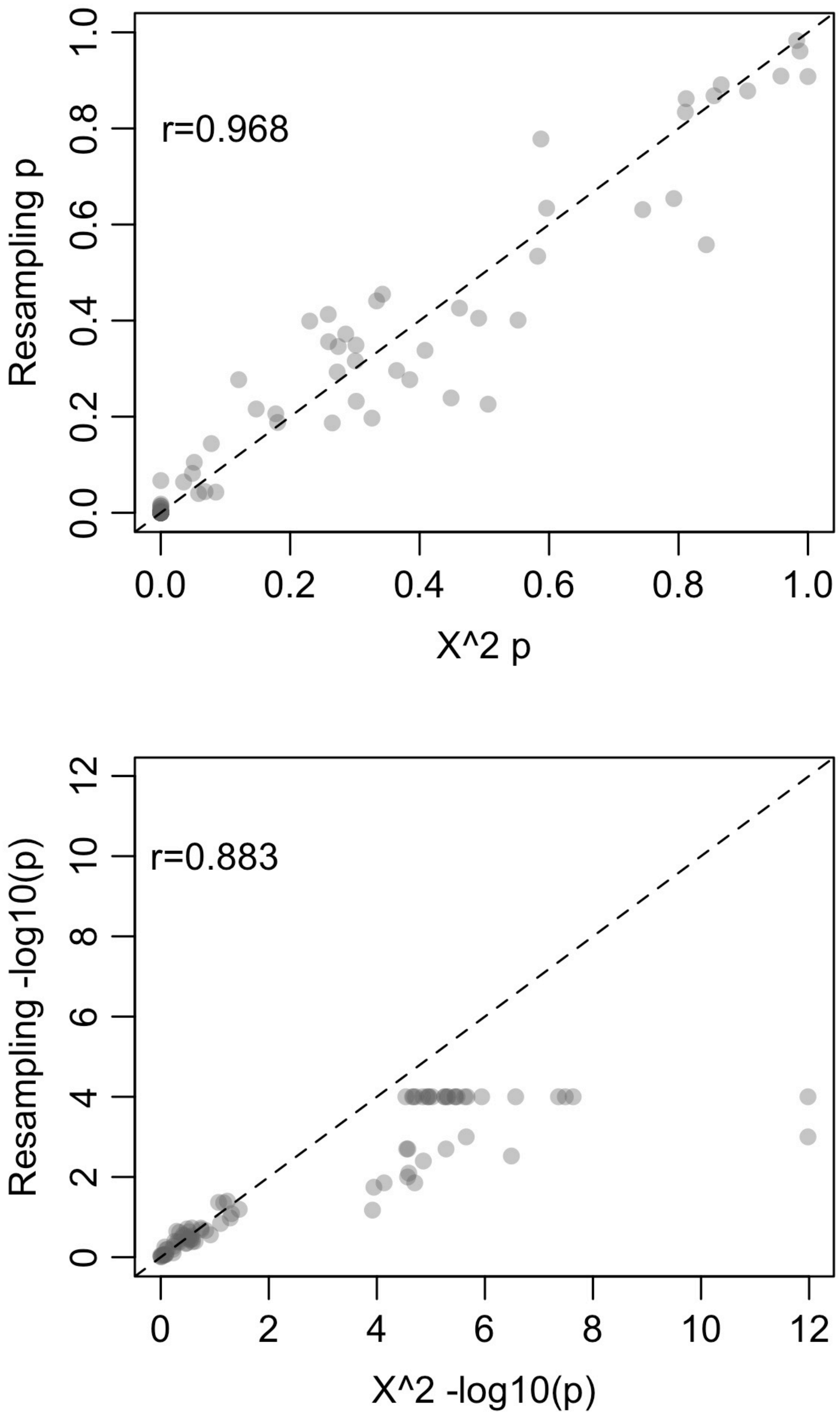


**Figure S35.** Comparison of p-values from a standard $X^2_m$ test vs. 1000 randomly resampled gene sets of the same size across a range of observed $X^2_m$ *p*-values. Note that in the bottom panel, all cases where the resampled *p*-value was <1/1000 (i.e., none of the resampled sets had larger mean $Z^2$ than the observed), -log10(*p*) was set to 4.
